# Genomic Population Structure of Atlantic surfclams: Cryptic Taxonomic Units and Population Connectivity

**DOI:** 10.64898/2026.07.27.740986

**Authors:** Hannah Hartung, Yuqing Chen, Matthew P. Hare

**Affiliations:** Department of Natural Resources and the Environment, Cornell University, Ithaca, NY 14853, USA

**Keywords:** mito-nuclear discordance, isolation by distance, ddRAD sequencing, hybridization, inshore-offshore parapatry, bivalve

## Abstract

The Atlantic surfclam, *Spisula solidissima solidissima* supports multimillion dollar harvests in the western North Atlantic, including Georges Bank and Mid-Atlantic Bight populations along the continental shelf. Surfclam populations in this part of the Exclusive Economic Zone (3-200 nm from shore) are managed as a single unit, but demographic connectivity has never been tested directly. This study analyzed >8,500 SNPs in 548 Atlantic surfclams sampled across the USA harvested range to infer population structure, genomic diversity, and gene flow patterns. Genomic analyses included eight sampled and 10 reference *Spisula solidissima similis* samples, a morphologically cryptic nominal subspecies previously known only from nearshore habitats. Novel morphologically cryptic population structure was identified in *S.s. solidissima.* One operational taxonomic unit (OTU-A) was found only in the inshore region of southern New England, south of Cape Cod. The other population unit, OTU-B, was found offshore and in Cape Cod Bay. The federal fishing grounds only had OTU-B clams. Populations within each OTU were connected by gene flow, justifying current fishery management practices. Nearshore state waters had mixed OTU stocks. Hybridization was analyzed between the two OTUs and between the two subspecies based on nuclear SNPs and asymmetrical mitochondrial DNA introgression. Finally, we demonstrated the utility of a novel SNP panel for diagnosing all three taxa and their hybrids.

## Introduction

In fisheries management, population structure can be an important variable within models used to predict sustainable levels of harvest, but it is often not formally incorporated into population dynamics models (Hilborn et al. 2003; Andersson et al. 2024). There is a risk that if spatial heterogeneities in vital rates or recruitment are not accounted for, biased population assessments can lead to overfishing (Kerr et al. 2017). Of course, knowledge of the population structure does not necessarily make a structured model the best choice in practice. For example, clinical population structure makes it ambiguous where to draw a distinction between stocks, and the data required for a robust structured model are not always possible to obtain. Nonetheless, awareness of possible spatial/temporal heterogeneity can lead to improved surveys and sensitivity testing of the impact of hypothesized structures on model results.

Detecting population structure is most challenging when evolutionary differentiation is morphologically cryptic, as is often the case among marine species (Knowlton 1993). Cryptic species are populations that are morphologically indistinguishable, yet genetically shown to be reproductively isolated. It stands to reason that morphological crypsis also obscures population structure below the species level (Cahill et al. 2023) to the detriment of fishery assessment accuracy (Hutchinson 2008). The concern is that morphologically cryptic divergence can be accompanied by behavioral, physiological, or life history trait divergence that also goes unrecognized (Bongaerts et al. 2021) yet is important for accurately modeling fishery impacts. Given the ever-increasing efficiency and power of genetic markers to reveal population structure (Allendorf et al. 2010; Benestan 2019; Carvalho 2025), and the confounding effects of range distribution shifts and associated community responses resulting from climate change (Marzloff et al. 2016; Pinsky et al. 2020), it is important that population structure assumptions in fisheries models are tested.

The Atlantic surfclam, *Spisula solidissima*, supports a commercial fishery that harvests 5- 15 thousand metric tons of clam meat each year in United States federal waters, with a $34 million dollar dockside value in 2023 (NOAA Fisheries, 2026). The federal fishery operates in the northeast and Mid-Atlantic Exclusive Economic Zone (EEZ), including Georges Bank, Nantucket Shoals, and the Mid-Atlantic Bight continental shelf (Hofmann et al., 2018; McCay et al., 2011; NEFSC, 2017). State fisheries in New York and New Jersey also occur within 3 nmi of the shore. Federal stock assessments of Atlantic surfclams have always treated all populations as a single assessment unit (Georges Bank to the mouth of Chesapeake) and applied fishery reference points based on analyses of a long time series for both fishery- and fishery-independent survey data. Differences in fishing mortality, vital rates, and their trends between the Georges Bank and Mid-Atlantic Bight surfclam populations instigated separate regional modeling with results combined for the single stock assessment (NEFSC 2017) and ultimately led to a combined single-stock model based on two stock areas (NEFSC 2022). The results of a coupled hydrodynamic and particle tracking model provided some support for the demographic independence of the Georges Bank relative to connectivity among Mid-Atlantic Bight populations (Zhang et al. 2015), raising the question of whether the fishery should be managed as two stocks.

*Spisula solidissima solidissima* has a complex life cycle (Fay et al. 1983) and is estimated to have single-generation larval dispersal distances averaging 100-200 km (Zhang et al. 2015). Under these circumstances, population health in one area can impact other distant populations in subsequent generations (Palumbi, 2003; Slatkin, 1993). Although high-fecundity species with planktonic larvae have long been assumed to have high connectivity across their range, genetic studies often find that this is not necessarily the case (Coscia et al., 2020; Maas et al., 2018; Underwood et al., 2020; Vendrami et al., 2021). Studies on recruitment in Atlantic surfclams have shown local effects on age structure and growth rate, indicating that there might be larval dispersal barriers on meso-geographic scales (Marzec et al., 2010; Wagner, 1987; Weinberg, 1999) or spatially heterogeneous post-settlement survival (Timbs et al. 2019). Understanding the patterns and mechanisms of realized connectivity among *S.s. solidissima* populations is important for sustainable fishery management.

Translating population genetic inferences into contemporary demographic insights of relevance to conservation and management requires the recognition that gene flow measures effective dispersal, requiring both movement and successful reproduction in the new location. For species with high early mortality, this approach disregards a large proportion of the early life cycle individuals that can frustrate direct demographic or close-kin approaches. In addition, indirect estimates of gene flow are temporally integrated over generations of demographic history (Slatkin 1987). Time-integrated estimates identify temporally dominant population connectivity features rather than being sensitive to process idiosyncrasies that might occur during an observational study using direct methods (Gagnaire et al. 2015; Helberg 2009).

*Spisula solidissima* comprises two morphologically cryptic subspecies: *Spisula solidissima solidissima* (Dillwyn, 1817), which includes commercially fished continental shelf populations, and nearshore *Spisula solidissima similis* (Say, 1822). *S.s. solidissima* is distributed from Cape Hatteras to the Gulf of Saint Lawrence, commonly inhabiting depths of 10-50 meters but also occurs nearshore (Abbott, 1974; Hofmann et al., 2018; Merrill & Ropes, 1969; Ropes, 1978). *S.s. similis*, sometimes referred to as the southern or Raveneli surfclam, has been reported to range from the Gulf of Mexico to Cape Hatteras (Walker & Heffernan, 1994) or Cape Cod (Johnson, 1915). More recently, it was documented in Long Island Sound and on the south coast of Massachusetts, where it had previously been misinterpreted as a diminutive inshore form of *S.s. solidissima* (Hare et al., 2010). Additionally, there is a growing interest in both taxa for aquaculture (Acquafredda 2026) and *S.s. solidissima* is a species of concern in the planning of Mid-Atlantic offshore wind farms (Bray et al., 2016; Munroe et al., 2022; Scheld et al., 2022; Stromp et al., 2023).

Using double-digest restriction-site associated DNA (ddRAD) sequencing, we examined the genetic relationships within and between *Spisula solidissima* populations to understand how many genetically differentiated groups exist in the currently fished area of the eastern U.S. and the extent to which they are connected by gene flow. Both nominal subspecies were analyzed to put *Spisula solidissima solidissima* variation in a broader context. In addition, to facilitate further study of morphologically cryptic *Spisula* taxa, a subset of highly differentiated markers was assembled into a genotyping by sequencing (GTseq) panel.

## Methods

### Sample Collection

A total of 922 surfclam samples were collected from suspected *S.s. solidissima* habitats. Samples were collected over a period of 23 years, from archived federal collections in 1999 to recent collections in 2019-2022 taken by dredge, diver, or rake (Fig 1; Table S.1). These included samples collected for this study (Long Island) and others provided by the New York State Department of Environmental Conservation (NYSDEC) survey in 2012 (NYSDEC 2013), Cape Cod Cooperative Extension, Barnstable County, Massachusetts for 2012 and 2017 samples, and the New England Fishery Science Center (NEFSC) federal surveys in 1999, 2019, and 2022 (Jacobson & Hennen, 2019; NEFSC, 2000). A subset of the 1999 federal survey samples previously had mitochondrial DNA (mtDNA) analyzed in Hare and Weinberg (2005). In addition, southern Long Island samples from the 2012 NYSDEC survey were analyzed using mtDNA and microsatellites by Fletcher and Hare (2024). Samples of *S.s. similis* collected near Washburn Island, Massachusetts, were used as reference samples in this study after confirming their taxon ID. Ten *S.s. similis* samples used in the ddRAD analysis came from Eel Pond (41.554, -70.546) and 28 used in the GTseq analysis were from western Washburn Island (41.552, -70.546).

**Figure 1:**
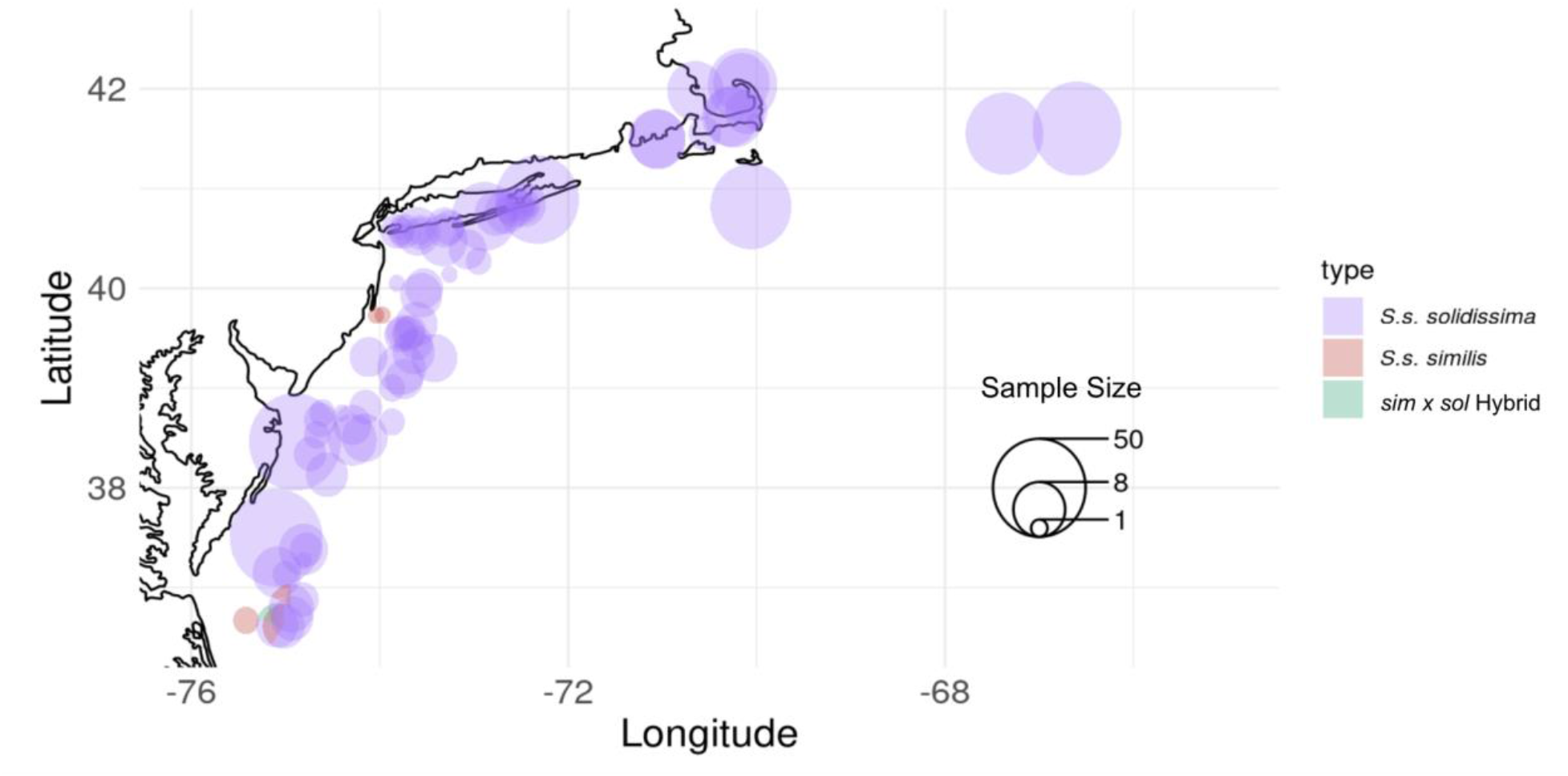
Sample sizes and collection locations for genomically analyzed S.s. solidissima and S.s. similis. Reference S.s. similis samples not included. “sim x sol Hybrid” refers to a single putative F1 hybrid between S.s. solidissima and S.s. similis near the southern terminus of sampling.

Shell dimensions were measured, and a subset of *S. s. solidissima* individuals collected by state and federal surveys were aged by counting annuli in the shells (Chute et al., 2016; Ivany et al., 2003; Jones et al., 1983). Tissue samples were stored in 90-100% ethanol until DNA extraction.

### DNA Extraction and ddRAD Sequencing

Of those sampled, a subset was chosen for sequencing, and 626 individuals provided good-quality DNA. Gill or adductor muscle tissue was extracted using the Qiagen DNeasy kit. Some adductor tissue collected during the 2019 sampling year did not yield good-quality DNA so re-extractions used the E.Z.N.A.® Mollusc DNA Kit from Omega Bio-Tek. Kit protocols were followed with the following adjustments: i) For DNeasy 96 well plate kits, we included RNase treatment; we centrifuged at 13,200 rpm after Buffer AW2 was added, incubated at 60 °C overnight, with final elution in 50-100 ul Tris-HCL warmed to 60 °C. ii) For Omega Bio-Tek kits, we made all of the same adjustments as for the DNeasy protocol, except incubation at 60 °C was only four hours and a third DNA wash was performed just before elution. Extracted DNA was stored in Tris at -20 °C.

We applied a DNA quality threshold of 260/230 ratio greater than 1, as measured using a Nanodrop ND-8000 spectrophotometer. We evaluated DNA degradation in a subset of samples from each year and site using 0.5% agarose gel electrophoresis. Samples without high-molecular-weight DNA were re-extracted using the E.Z.N.A.® Mollusc DNA Kit. Additionally, for library preparation, we required samples to have >700 ng total DNA, as measured by a Qubit 2.0 Fluorometer.

Reduced representation genomic sequencing was performed using double-digest restriction site-associated DNA sequencing (ddRAD-seq; Peterson et al., 2012). The sequenced samples were chosen to maximize geographic breadth and to include temporal comparisons.

Library preparation and DNA sequencing were performed by the University of Minnesota Genomics Center (UMGC) in three batches: i) pilot studies for each *Spisula* subspecies to establish *Pst*I and *Msp*I enzymes as a desirable combination and to estimate the sequencing effort needed; ii) 415 *S.s. solidissima* individuals from most locations included in this study, plus *S.s. similis* samples for another project, including the 10 reference samples analyzed here; and iii) 211 surfclams from the 2022 Mid-Atlantic Bight federal survey plus replicate samples to check for batch effects. Pooled ddRAD libraries were filtered to fragment lengths of 450-600 base pairs (bp) before Illumina sequencing with 150 bp paired-end reads using a NovaSeq platform. Reads were demultiplexed by UMGC.

### Sequence Quality Filtering, SNP and Haplotype Calling

Raw Illumina reads were assessed for quality using `FASTQC`. Individuals were eliminated from downstream analyses if they had sequence quality < 28 for more than 10% of bases or less than 1,000,000 total raw reads. Raw reads were trimmed i) first using the UMGC-recommended trimming script `gbstrim.pl`; ii) followed by `TRIMMOMATIC` version 0.39 (Bolger et al., 2014) to remove adapters and trim using a sliding window with default parameters (4:15), requiring read length ≥ 50 bp; iii) finally, using `SEQTK TRIMFQ`, eight bases were cut off at both ends of the R2 reads to remove UMGC sequence padding. All R1 and R2 reads were resynced into pairs.

Using `dDocent` version 2.8.13, the trimmed reads were then *de novo* assembled into a reference catalog of contigs following the recommended Assembly Tutorial pipeline (Puritz, 2016; Puritz et al., 2014). The *S.s. solidissima* reference catalog was created by excluding *S.s. similis* samples and using an equal number of individuals randomly drawn from each sampling site, except those collected in 1999 or 2022, totaling 66 individuals (O’Leary et al., 2018).

Trimmed reads for all samples (both nominal subspecies) were then mapped to the reference contig catalog. Alignment and SNP/indel calling were performed using `dDocent`’s default `bwa` and `FREEBAYES` parameters.

Two filtered LD-pruned SNP datasets were generated. One included all samples plus 10 reference *S.s similis* individuals to determine taxon assignment between *S.s. solidissima* and *S.s similis*. The other dataset included only *S.s. solidissima* individuals for most downstream analyses. The filtering criteria for variable nucleotide sites were identical for the two datasets (Table S.2) based on Puritz’s SNP Filtering Tutorial (Puritz, 2016) with some modified parameters. First, samples with more than 10% missing data across all called loci, or greater than 0.125 relatedness with another sample, were flagged and removed with `VCFtools` (Danecek et al., 2011; Manichaikul et al., 2010). To remove erroneous calls or loci with high levels of missing data, variants were removed if they had i) a quality value lower than 30, ii) minor allele frequency of less than 0.05 across all samples, iii) missing data in more than 15% of all individuals, iv) a mean coverage depth of less than 20, or v) more than two alleles. Loci were also removed if there was more than 10% missing data among samples from any of the eight regional populations (Table 1). Further filtering was performed with `vcflib` version 1.0.1 (Garrison et al., 2021) following the default recommendations of Puritz’s SNP Filtering Tutorial unless otherwise noted. Loci were excluded if they had heterozygous allele balance < 0.2 and > 0.8 or < 0.01 (O’Leary et al., 2018), reads from both strands, discrepancy in mapping quality or depth between reference and alternate alleles, or if only unpaired reads supported an alternate allele. The vcf was then re-coded to remove indels, and SNPs were filtered to remove population-specific deviations from Hardy-Weinberg equilibrium (*P* < 0.001).

**Table 1:** Sampling information, analytical region categories and number of individuals identified as S.s. solidissima OTU-A, OTU-B, or AxB hybrids (individuals with >10% admixture). Latitude and longitude coordinates are low precision to reflect aggregation of dispersed individual collections into analytical groups.

| Site | Year | Latitude | Longitude | Region | Number of Samples Sequenced | OTU A | OTU B | OTU Hybrid |
| --- | --- | --- | --- | --- | --- | --- | --- | --- |
| GBE1 | 2019 | 41.60 | -66.60 | George's Bank (GBE) | 44 | 0 | 43 | 0 |
| GBE2 | 1999 | 41.55 | -67.37 | GBE | 26 | 0 | 26 | 0 |
| CCB5 | 2012 | 42.04 | -70.15 | Cape Cod Bay (CCB) | 18 | 0 | 16 | 2 |
| CCB6 | 2017 | 42.06 | -70.16 | CCB | 11 | 0 | 8 | 2 |
| CCB4 | 2017 | 41.98 | -70.65 | CCB | 10 | 0 | 7 | 3 |
| CCB1 | 2017 | 41.77 | -70.11 | CCB | 5 | 0 | 5 | 0 |
| CCB2 | 2017 | 41.72 | -70.24 | CCB | 11 | 0 | 8 | 3 |
| CCB3 | 2017 | 41.72 | -70.28 | CCB | 10 | 0 | 9 | 1 |
| SCC1 | 2017 | 41.49 | -71.05 | Southern Cape Cod (SCC) | 20 | 20 | 0 | 0 |
| SCC2 | 2012 | 41.55 | -70.53 | SCC | 4 | 0 | 4 | 0 |
| NAT | 2020 | 40.82 | -70.06 | Nantucket Shoals (NAT) | 30 | 0 | 30 | 0 |
| SLI1 | 2019 | 40.89 | -72.33 | Southern Long Island (SLI) | 34 | 34 | 0 | 0 |
| SLI2 | 2012<br>2019<br>2022 | 40.82 | -72.46 | SLI | 12 | 8 | 4 | 0 |
| SLI3 | 2012 | 40.78 | -72.63 | SLI | 11 | 6 | 4 | 1 |
| SLI4 | 2012<br>2019 | 40.73 | -72.86 | SLI | 20 | 18 | 1 | 1 |
| SLI5 | 2012 | 40.61 | -73.29 | SLI | 7 | 6 | 1 | 0 |
| SLI6 | 2012 | 40.57 | -73.68 | SLI | 20 | 17 | 1 | 2 |
| NJ1 | 2022 | 40.39 | -73.33 | New Jersey (NJ) | 15 | 0 | 15 | 0 |
| NJ2 | 2022 | 39.53 | -73.68 | NJ | 55 | 0 | 55 | 0 |
| DEL1 | 2020 | 38.46 | -74.90 | Delmarva (DEL) | 49 | 0 | 49 | 0 |
| DEL2 | 1999 | 37.50 | -75.10 | DEL | 50 | 0 | 50 | 0 |
| DEL3 | 2022 | 38.63 | -74.29 | DEL | 40 | 0 | 40 | 0 |
| DEL4 | 2022 | 37.27 | -74.81 | DEL | 20 | 1 | 19 | 0 |
| DEL5 | 2022 | 36.69 | -74.94 | DEL | 20 | 0 | 20 | 0 |
| **Total** |  |  |  |  | **540** | **110** | **415** | **15** |

From the filtered but not LD-pruned *S.s. solidissima* dataset, we identified haplotypes using `rad_haplotyper.pl` with default parameters (Willis et al., 2017). This script parses through BAM files to link SNPs along paired reads into haplotypes and outputs them in a Genepop format. Haplotypes were used to estimate genetic diversity.

Finally, after the SNPs were filtered for quality, datasets with low linkage disequilibrium (LD) were prepared to reduce population structure artifacts caused by allelic correlations. The highest LD was expected to be within contigs, so each contig was filtered to a single SNP with the highest minor allele frequency using `vcftools` to both output allele frequencies per site and filter the list of selected SNPs. Further pruning of LD with `PLINK` version 1.07 (Purcell et al., 2007) removed one contig from each pair of contigs having r^2^ > 0.8.

### Mitochondrial DNA Marker

To study the discordance between mitochondrial and nuclear DNA (mtDNA and nDNA, respectively), we employed a simple restriction fragment length polymorphism (RFLP) marker based on the PCR amplification of mitochondrial Cytochrome Oxidase I (mtCOI) gene segments. The PCR primers and amplification protocol used were the same as those reported by Hare et al. (2010). Here, we used the restriction enzyme *Sph*I-HF (New England Biolabs) to digest 5 µL of the 600 bp PCR product in a reaction containing 3 units of enzyme, 3.7 ul of sterile water, and 1 ul of 10x rCutsmart buffer in a total volume of 10 µL. After digestion at 37 °C for 60 min, 10 µL was loaded in a 2% agarose gel and run for 60 min at 200v with visualization using GelRed (Biotium, Fremont, CA).

### Population Genetic Structure and Genetic Diversity

All population structure analyses used the LD-pruned SNP data. For a model-free initial exploration of population structure, we conducted principal component analysis (PCA) using `snpgdsPCA` from the `SNPRelate` R package in R version 4.2.1. To assign individuals to inferred population clusters and estimate individual admixture proportions, `STRUCTURE` version 2.3.4 was used (Falush et al., 2003; Pritchard et al., 2007) with the correlated allele frequency option `FREQSCORR 1`. Analyses were run assuming a number of clusters (K) from 1 to the number of sampling regions (8; Table 1), and the appropriate K was selected based on which K represented the greatest average change in likelihood (ΔK), averaged over 20 runs for each K > 1 (Evanno et al., 2005).

To quantify genetic differentiation between populations, the combined *S.s. solidissima* and *S.s. similis* LD-pruned dataset was used, and admixed samples were excluded based on a stringent admixture threshold (<1% assignment of genomic markers to alternate source populations based on STRUCTURE analysis). The pairwise Weir and Cockerham weighted *F*_ST_ (wc*F*_ST_) was estimated using vcftools between the two *S.s. solidissima* OTUs (operational taxonomic units; see Results) and *S.s similis*. The number of fixed SNP differences between each population pair was also estimated pairwise based on individual SNP wc*F*_ST_ = 1.0.

Several genetic diversity metrics were estimated from the ddRAD data. Using *S.s. solidissima* haplotype data, we estimated the expected heterozygosity (H_E_), haplotype richness, and inbreeding coefficient (F_IS_) within populations and their standard deviations using `arlequin` version 3.5.2 (Excoffier & Lischer, 2010). For haplotype richness, we randomly subsampled each population to include the same number of samples as the population with the smallest total sample size (N = 23) to eliminate sample size bias. Using filtered SNP data, the average number of SNPs per dDocent-assembled contig was used as an alternate measure of diversity, based on sample sizes of n=23. SNP counts were extracted from each contig in the subsampled individuals using `vcftools`. In addition, for comparison with the literature, `vcftools --hardy` and `awk` were used to calculate SNP-based expected heterozygosity and its standard deviation from the filtered SNP data. Diversity estimates by region were based on all *S.s. solidissima* samples, but estimates within the OTUs A and B had OTU hybrids excluded (>10% admixture). Statistical testing between OTUs A and B used a two-tailed t-test.

### Estimating Gene Flow

The LD-pruned *S.s. solidissima* SNP dataset was used to estimate relative levels of migration among populations within each OTU. For all gene flow analyses described below, we assessed OTUs A and B separately and excluded the Delmarva OTU-A individual and any individuals with >10% admixture between the OTUs based on the `STRUCTURE` analysis (Fig. 3).

We applied two methods that test for and map genetic differentiation relative to an equilibrium isolation by distance (IBD) model to summarize the locations where gene flow is relatively greater or less than the overall mean. We compared `SpaceMix` version 0.13 (Bradburd et al., 2016) estimates of genomic differentiation in geogenetic space to the Estimated Effective Migration Surfaces from `EEMS` (Petkova et al., 2016).

For `EEMS` analyses, half-missing genotype calls were set to missing data before computing the matrix of average pairwise differences using a `bed2diffs` script. Preliminary `EEMS` analyses were also run at the maximum recommended 1000 demes, and default 500 demes. Using five hundred demes gave noticeably reduced resolution, while 700 provided a very similar resolution to 1000 demes, and so was used for subsequent analyses. `EEMS` was implemented for OTU-A (OTU-B) with a burn-in of 1,000,000 (5,000,000) generations, and the MCMC iterated over a total of 10,000,000 (50,000,000) generations, recording every 1000th result. To check for convergence, we used plotting tools to examine posteriors using the `reemsplots2` R package (Petkova et al., 2016). The mean posterior effective migration rate was visualized using the `reemsplots2` R package.

For the `SpaceMix` analysis, small samples were grouped according to geographic proximity, as shown in Fig. 6a. We ran both the “target” and “source_and_target” models and in each case employed 10 fast runs iterated for 100,000 generations each to inform the program’s starting state. The target model explains the genetic covariances entirely by distorting the geographic sample position in ‘geogenetic’ space to reflect deviations from isolation by distance expectations. The “source_and_target” model also attempts to explain anomalously high covariance from afar in terms of recent admixture (`SpaceMix` documentation). From the fast run with highest ending posterior probability, we used the same parameter values to start a long run of 10,000,000 generations under default conditions. Sampling from this run was performed every 1,000 iterations for a total of 10,000 draws from the posterior. Run convergence was assessed in R, using the functions and plotting tools described in the `SpaceMix` vignette. Geogenetic plots were constructed from the parameter values averaged over the long run.

### GTseq SNP Panel

Our goal was to generate a tool that can efficiently and cost-effectively discriminate among *S.s. similis* and two *S.s. solidissima* OTUs. Genotyping by thousands (GTseq; Campbell et al. 2015) is a multiplex PCR method for generating amplicons from small regions of DNA containing target SNPs and preparing a library for high-throughput sequencing from which genotypes are called. The filtered ddRAD SNPs were rank ordered by F_ST_ among the three OTUs (*S.s. solidissima* OTU-A and OTU-B, *S.s. similis,*) and 370 of the most highly differentiated (F_ST_>0.85 between OTU-A and OTU-B) were chosen for panel development by GTseek LLC (http://gtseek.com/). GTseek employed a proprietary pipeline for multiplex PCR primer design from 163 target amplicon sequences, maintaining only one SNP per ddRAD contig and in the case of SNP pairs with high linkage disequilibrium. After two rounds of GTseq panel optimization on a set of 93 samples, including both OTUs and *S.s. similis*, 113 markers remained in the final panel reported here (Supplementary Table S.3). Zymo column clean-ups of the extracted DNA were needed to obtain more homogeneously positive results in terms of raw sequence output. Here, the discriminatory ability of these final GTseq markers was illustrated using PCA based on the `dudi.pca` command in R to cluster reference samples for the three taxa along with hybrids.

## Results

Among the acquired samples, 626 were selected and successfully sequenced. ddRAD sequencing yielded an average of 3.56 million trimmed paired reads per individual for 548 surfclam samples that passed filters (including sampled *S.s. similis* but not reference samples). The filtering removed 78 samples because of low sequence quality, relatedness, or contamination. Minimal batch effects were found by comparing PCA axis 1 and 2 concordance for 12 individuals sequenced across both runs (data not shown; Lou & Therkildsen, 2022). After removal of all *S.s. similis*, the assembly of 66 *S.s. solidissima* sequences (20 OTU-A and 46 OTU-B, see below) into a dDocent reference catalog resulted in 62,302 contigs. Once all trimmed reads (both nominal subspecies) were mapped to the reference contigs, variant detection identified 2,898,136 raw SNPs with an average depth of coverage per locus (for the mappable portions of the reference) of 77.6 reads. Quality filtering generated 22,683 SNPs, which were further LD-pruned to 8,628 SNPs. By mapping only *S.s. solidissima* reads, quality filtering generated 30,137 SNPs, and 9,634 remained after LD-pruning to produce the primary dataset used below unless stated otherwise. The filtering steps are summarized in Table S.2.

### Operational Taxonomic Units and Population Structure

In a preliminary PCA using LD-pruned SNPs from all samples, seven Mid-Atlantic Bight samples from the 2022 federal survey clustered with 10 reference *S.s. similis* and another individual was intermediate between *S.s. similis* and *S.s. solidissima* (Fig. S.1). Collections of *S.s. similis* were from two locations: 100 km offshore from the mouth of the Chesapeake at depths of 20-31 m, and 6 km off Barnegat Inlet, NJ, at a depth of 14-20 m (Fig. 1). After Removing *S.s. similis* and its hybrid to focus on *S.s. solidissima*, PCA calculated with LD-pruned SNPs showed clustering of samples into two distinct groups along the PC1 axis (Fig. 2). We refer to these genetically distinct *S.s. solidissima* groups as operational taxonomic units (OTUs), arbitrarily labeled A and B. Axis 1 in the PCA explained 15.3% of the overall genetic variance and included some intermediate individuals between the OTU-A and OTU-B clusters. Axis 2 explained only 0.35% of the genetic variance and spread-out populations according to geography; across latitude for OTU-B on the left of the plot and across southern New England longitudes for OTU-A on the right (Fig. 2). PC3 did not resolve further differentiation among individuals (figure not shown).

**Figure 2:**
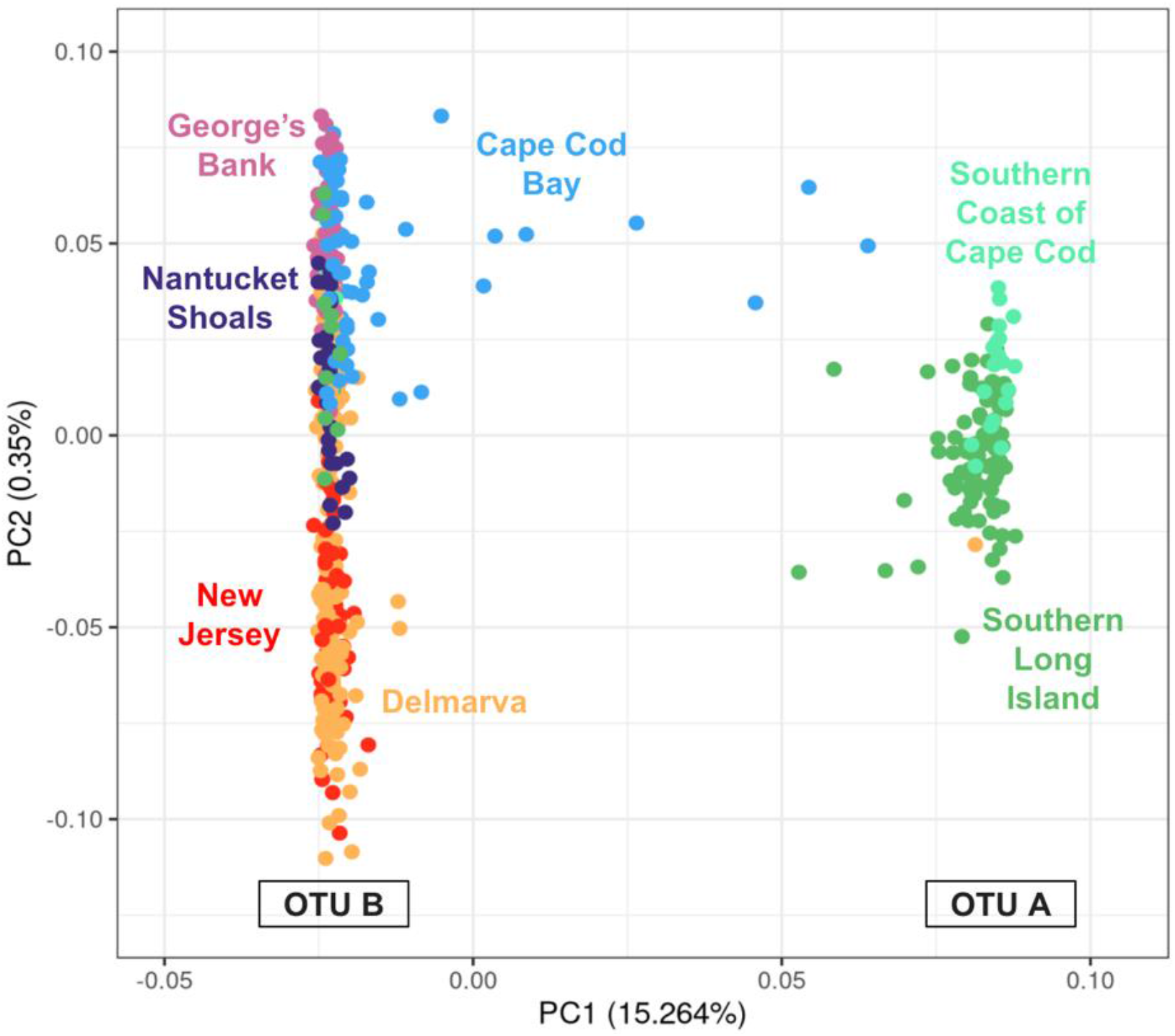
PCA plot summarizing allele frequency differentiation among S.s. solidissima individuals based on LD-pruned SNPs. PC axis percentages indicate the percentage of variation in the dataset summarized by each component axis. Genetically differentiated clusters of S.s. solidissima along PC axis 1 are assigned operational taxonomic unit (OTU) labels, with OTU-B clustered on the left of the PC1 axis and OTU-A on the right. Colors of symbols distinguish individuals from each of the seven regions listed in Table 1.

The most dramatic population structure within *S.s. solidissima* was the deep, phenotypically cryptic divergence of OTUs A and B despite co-occurrence in some regions. In the `STRUCTURE` admixture analysis performed with LD-pruned SNPs, the highest support was obtained for K = 2 source populations within *S.s. solidissima*, which separated individuals into OTUs A and B, with a few individuals showing mixed ancestry (Fig. 3). Counts of hybrid individuals in Table 1 are based on a criterion of >10% minor sources, and this criterion is used throughout, unless stated otherwise. Offshore sites (George’s Bank, New Jersey, Delmarva, and Nantucket Shoals) had largely unadmixed OTU-B individuals. Cape Cod Bay showed the highest proportion of OTU-AxB admixed individuals and included individuals that could be F1 hybrids between the OTUs (i.e., admixture proportions close to 50/50 in Fig. 3) and even F1 backcrossed to OTU-A, but this region had no pure OTU-A individuals observed. Southern Cape Cod and Southern Long Island contained individuals from both OTUs and a few hybrids in the latter region, but *S.s. solidissima* clams were predominantly pure OTU-A in these nearshore regions (Fig. 3). The `STRUCTURE` results for K=3+ resolved no additional geographical distinctions (Fig. S.2).

**Figure 3:**
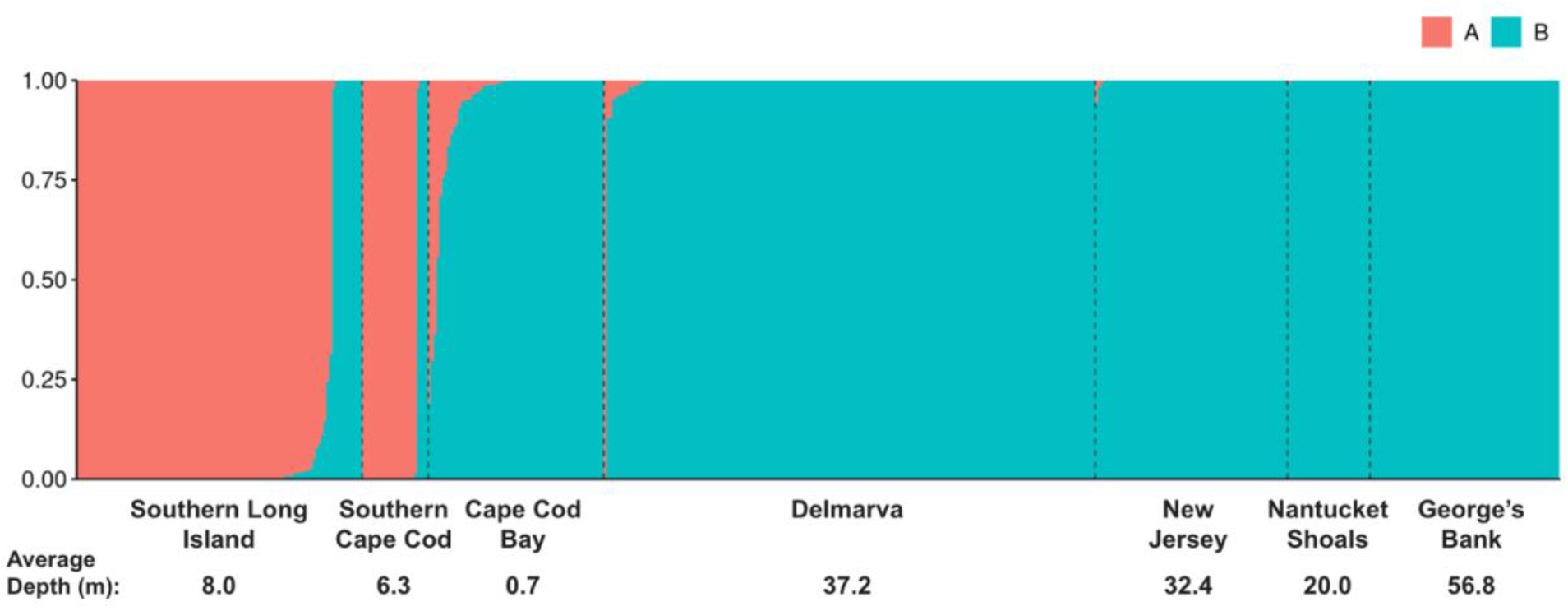
Bayesian clustering results from STRUCTURE applied to all S.s. solidissima samples with K=2 ancestral sources assumed. Each individual surfclam is represented by a vertical bar that is either blue (indicating OTU-B), orange (OTU-A), or a combination indicating admixture. Black vertical dashed lines separate collection regions detailed in Table 1. In populations containing admixed individuals, samples are ordered from fully OTU-B to increasing proportions of admixture or pure OTU-A. Average depth of collection for each population is listed across the bottom.

The *S.s. solidissima* samples that showed admixture in the STRUCTURE results (Fig. 3) were also spatially intermediate samples between the two clusters in the PCA along the PC1 axis (Fig. 2). Notably, fully OTU-A individuals only appeared in Southern Long Island and Southern Cape Cod, not offshore or in Cape Cod Bay. All Southern Long Island samples were collected within 1.5 km from shore (mostly much closer) at depths ranging from 2 to 16 m. If OTU-A occurs in deeper waters, it was not apparent from the federal survey samples in 2022, which were all OTU-B, including a single clam at 25 m depth just 4 km from Long Island, and 70 samples along NJ at depths ranging from 21 to 44 m that were no closer than 23 km from shore. One exception was a single OTU-A individual captured in the 2022 federal survey 74.4 km off Delmarva at a depth of 45 m (Fig. 4, difficult to see in Fig. 3). The total 2022 federal survey Delmarva sample consisted of 80 clams at an average depth of 37.2 m (19 – 55 m). The OTU-A geographic outlier was confirmed by repeated DNA extraction and genotyping.

**Figure 4:**
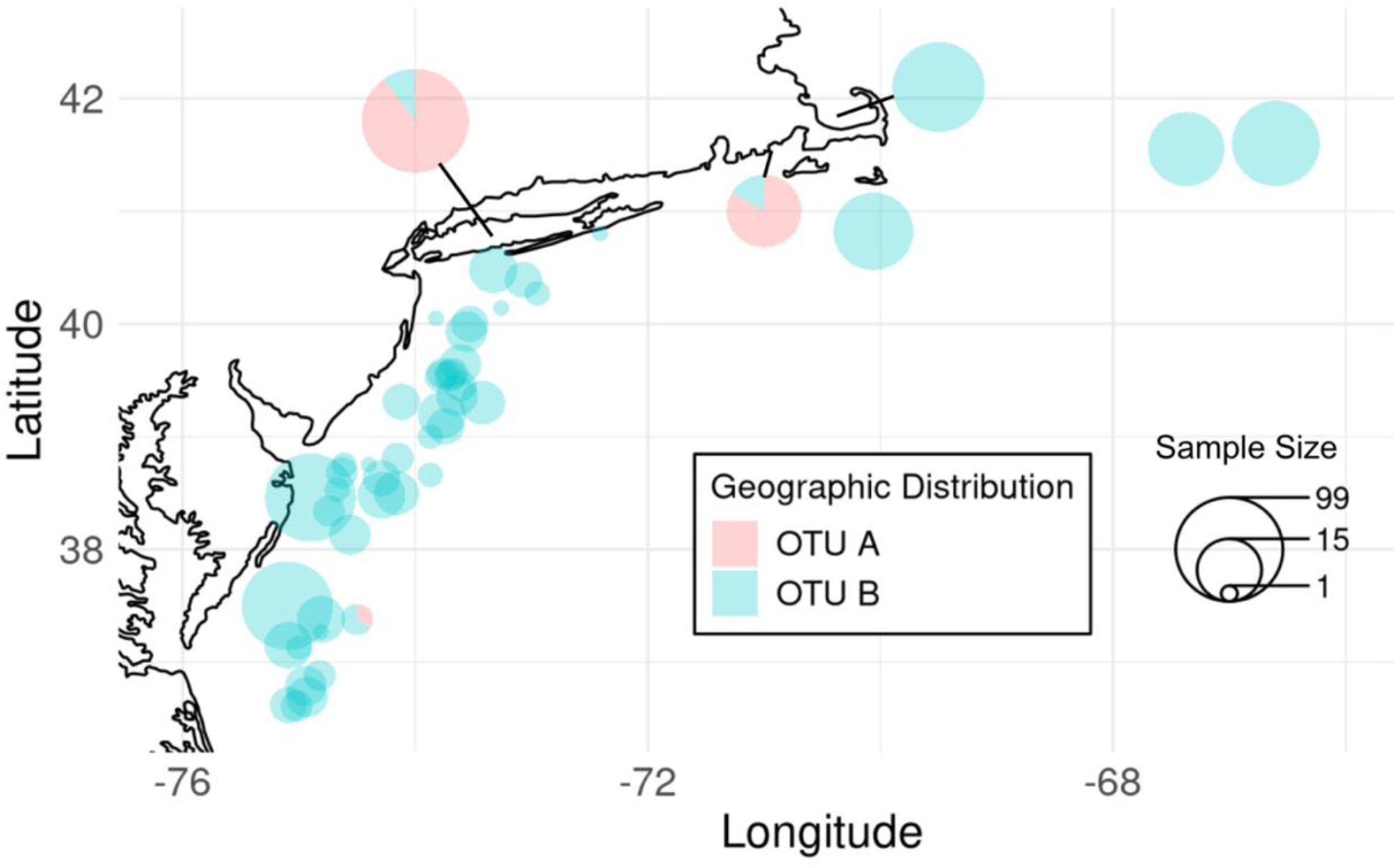
Distribution of non-admixed S.s. solidissima OTU-A and OTU-B individuals across all collections, using >10% admixture to define and exclude hybrids. Collection sites in Southern Long Island, Southern Cape Cod and Cape Cod Bay were consolidated into single pie charts to simplify this figure. Pie chart sizes are proportional to the number of individuals sampled at that site. Pie diagram colors indicate the proportion of individuals composed of OTU-A (pink) or OTU-B (blue) genotypes as determined by the K=2 STRUCTURE analysis (Fig. 3).

The *Sph*I RFLP distinguishes mtCOI variants that are at very different frequencies in OTUs A and B. Only one *Sph*I cut site occurs in the mtCOI most commonly found in OTU-A, making fragments of sizes 450 and 150 bp. In mtCOI from most OTU-B clams, three fragments were produced: 450, 90, and 60. The smaller fragments together appeared as an indistinct gel smear, in contrast to the OTU-A pattern. A subset of 316 surfclam samples were genotyped by *Sph*I RFLP. Focusing first on hybrids by applying a >1% minor admixture-source criterion to identify them based on nDNA, 43 hybrids also had mtCOI genotyped, with majority-OTU-A nDNA variation in 16 and majority-OTU-B in 27. Among the majority-OTU-A nDNA hybrids (all but 3 collected from Southern Long Island), only one (6.3%) had a mismatched OTU-B mtCOI haplotype, collected in Cape Cod Bay. For the majority-OTU-B nDNA hybrids, the mtDNA mismatch proportion was larger (18.5%). Most of these majority-OTU-B nDNA hybrids occurred in Cape Cod Bay, but a few occurred in southern Cape Cod, Southern Long Island, and offshore in the Delmarva federal survey samples.

Mitochondrial DNA also mismatched nDNA patterns in many ‘pure’ OTU-B clams away from nearshore mixed OTU zones. Among 273 surfclams that had <1% minor admixture-source (i.e., zero or nearly zero mixed ancestry in nDNA, referred to as ‘pure’) and were mtCOI- genotyped, 33 cases of mitonuclear discordance were found. Among the 183 OTU-B nDNA clams, 33 (18%) had the OTU-A mtCOI variant. In contrast, none of the 90 pure-OTU-A nDNA clams had the OTU-B mtCOI variant. Only seven (7.8%) of these mito-nuclear discordant clams were from nearshore populations where OTUs A and B most often co-occurred. The rest came from the two Georges Bank samples, Nantucket Shoals, and the two most inshore Delmarva samples (Table S.1).

The level of genetic differentiation between OTU-A and OTU-B samples (excluding those admixed with ≥1% minor sources, n=99 OTU-A and 385 OTU-B), as measured in nDNA with Weir and Cockerham weighted *F*ST, was 0.267 based on the LD-pruned data. Using the number of SNP loci with reciprocally fixed differences in the 3-taxon dataset to inform taxon diagnosability, there were 18 fixed differences between OTUs A and B (0.19% of LD-pruned SNP loci). A comparison of *S.s. similis* to OTU-A and OTU-B showed 62 and 4 fixed differences (0.72% and 0.05%, respectively).

### Genetic Diversity

Across the three diversity metrics reported in Table 2, OTU-B had significantly higher diversity than OTU-A. There was no apparent trend across geography within each OTU, but the diversity was higher in regions where hybrids were identified. Therefore, statistical testing was restricted to the OTU contrast with hybrids excluded (≥ 10% minor source admixture). The mean contig length was 253.6 bp with an overall average number of SNPs per contig of 3.01 ± 2.15 for all samples and a significant difference between OTUs (Table 2). When contigs were analyzed as haplotypes, there was an average of 3.24 ± 2.21 haplotypes segregating per locus in OTU-A compared with 3.54 ± 2.25 in OTU-B (range 1 – 33 for both). Expected heterozygosity, as calculated from haplotypes, was 0.35 ± 0.24 and 0.39 ± 0.22 in OTUs A and B (Table 2). For comparability to the literature, filtered SNP H_e_ in OTUs A and B was 0.20 ± 0.17 and 0.22 ± 0.14 respectively (not shown). The inbreeding coefficients calculated at the SNP level indicated a heterozygote deficit for all regions and both OTUs, with a slightly larger deficit in OTU-A. A higher-than-average F_IS_ was found in the Nantucket Shoals population sample.

**Table 2:**
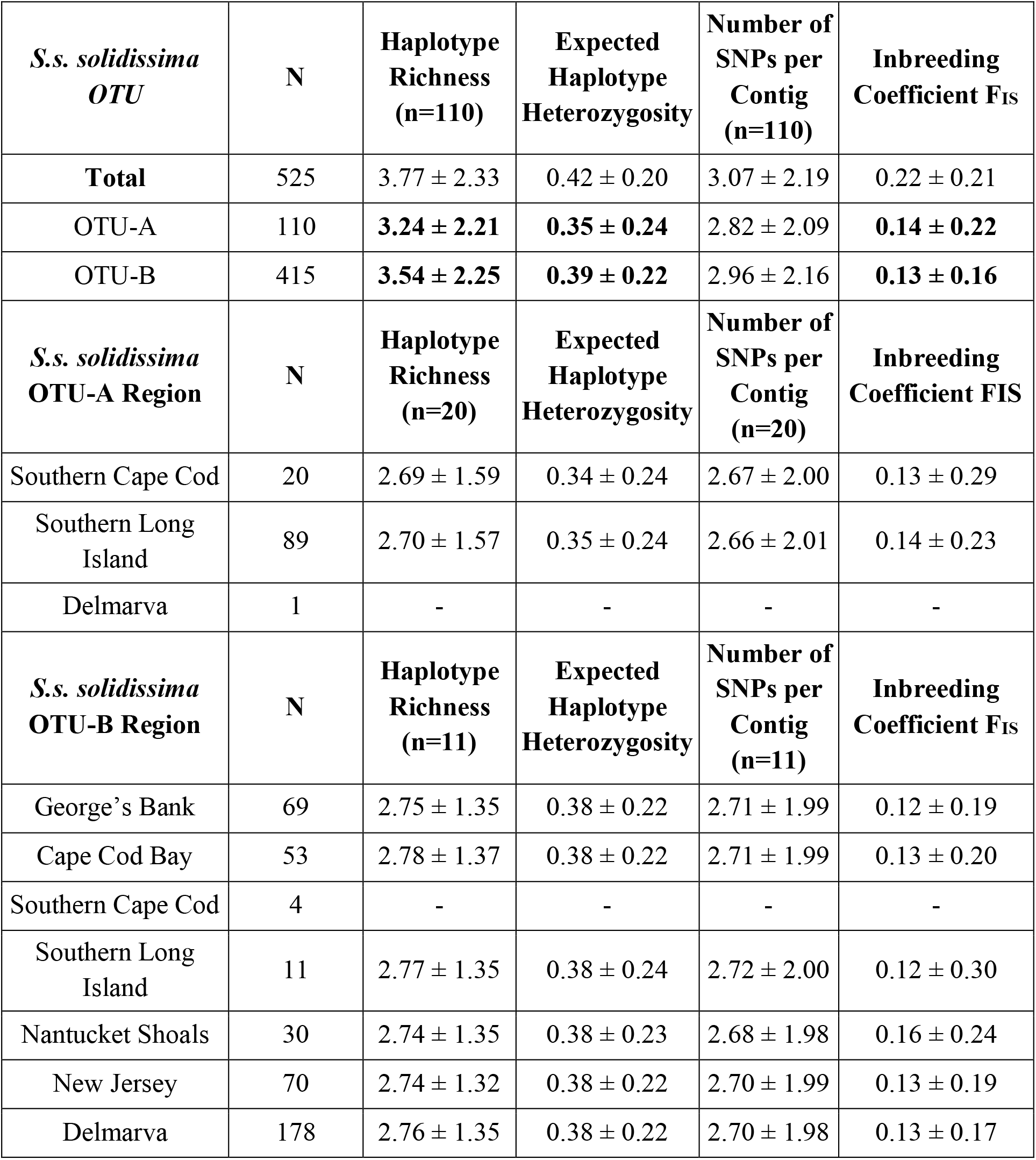
Summary of diversity statistics for sampled regions, all calculated including individuals admixed between OTUs A and B except for comparison of OTU-A and OTU-B at the top. Average values across all filtered loci are reported ± standard deviation. OTU refers to the operational taxonomic unit divisions found in S.s. solidissima (Fig. 2) and boldface font is used to denote statistical significance (P < 0.05) for 2-tailed t tests applied only between OTUs A and B. N is the number of samples sequenced from each region as defined in Table 1, but excludes 15 hybrids (admixture ≥ 10%) in the OTU and Total rows. Where column headers include parenthetical n individuals it refers to random subsample sizes based on the smallest sample size among compared sample groups.

| *S.s. solidissima*<br>OTU | N | Haplotype<br>Richness<br>(n=110) | Expected<br>Haplotype<br>Heterozygosity | Number of<br>SNPs per<br>Contig<br>(n=110) | Inbreeding<br>Coefficient $F_{IS}$ |
| --- | --- | --- | --- | --- | --- |
| **Total** | 525 | $3.77 \pm 2.33$ | $0.42 \pm 0.20$ | $3.07 \pm 2.19$ | $0.22 \pm 0.21$ |
| OTU-A | 110 | **$3.24 \pm 2.21$** | **$0.35 \pm 0.24$** | $2.82 \pm 2.09$ | **$0.14 \pm 0.22$** |
| OTU-B | 415 | **$3.54 \pm 2.25$** | **$0.39 \pm 0.22$** | $2.96 \pm 2.16$ | **$0.13 \pm 0.16$** |
| *S.s. solidissima*<br>OTU-A Region | N | Haplotype<br>Richness<br>(n=20) | Expected<br>Haplotype<br>Heterozygosity | Number of<br>SNPs per<br>Contig<br>(n=20) | Inbreeding<br>Coefficient $F_{IS}$ |
| Southern Cape Cod | 20 | $2.69 \pm 1.59$ | $0.34 \pm 0.24$ | $2.67 \pm 2.00$ | $0.13 \pm 0.29$ |
| Southern Long<br>Island | 89 | $2.70 \pm 1.57$ | $0.35 \pm 0.24$ | $2.66 \pm 2.01$ | $0.14 \pm 0.23$ |
| Delmarva | 1 | - | - | - | - |
| *S.s. solidissima*<br>OTU-B Region | N | Haplotype<br>Richness<br>(n=11) | Expected<br>Haplotype<br>Heterozygosity | Number of<br>SNPs per<br>Contig<br>(n=11) | Inbreeding<br>Coefficient $F_{IS}$ |
| George's Bank | 69 | $2.75 \pm 1.35$ | $0.38 \pm 0.22$ | $2.71 \pm 1.99$ | $0.12 \pm 0.19$ |
| Cape Cod Bay | 53 | $2.78 \pm 1.37$ | $0.38 \pm 0.22$ | $2.71 \pm 1.99$ | $0.13 \pm 0.20$ |
| Southern Cape Cod | 4 | - | - | - | - |
| Southern Long<br>Island | 11 | $2.77 \pm 1.35$ | $0.38 \pm 0.24$ | $2.72 \pm 2.00$ | $0.12 \pm 0.30$ |
| Nantucket Shoals | 30 | $2.74 \pm 1.35$ | $0.38 \pm 0.23$ | $2.68 \pm 1.98$ | $0.16 \pm 0.24$ |
| New Jersey | 70 | $2.74 \pm 1.32$ | $0.38 \pm 0.22$ | $2.70 \pm 1.99$ | $0.13 \pm 0.19$ |
| Delmarva | 178 | $2.76 \pm 1.35$ | $0.38 \pm 0.22$ | $2.70 \pm 1.98$ | $0.13 \pm 0.17$ |

### Estimates of Gene Flow

The spatial concordance to geography apparent within OTU clusters from the *S.s. solidissima* PCA (Fig. 2) is consistent with an isolation by distance (IBD) equilibrium between gene flow and genetic drift (Novembre and Stephens, 2008). To further assess population connectivity using quantitative model-based estimators of gene flow, we employed Estimated Effective Migration Surfaces (`EEMS`) and `SpaceMix`. `EEMS` and `SpaceMix` both use Bayesian MCMC to report posterior results relative to expectations from an IBD equilibrium model based on stepping stone migration or an approximation thereof and use maps to illustrate effective migration (gene flow) inferred based on the decay of genetic similarity across space. We assessed *S.s. solidissima* OTUs A and B separately, excluding one OTU-A individual off Delmarva and any individuals with > 10% admixture between the OTUs based on `STRUCTURE` analysis (Fig. 3).

With both the `EEMS` and `SpaceMix` runs, the log posterior stabilized long before the final MCMC iterations, indicating convergence (Fig. S.3). When long-distance admixture was included in the SpaceMix model it resulted in very low and similar admixture proportions in every population (0.002 – 0.006), so only results from the “target” model are presented. The greatest agreement between the results from the two programs for OTU-B was the inference of gene flow connectivity between samples in Cape Cod Bay and southern Cape Cod, as well as gene flow barriers between Cape Cod Bay and both Georges Bank and Nantucket Shoals (Fig. 5 and 6). `EEMS` inferred OTU-B gene flow constraints around CCB + SCC with log(m) approaching -2 (Fig. 5a). With `SpaceMix`, OTU-B from Cape Cod Bay clustered with southern Cape Cod in relative isolation (“geogenetic” separation) from Georges Bank and Nantucket Shoals (Fig. 6b). When differentiation is consistent with equilibrium IBD, the position of `SpaceMix` population labels in 2D geogenetic space reflects the geographic sample spacing (Bradburd et al. 2016). Higher than average gene flow brings sample labels (and their associated ellipses) closer together in geogenetic space, and gene flow constraints spread them apart relative to the sampling geography. The colored 95% posterior probability ellipses reflect uncertainty in the spatial relationships, and therefore, in the inferred relative levels of gene flow. In `the SpaceMix` geogenetic space, Georges Bank and Nantucket Shoals overlap with southern Long Island and are closer to the two most inshore Delmarva population samples (DEL1 and DEL2) than to Cape Cod Bay or southern Cape Cod, which is consistent with a gene flow barrier to the latter two locations. Southern Long Island populations are clustered and arranged orthogonally in geogenetic space to the NJ-Delmarva series, mimicking geography. However, the arrangement of southern Long Island populations is non-geographic at smaller scales; the far western SLI6 is shown furthest from geographically proximate NJ1 and instead overlaps both the Nantucket

**Figure 5:**
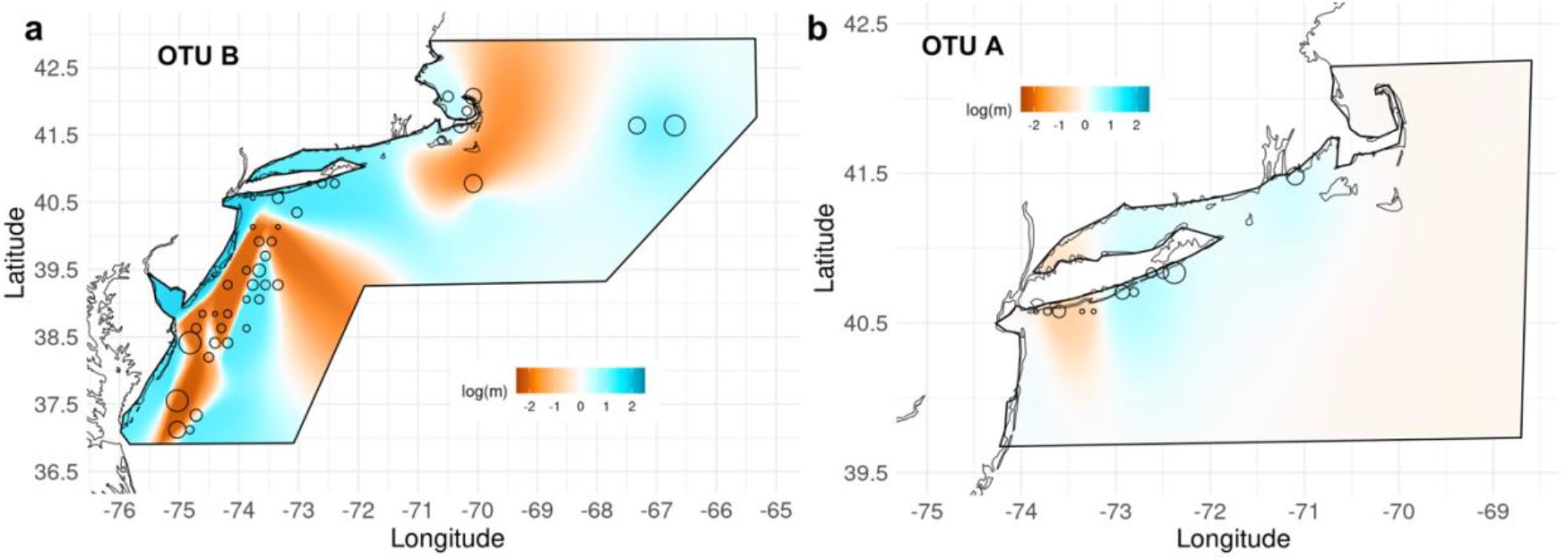
Estimates of population connectivity inferred with EEMS. Black polygons overlayed on a map of the study area describe the area in which the EEMS analysis was conducted for OTU-B (a) and OTU-A (b). Circles show sampling locations scaled by the log of the number of samples collected at each location. Estimated effective migration rates (m) on the log_10_ scale. Estimates of connectivity are illustrated relative to the null hypothesis of isolation by distance shown in white (log(m) = 0). Darker orange areas show inferred regions of reduced gene flow whereas darker blue areas represent areas of greater relative gene flow.

Shoals and Georges Bank. Note that few OTU-B surfclams were sampled along southern Long Island and the SLI2-SLI6 analytical groups had only 1-4 OTU-B samples represented in each. The New Jersey and Delmarva samples had geogenetic positions roughly concordant with geography, except that the offshore DEL3, DEL4, and DEL5 populations overlapped, indicating above-average gene flow among them. Additional `SpaceMix` indications of below-average gene flow include the broad spread of Cape Cod Bay populations in the geogenetic space, suggesting more internal gene flow constraints than the geographic scale of the bay implies. The status of Nantucket Shoals differs between the analyses. It is clustered with Georges Bank in the PCA (Fig. 2) and `SpaceMix` (Fig. 6b) but is shown by `EEMS` to be enveloped within a region of relatively low effective migration. Regions of low gene flow indicated by `EEMS` similarly encompassed population samples along New Jersey and Delmarva, perhaps suggesting a nearshore vs. offshore divide. For OTU-A, both `EEMS` and `SpaceMix` support IBD equilibrium conditions, except for inflated gene flow between eastern Long Island and Southern Cape Cod (Fig. 5b, 6c).

**Figure 6:**
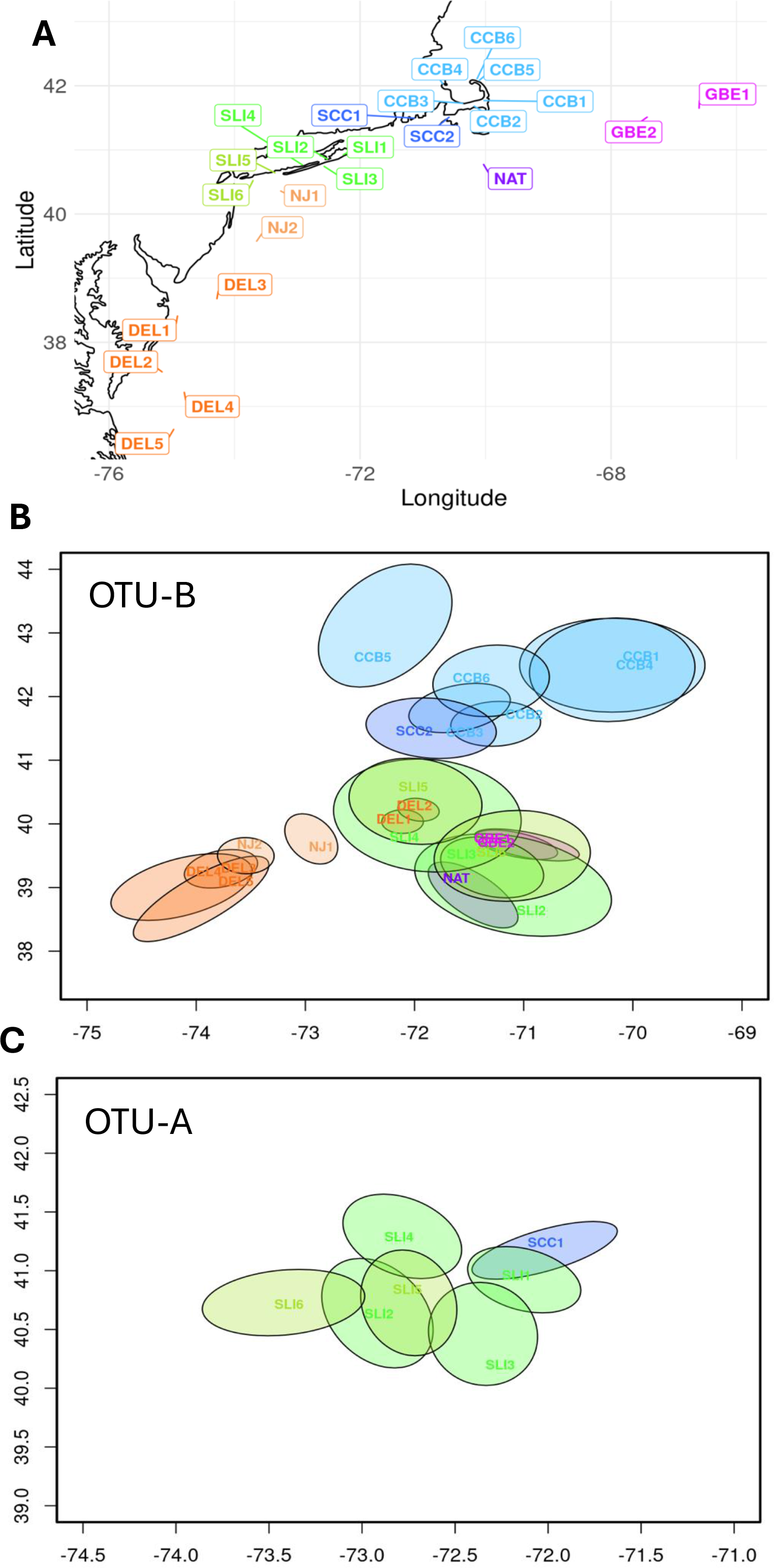
Estimates of population connectivity inferred with `SpaceMix`. **(a)** Map of OTU-A and OTU-B sample locations used as priors for `SpaceMix` runs. **(b-c)** `Spacemix` results for populations within OTU-B and OTU-A, respectively. Population labels are colored by sampling region as shown in (a), and shown in “geogenetic space” which uses geographic sampling location as priors to infer population relationships based on genetic covariances. The axes show an adjusted “latitude” and “longitude” in geogenetic space rather than actual cartesian coordinates. Population labels are point estimates for geogenetic location and similarly colored ellipses show 95% posterior probability ellipses.

### GTseq Panel

Optimization of the GTseq panel led to 113 SNP markers on different contigs (Table S.3). GTseq genotyping was performed on 19 *S.s. solidissima* OTU-A, 22 OTU-B, 29 *S.s. similis*, and 16 hybrids. The average sequence read count was 213,559 per individual, leading to an average percentage of loci genotyped per individual of 86.1% (range, 14.2 to 98.2). Sensitivity tests comparing duplicates across runs (with different levels of missing genotypes) suggested a conservative expectation that >100,000 reads provided >75% loci genotyped and accurate taxon ID for individuals. Principal Component Analysis clustered the GTseq data with 100% concordance to ddRAD results (Fig. S.4), although the hybrid admixture proportions were expected to differ based on GTseq vs. ddRAD. The average F_ST_ among the taxa based on GTseq was out of proportion with the ddRAD results (data not shown) because highly differentiated loci were chosen for the GTseq panel with an emphasis on distinguishing the two OTUs.

## Discussion

The commercially valuable federal fishery for *Spisula solidissima solidissima* in the U.S. EEZ is managed as a single stock unit. If it contains distinct segments with differences in productivity or natural mortality, single stock management that ignores these differences can potentially lead to overfishing of less productive populations (Kerr et al. 2017). For example, temporal DNA analyses of archived otoliths from Atlantic cod in West Greenland before and during population collapse showed that genetically distinct populations covaried in abundance with environmental changes and, given the lack of fishery management at the population scale, produced a vulnerability to high fishing pressure that led to population collapse (Bonanomi et al. 2015).

In this study, extensive geographic sampling and high-resolution genomic analyses resulted in two discoveries and several inferences with potential importance for surfclam taxonomic classification and fisheries management. Genetically distinct sister lineages exist in *S.s. solidissima* with OTU-B exclusively inhabiting the commercially fished continental shelf as well as Cape Cod Bay, whereas the novel OTU-A showed numerical dominance in shallower non-estuarine habitats of southern New England and the south shore of Long Island, NY. The genomic differentiation measured in this study is distributed across the genome and is presumed to reflect genetic drift and demographic history. Even if non-neutral SNPs contributing to local adaptation were sampled by our ddRAD method, their contribution to observed differentiation is expected to be minor because this reduced representation approach samples less than 1% of the genome (Lowry et al. 2017; Manel et al., 2016).

Patterns of gene flow within each OTU mostly agreed with expectations from isolation by distance equilibrium, providing strong support for moderate gene flow connectivity among all fished populations. The nominal *S.s. similis* is shown here to be a close relative of *S.s. solidissima* with patterns of genetic differentiation that further support the full species status previously advocated based on much less genetic evidence (Hare and Weinberg 2005). A small number of hybrid individuals were found at both divergence levels of this 3-taxon clade, including the first report of a natural hybrid between *S.s. solidissima* and *S.s. similis*. Further research, enabled by the marker panel described here, can help evaluate reproductive isolation and phenotypic differences between OTUs A and B.

The population genetic approaches used here to assess gene flow offer the strength that inferences are based on effective dispersal over recent evolutionary time, ignoring the vast majority of dispersal propagules that die before settlement or fail to reproduce after settlement (Lowe & Allendorf 2010; Neigel 1997; Slatkin 1987). The downside is that observed genetic patterns result from the temporal integration of population and environmental processes over many generations, potentially obscuring any contemporary change in processes. With climate change rapidly generating distributional and community composition changes for surfclams on the western North Atlantic continental shelf (Hofmann et al., 2018; Spencer et al. 2024; Weinberg 2005), there may be a growing mismatch between the within-taxon gene flow inferences made here, based on equilibrium theory and pertaining to at least dozens of past generations, relative to contemporary processes and their impact on population growth and resilience. Nonetheless, this study provides an important baseline that is only possible after clarifying cryptic evolutionary lineages. Below, we explore these findings and relate them to surfclam life history, comparative phylogeography, and their implications for fishery management.

### Biogeography and Taxon Boundaries in Spisula solidissima spp

Despite well documented *S.s. solidissima* phenotypic diversity across habitats (Cerrato & Keith, 1992; Davidson et al., 2003; Jones et al., 1978; Marzec et al., 2010), no previous study has detected genetic population structure within this taxon. This is not due to a subtle level of differentiation. The observed *F*_ST_ of 0.267 between OTUs A and B was larger than that typically found among populations in marine species with high gene flow. For example, eastern oysters had an average *F*_ST_ of 0.009 among Canadian populations (ddRAD SNPs: Bernatchez et al,. 2018), average *F*_ST_ = 0.031 among wild Atlantic populations (SNP array; Zhao et al. 2024), and average *F*_ST_ = 0.105 between the Atlantic and Gulf of Mexico (Puritz et al. 2022). North Atlantic Redfish (*Sebastes mentella*) had average *F*_ST_ = 0.053 among ‘shallow pelagic’, ‘deep pelagic’ and ‘demersal slope’ ecotypes (ddRAD neutral marker subset; Saha et al. 2021). Counter examples with high differentiation generally include a known biogeographic gene flow barrier, such as Cape Cod separating populations of sugar kelp (*Saccharina latissima*) with an average *F*_ST_ of 0.26 between the Gulf of Maine and southern New England (ddRAD SNPs; Mao et al. 2020).

The fixed difference statistic was used to emphasize the genetic diagnosability of all three *Spisula* taxa. The average *F*_ST_ of 0.267 between OTUs A and B included 18 fully fixed SNP differences, but these represented only 0.19% of LD-pruned SNPs in the 3-taxon dataset, one end of a broad distribution of SNP differentiation levels. When compared to *S.s. similis*, the much lower number of fixed differences in OTU-B relative to the contrast with OTU-A has two possible causes. Assuming that all these loci are alternately fixed via stochastic lineage sorting (genetic drift) from polymorphic ancestral populations, historical periods of relatively small effective population size in OTU-B may have amplified drift effects and increased the rate of fixation for ancestral polymorphisms relative to *S.s. similis*. Alternatively, gene flow between *S.s. similis* and OTU-B in the past could also generate a relatively smaller number of fixed differences.

While OTU-A individuals were found exclusively inshore (one exception in lower Delmarva region), OTU-B mostly occurred on the continental shelf. A few other examples of inshore/offshore genetic population structures exist in the North Atlantic: northern shrimp with months-long planktonic larval duration (Hansen et al. 2021; Skarðhamar et al. 2025), sea scallops in eastern Canada despite 40 d planktonic larval duration (Lehnert et al. 2019) and North Atlantic cod with spawning site fidelity (Bonanomi et al. 2016; Ruzzante et al. 1996; Sinclair-Waters et al. 2018). The OTU-B clams were abundant inshore only within Cape Cod Bay, where non-hybrid OTU-A was unobserved but likely present. In contrast, OTU-B was sparse inshore along southern Long Island (SLI), where they co-occurred with the numerically dominant OTU-A. This mixed SLI population is apparent in Fig. 2, where most SLI samples clustered on the right side of PCA axis 1 (OTU-A), but 11 non-hybrid SLI clams clustered with OTU-B.

Morphological crypsis between these OTUs is most apparent in this SLI overlap zone. For the 10 SLI OTU-B clams we obtained age/length data for, they averaged 6.1 yrs (2-11) and 121.1 mm shell length (62-167). The 50 non-hybrid OTU-A clams along southern Long Island, for which we have both length and age data averaged 10.6 yrs (2-25) and had 119.5 average shell length (58.7-159). At least from this small inshore sample, OTU-B individuals did not stand out in terms of age or length, nor were they in deeper nearshore waters on average. When all available size and age data for genotyped samples in this study were combined from all sampled regions, they suggested that growth rates might be distinct, but asymptotic sizes between OTUs A and B differed by only 10 mm (Table S1; Fig. S.5). It should be noted that our sampling methods did not focus on obtaining the full spectrum of sizes/ages across our sampling sites. In addition, size-at-age data for OTU-B, collected from diverse habitats, showed wide variance around the estimated von Bertalanffy curve. More direct size-at-age studies should be performed before drawing conclusions regarding OTUs A vs. B life history and plasticity.

As Cape Cod Bay is a significantly different environment than southern Long Island in terms of temperature, hydrodynamics, and vulnerability to hypoxia (Jiang & Zhao, 2006; Sankaranarayanan et al., 2014; Scully et al. 2022; Warner et al. 2014), it is impossible to know from the available data the relative role of historical biogeography, competition in sympatry, physiological tolerances, etc., generating the differences in nearshore frequency for OTUs A and B clams in these places. Cape Cod serves as a zoogeographic boundary for many species that have northern or southern range limits there (Pappalardo et al. 2015). Comparative zoogeographic analyses at Cape Cod identified shallow-depth species (< 20 m) as being more limited by Cape Cod as a northern boundary. In addition, relative to short planktonic larval duration (PLD) < 3d, longer PLD species tend to be more subject to Cape Cod as a northern range barrier, especially in shallower waters (Pappalardo et al. 2015). These process inferences are consistent with OTU-A experiencing Cape Cod as a northern range boundary, while the deeper-habitat OTU-B is not affected in this way. Thus, perhaps until recently, OTU-B might have had no competition from OTU-A in Cape Cod Bay.

Range distribution overlap zones make hybridization possible, but they are not the only locations where OTU AxB hybrid-ancestry individuals were found. We distinguish between recent hybrids in overlap zones (F1 and recent backcross types), diagnosed based on nDNA marker admixture within individuals, versus individuals with fully OTU-A or OTU-B nDNA (referred to as ‘pure’) and hybrid ancestry evidenced by mito-nuclear discordance. The latter occur both within and outside of OTU A and B overlap zones.

When hybrids are present but rare in geographic overlap zones between taxa (i.e., there is no hybrid swarm), it reinforces the inference that intrinsic or extrinsic reproductive barriers maintain the isolation of each lineage along independent evolutionary trajectories (Barton & Gale 1993; Jiggins and Mallet 2000). This study expanded the known area of overlap between *S.s. solidissima* and *S.s. similis* beyond southern New England (Fletcher and Hare 2024) and into the Mid-Atlantic Bight off of New Jersey (see also Wisner 2023). However, the only known natural hybrid between these nominal subspecies is the one reported here offshore from the Chesapeake Bay mouth. Previously, mitochondrial DNA variation in a single clam from the same lower Delmarva region was identified as characteristic of *S.s. similis*, but no nDNA variation was reported to evaluate hybrid ancestry (Wisner 2023). In contrast to the nominal subspecies, multiple hybrids were found in the overlap zones between OTUs A and B, albeit at low frequency. The low rate of hybridization among these three cryptic taxa suggests that reproductive isolating mechanisms operate prezygotically to limit hybrid crosses (e.g., different spawning seasons) or postzygotically to limit introgression (selection against hybrids).

Considering recent hybrids in Cape Cod Bay, where OTU-B is numerically dominant, 11 of 64 clams (17.2%) showed at least 10% nDNA admixture with OTU-A. A few had majority- OTU-A nDNA genomes, which is not possible without an F1 hybrid backcross to an OTU-A individual. Thus, it is likely that ‘pure’ OTU-A individuals exist in Cape Cod Bay even though they were not sampled in this study. It is possible that their immigration into the bay was by larvae passing through the Cape Cod Canal, or that aquaculture hatcheries supplying Cape Cod Bay lease farms are unwittingly using OTU-A broodstock from Southern Cape Cod to produce and sell seed clams. In contrast to Cape Cod Bay, OTU-A was numerically dominant in the southern Long Island overlap zone; 11 pure OTU-B individuals were sampled from a total of 104 *S.s. solidissima* clams. Only four (3.8%) OTU AxB hybrids were found (based on the 10% admixture criterion), all with majority-OTU-A nDNA genomes. Thus, the rate of recent hybridization is much lower along southern Long Island and in Buzzards Bay (SCC1; 0 hybrids out of n=20) than in Cape Cod Bay.

Based on hybridization patterns, we hypothesize that Cape Cod Bay is experiencing a dynamic recent invasion of OTU-A, whereas low hybridization rates in southern Long Island and southern New England may represent a parapatric equilibrium. This implies that OTUs A and B have extrinsic reproductive barriers that, while not complete, are currently maintaining the evolutionary integrity of each lineage in areas south of Cape Cod (Chan & Levin 2005), but less so in Cape Cod Bay. These might include spawn phenology, con-specific (OTU) sperm precedence, or environmentally contingent hybrid fitness.

Mitochondrial DNA is uniquely informative about older hybrid ancestry as well as indicating the directionality of the original hybrid cross. Assuming maternal transmission of mtDNA in surfclams (White et al. 2008), mtDNA reliably identifies the maternal source population that generated the original hybrid cross. In contrast, both parents contribute nDNA equally to the original F1 hybrid, but because subsequent backcrossing is most likely to the local numerically dominant taxon, the nDNA from the numerically minor taxon eventually gets diluted-out, leaving mtDNA on a ‘pure’ foreign nDNA background if not selected against (mtDNA is a nonrecombining cytoplasmic genome and experiences no such Mendelian dilution each generation). Thus, hybrid ancestry can be inferred using tests for mismatched (discordant) mtDNA against a ‘pure’ OTU nDNA background. The number of generations since original hybridization in a discordant individual’s pedigree is unknown, but the original (maternal) source population is identifiable.

For the *S.s. solidissima* OTUs, mito-nuclear discordance shows a broader geographic distribution than seen for recent hybrids. Here, a threshold of <1% minor admixture-source was applied to define a ‘pure’ OTU individual based on nDNA, with most such samples involving `STRUCTURE` Q = 0 for minor source. OTU-B mtDNA was not seen in any of the 90 ‘pure’ OTU-A clams (relative to 6.3% within recent hybrids), whereas 18% of ‘pure’ OTU-B clams had mtDNA from OTU-A (relative to 18.3% of recent hybrids). The vast majority of clams with OTU-A mtDNA paired with ‘pure’ OTU-B nDNA was seen at Georges Bank, Nantucket Shoals, and in the Mid-Atlantic Bight, with only 7.8% of such clams found nearshore in OTU overlap zones. This is a telling spatial asymmetry because the overlap zones are geographically restricted sources of recent hybridization. Assuming the current OTU overlap zones represent long term constraints on OTU-A distribution, all offshore mito-nuclear discordant clams have ancestry to hybrids in the nearshore overlap zones. In other words, offshore mito-nuclear discordance is evidence of nearshore to offshore gene flow (over an indeterminant number of generations). A similar frequency of mito-nuclear discordance occurred in 1999 vs. 2020 replicate samples for DEL1+2, and 1999 vs. 2019 for GBE1+2, indicating that hybrid ancestry is a persistent minor component of these continental shelf populations. These spatial gene flow insights are most interesting in the Mid-Atlantic Bight where mito-nuclear discordance was restricted to clams in the relatively nearshore DEL1 and DEL2 continental shelf samples, not in 2022 DEL3, 4 or 5 samples from further offshore. Furthermore, the presence of OTU-A mtDNA on the nearshore continental shelf of Delmarva implies that there are (or were) nearshore OTU-A populations in this area that did not get sampled in this study.

Most cases of animal mtDNA introgression involve foreign mtDNA, i.e. mito-nuclear discordance, at moderate to high frequencies (>50%) (Toews and Brelsford 2012). In the cited meta-analysis, almost all cases with introgressed mtDNA at low frequency (<50%) involved a restricted geographic extent of discordance (<50% of the taxon range where introgression could have spread; Toews and Brelsford 2012), as in *S.s. solidissima*. In this context, mito-nuclear discordance can signal both past hybridization and highlight patterns of historical population connectivity (Toews and Brelsford 2012). It is possible that stepping-stone gene flow across generations and across habitats could be accompanied by changes in selection pressures on mito- nuclear discordant lineages. However, the temporally constant frequency of OTU-A mtDNA discordance offshore on the continental shelf argues that this is not the case, except perhaps across the Delmarva continental shelf. The absence of OTU-A mtDNA in Delmarva populations further offshore than DEL1 and DEL2 could be due to gene flow connectivity barriers (see Zhang et al. 2016 for possible mechanisms) or spatial differences in the fitness of mito-nuclear discordant genotypes.

The genealogical and evolutionary behavior of mtDNA in this system may be informative about gene flow connectivity and/or adaptive introgression, but only if the simpler alternative hypothesis of incomplete lineage sorting (ILS) of ancestral polymorphism is rejected. If two types of mtDNA existed in the common ancestor of OTUs A and B, then it is possible that ILS, a genetic drift process, led to near alternate fixation of these mtDNA types in OTU-A and B populations. Under the ILS hypothesis, apparent mito-nuclear discordance is merely lingering ancestral polymorphism (Sloan et al. 2017). Given hybridization in a contact zone in addition to ILS, more of the mito-nuclear discordant pattern might be expected in the contact zones. Outside those zones, the ILS hypothesis does not predict spatial heterogeneity in the distribution of mito- nuclear discordance (Funk & Omland 2003) as observed here in the Mid-Atlantic Bight. Thus, the geography of the surfclam mito-nuclear discordance in OTU-B populations is inconsistent with the ILS hypothesis, and therefore gene flow constraints and/or selection provide better explanations for the data.

In taxon overlap zones where hybridization occurs, strong discrepancies in abundance, such as documented here, create likely mating asymmetries (Burgess et al. 2005; Qvarnström et al. 2022). Broadcast spawning invertebrates such as surfclams that release gametes into the water have mate-choice mechanisms that, as a function of molecular gametic interactions after spawning, confer within-species and between-species choosiness of eggs for certain sperm (Evans et al. 2020; Geyer and Palumbi 2005). Coupled with Allee effects, weaknesses in this choosiness gives females of the less abundant taxon a higher probability of hybridizing than the reverse heterospecific pairing (Qvarnström et al. 2022; Wirtz 1999). In southern Long Island, OTU-B is far less abundant than OTU-A, and as a function of this rarity, the majority of F1 hybrids are expected to inherit mtDNA from an OTU-B maternal parent. Subsequent backcrossing is most likely with the numerically dominant taxon (OTU-A), further introgressing the OTU-B mtDNA into the OTU-A population. Thus, in this region, OTU-B mtDNA is more likely to introgress into the OTU-A population than vice versa based on the first principles of gamete fertilization dynamics and with all else being equal. However, this directionality was rarely observed compared with the opposite (OTU-A dam) in recent hybrids. In Cape Cod Bay, where OTU-A clams are relatively rare, the observed mtDNA introgression asymmetry is again opposite that expected; only 18.5% of nDNA hybrids had OTU-A mtDNA. Thus, in both locations mito-nuclear discordant hybrids produced by the less abundant female were under- represented compared with expectations under the “sexual selection hypothesis” (Wirtz 1999).

Similar area-specific patterns of unidirectional mtDNA introgression have been reported in other species where relative parental population abundance failed to predict outcomes (Dowling et al. 1989). Other factors that can contribute to the direction and degree of introgression asymmetry include selection against mito-nuclear combinations (with potentially environment-specific fitnesses), taxon differences in heterospecific fertilization error rates, and genetic drift in small hybrid populations (Qvarnström et al. 2022). Further research is needed to determine the relative importance of these factors in the surfclam hybrid populations. Whatever causes the mito-nuclear discordance asymmetries in the nearshore OTU overlap zone hybrids, similar asymmetries were found in the ‘pure’ OTU-A or OTU-B clams, i.e. those indicating historical hybridization. In other words, mito-nuclear discordance asymmetry patterns seem to be maintained over time in hybrid lineages.

In terms of biodiversity conservation, there is value in recognizing, taxonomically naming, and further studying the two OTUs, especially given the possibly expanded opportunities for aquaculture utilization. Even though there is fixed SNP diagnosability for all three taxa, we recommend recognition of the OTUs as Evolutionary Significant Units (ESU; Waples 1995; Coates et al. 2018) within *S.s. solidissima* until quantitative phenotypic comparisons can be made and habitat affinities are better understood. In addition, a more complete population-phylogenetic analysis is needed for the trio of taxa with the benefit of additional *S.s. similis* sampling (work in progress). Note that the introgression of mtDNA variants across *Spisula* taxa, as demonstrated here, means that mtDNA analysis alone is not conclusive regarding the evolutionary lineage a clam belongs to, underscoring the need for an efficient nDNA tool for taxon diagnosis.

### Within-taxon Diversity and Gene Flow

Both *S. solidissima* OTUs were found to have high within-population genetic diversity, based on several metrics. For example, the 0.20 filtered-SNP expected heterozygosity in OTU-B was similar to the average reported for wild eastern oyster populations by Zhao et al. (2024; 0.27) and for hard clams by Ropp et al. (2023; 0.172). It is unsurprising that OTU-B had a slightly higher genetic diversity than OTU-A, given the more spatially extensive continental shelf habitat inhabited by OTU-B. Neither OTU showed a geographic diversity trend.

The continental shelf populations of surfclams subject to the highest fishing pressure were all found to be OTU-B individuals with moderate gene flow connections. The Cape Cod Bay population of OTU-B showed reduced gene flow with shelf OTU-B populations (below IBD expectations) based on both `EEMS` and `SpaceMix`. Gene flow was assessed after removing OTU- AxB hybrids determined from nDNA; therefore, the recent OTU-A ancestry in Cape Cod Bay did not generate the appearance of gene flow isolation for OTU-B. The similar mito-nuclear discordance pattern and rate in Cape Cod Bay and offshore populations is consistent with non-zero gene flow between these populations in agreement with the `EEMS` and `SpaceMix` results. As discussed above, OTU-A ancestry in Cape Cod Bay is hypothesized to be a result of recent anthropogenic colonization from populations that formerly had their northern limit at Cape Cod.

However, not all gene flow results led to clear inferences. For example, `SpaceMix` and `EEMS` provided conflicting results regarding the connectivity of Nantucket Shoals to other offshore sites. While `SpaceMix` clustered all offshore sites in a geogenetic arc concordant with model-predicted stepping-stone larval dispersal trajectories (Zhang et al. 2015), the `EEMS` result suggests that Nantucket is relatively isolated. This conflicting result could have been caused by our nonuniform geographic sampling, as `EEMS` operates best with more uniform sampling and a greater number of sampled sites (Petkova et al., 2016). The New Jersey and Delmarva EEMS result for OTU-B showing a longitudinal gene flow barrier also is difficult to interpret and reconcile with the `SpaceMix` result.

The biophysical model of Zhang et al. (2015) focused on surfclam larval dispersal on continental shelf habitats, which are most relevant for OTU-B. Releasing larval “particles” throughout the surfclam April–October reproductive season generated predictions for a dominant stepping stone pattern of dispersal to the west along southern New England and then further southwest along the shelf. Surfclam larval dispersal and settlement was inferred to be a combination of nearly unidirectional movement to the adjacent ‘downstream’ region and substantial self-recruitment within each region. The Georges Bank region had no immigrants, but had high (80.7%) larval retention (Zhang et al. 2015). Their model included many biological parameters, including larval behavior, growth, physiological tolerances, and mortality resulting from slow growth or settlement in poor quality habitat. With all these biological parameters ‘turned on’, settlement proportions at the end of the 35 d planktonic larval duration (i.e., connectivity) were 2 – 20 times higher than proportions incorporating growth/settlement mortality. Thus, simulation results about larval connectivity may have been sensitive to mortality parameterization. For example, connectivity between Georges Bank, Nantucket Shoals, and Long Island was only observed based on transport alone (i.e., with mortality shut off), suggesting that constraints on successful settlement can isolate downstream regions from Georges Bank and Nantucket Shoals as recruitment sources. In contrast, the OTU-B population genetic inferences reported here, based on `SpaceMix`, provide no indication that Georges Bank and/or Nantucket Shoals have low gene flow connection with southern Long Island populations or the Mid-Atlantic Bight. These population genetic inferences describe the gene flow rates most likely to produce the observed population structure under an equilibrium isolation by distance model. The population genetic inferences reflect an integration of all the biological parameters incorporated into the biophysical model, but with a focus on gene flow instead of dispersal. Relatively low average connectivity (possibly too small for the biophysical model methods to capture) can have large evolutionary and significant demographic homogenizing effects (Legrand et al. 2022; Lowe & Allendorf 2010). Interestingly, the greatest geogenetic connection of Nantucket Shoals and Georges Bank populations to the Mid-Atlantic Bight involved more inshore DEL1 and DEL2 population samples from 1999 and 2020, not the northernmost NJ populations or outer shelf Delmarva populations. The biophysical model of Zhang et al. (2015) produced seasonally variable onshore transport along NJ and Delmarva that may be related to this finding.

Across both the `SpaceMix` and `EEMS` results for OTU A, the gene flow inferred between southern Long Island and southern Cape Cod approximates what would be expected under equilibrium IBD, with slight variation in connectivity patterns. From `EEMS`, the value of log(m) is near zero between SCC1 and Long Island, reflecting the IBD average, and slightly higher than average gene flow is inferred among SLI1-5 (Fig. 5). The westernmost SLI6 is painted orange by `EEMS`, indicating below-average gene flow with southern New England. The `SpaceMix` result for OTU-A sets SLI6 slightly apart to the west and has an overlapping cluster of the other southern Long Island samples. The geogenetic position of SCC1 is relatively closer to southern Long Island than the geographic reality, indicating higher than average IBD gene flow (compare Fig. 6a and c).

### Implications for Fisheries Management and Aquaculture

The most recent stock assessments for U.S. *S.s. solidissima* populations, treating them as a single spatial management unit as established under the 1977 Fisheries Management Plan, have concluded that they are not overfished and that overfishing does not occur (NEFSC 2022). The results of this study provide support for continued management as a single unit, given the gene flow connectivity inferred among shelf populations of OTU-B that are subject to the federal fishery in the EEZ. In fact, the gene flow results here suggest that over the long term, there may be less larval movement asymmetry than indicated by biophysical modeling (Zhang et al. 2015). There was no ‘downstream’ increase in OTU-B genetic diversity, as expected if the gene flow was unidirectional (Xuereb et al. 2018). Mixed stock complications appear to be present nearshore in state waters along southern Long Island and New England as a result of slight OTU- A and OTU-B range overlap and to a small degree from *S.s. similis* along New Jersey and near the southern edge of the OTU-B range.

The discovery of the cryptic taxon OTU-A complicates the prediction of climate change effects on surfclam populations. Instead of trying to model the physiological and evolutionary responses of one population, the potentially divergent physiologies and population interactions of the two taxa need to be considered along with their hybridization potential. Continental shelf OTU-B populations have responded to ocean warming events by contracting from their southern and inshore ranges in the south and moving into deeper and cooler waters (Weinberg, 2005, NEFSC 2013, 2017, Narváez et al. 2015, Timbs et al. 2019). The range boundary between OTUs A and B is uncertain, but at least near Long Island, it appears to occur along the inner shelf where influences of the cold pool have been minimal (Horwitz et al. 2023) and therefore, the risks from bottom water warm temperature anomalies are high. To better understand habitat use patterns, it is important for state and federal surfclam surveys to generate a baseline understanding of the relative densities of OTUs A and B at various depths, in addition to research characterizing physiological and life history differences.

The GTseq SNP panel described here is a valuable identification assay for population monitoring and physiological studies. Although faster and cheaper tools using fewer loci can be developed, several considerations argue for the value of testing with more than a few loci for certain goals. No single nDNA marker is known to definitively diagnose OTU-A, OTU-B, and *S.s. similis*. The *Sph*I RFLP in mitochondrial COI employed here easily distinguishes *S.s. solidissima* OTUs from each other and from *S.s. similis*, but because of historical hybridization, it will not always accurately reflect the ancestry of the nuclear genome. To diagnose OTU differences in areas with possible hybridization, a higher-resolution multilocus nDNA panel is required to quantify the relative admixture proportions. Because the selection of loci for the GTseq panel was not random across the genome, this tool does not necessarily provide absolute admixture proportions that represent the genomic average, but it can precisely assess relative proportions among the samples analyzed.

Previous studies have shown that *S.s. similis* has higher heat tolerance than *S.s. solidissima* and it has been suggested that a cross between the two taxa could produce a more heat-tolerant aquaculture strain (Acquafredda et al., 2021; Hurley & Walker, 1997; Walker, 1998; Weinberg, 2005). Inshore OTU-A may also have a higher heat tolerance than OTU-B and have different susceptibilities to climate change (Cerrato & Keith, 1992; Hofmann et al., 2018). The moderate frequency of hybrid individuals between OTUs A and B in nature, relative to rare *S.s. solidissima* x *S.s. similis* hybrids, may not be a good indicator of which hatchery crosses are more likely to yield a viable and useful aquaculture product (Mike Acquafredda pers. comm.). This is because there can be important reproductive barriers in nature (e.g., spawn timing) that are easily overcome with hatchery manipulation.

Sampling for this study provided the first definitive evidence of *S.s. similis* in the Mid- Atlantic Bight, geographically intermediate between known populations in southern New England and south of Cape Hatteras, as well as the first evidence for its occurrence on the continental shelf (but see Wisner et al. 2024). The *S.s. similis* collection locations suggest that the shelf occurrences of *S.s. similis* may generally be proximate to estuarine outflows. If nearshore *S.s. solidissima* fisheries include regions that are likely to contain *S.s. similis* stocks, it may be important for regulations to account for both taxa, especially if they are taxonomically elevated to full species, as recommended by Hare et al. (2010) and Wisner et al. (2024).

Similarly, if aquaculture seeding of surfclams on lease sites respects the natural range distribution limits, OTU-B *S.s. solidissima* allows for the least constraint on where to deploy seed clams, whereas our results indicate that *S.s. similis* and probably OTU-A have a natural northern range limit at Cape Cod. Further investigation should examine the sources of gene flow generating OTU AxB ancestry in some Cape Cod Bay locations.

## ACKNOWLEDGEMENTS

This work was supported by funding from the Mid-Atlantic Fishery Management Council under federal award #NA15NMF4410006. Thanks to Jessica Coakley and Daniel Hennen for help along the way, especially mitigating the pandemic. We are grateful to Matt Weeks and David Ullman for collecting surfclams and to Nicole Charriere, Peter Chase and Atlantic Capes Fisheries for supplying samples. Thanks also to Eric Robillard and the Northeast Fisheries Science Center Population & Ecosystems Monitoring & Analysis Division for their expertise and assistance in surfclam ageing. We would like to thank Harmony Borchardt-Wier for her invaluable assistance, advice, and support throughout this project, especially regarding library protocols and troubleshooting DNA quality.

## DATA AVAILABILITY

Fastq sequence reads were deposited in the SRA (BioProject PRJNA1493695). Sample metadata are in supplementary table S.1. VCF files and scripts were deposited in Cornell eCommons at https://doi.org/10.7298/asdq-g871.

## AUTHOR CONTRIBUTIONS

H. H. performed sample processing and extraction. H. H. and Y. C. conducted analyses and visualized the results. M. P. H. conceived the project, acquired funding and provided supervision. All authors wrote, edited and reviewed the manuscript.

## Ethical Compliance

Field collection of surf clams was conducted under permit number 2019- 1169 issued by the New York State Department of Environmental Conservation. All other samples were obtained from federal survey collections. No species listed as threatened, endangered, or protected under the IUCN Red List or local legislation were targeted or collected.

