## Supplemental Tables and Figures for "Genomic Population Structure of Atlantic surfclams: Cryptic Taxonomic Units and Population Connectivity"

#### Table S.1: Individual sample metadata

| sample | subspecies | collection year | Collection location or source | Analytical Group Abbreviation | STRUCTURE Q (OUT-A) | STRUCTURE Q (OTU-B) | ddRAD taxon, 10% hybrid threshold | ddRAD taxon, 1% hybrid threshold | majority taxon in hybrid | mtCOI *Sph* I | mtCOI *Bsp* 1286I | GTseq | analytical group location | Latitude | Longitude | Length (mm) | Age (yrs) | Width (mm) | Depth (mm) | weight (g) | water depth (m) | Region Avg water depth (m) | Notes |
|---|---|---|---|---|---|---|---|---|---|---|---|---|---|---|---|---|---|---|---|---|---|---|---|
| BRW_001 | solidissima | 2017 | Brewster, MA | CCB1 | 0 | 1 | OTU-B | B | | mtB | NA | | Cape Cod Bay | 41.770141 | -70.113109 | 146.8 | NA | 108 | 63.3 | 481 | 0.3 - 1.2 | 0.72 | |
| BRW_002 | solidissima | 2017 | Brewster, MA | CCB1 | 0.045 | 0.955 | OTU-B | Hyb | majority B | no amp | NA | | Cape Cod Bay | 41.770141 | -70.113109 | 161.9 | NA | 112 | 64.5 | 571.5 | 0.3 - 1.2 | | |
| BRW_003 | solidissima | 2017 | Brewster, MA | CCB1 | 0 | 1 | OTU-B | B | | no amp | NA | | Cape Cod Bay | 41.770141 | -70.113109 | 99.6 | NA | 72.6 | 44.4 | 135 | 0.3 - 1.2 | | |
| BRW_004 | solidissima | 2017 | Brewster, MA | CCB1 | 0 | 1 | OTU-B | B | | mtB | NA | | Cape Cod Bay | 41.770141 | -70.113109 | 153.6 | NA | 105.4 | 63.1 | 563 | 0.3 - 1.2 | | |
| BRW_005 | solidissima | 2017 | Brewster, MA | CCB1 | 0 | 1 | OTU-B | B | | mtA | NA | | Cape Cod Bay | 41.770141 | -70.113109 | 146.4 | NA | 102.2 | 64 | 406 | 0.3 - 1.2 | | |
| CGC_001 | solidissima | 2017 | Yarmouth, Chase Ga | CCB2 | 0 | 1 | OTU-B | B | | not done | NA | | Cape Cod Bay | 41.723891 | -70.236156 | 31.8 | NA | 25.4 | 15.1 | 7 | 0.3 - 1.2 | | |
| CGC_002 | solidissima | 2017 | Yarmouth, Chase Ga | CCB2 | 0 | 1 | OTU-B | B | | mtA | NA | | Cape Cod Bay | 41.723891 | -70.236156 | 62.5 | NA | 46.2 | 27 | 38 | 0.3 - 1.2 | | |
| CGC_003 | solidissima | 2017 | Yarmouth, Chase Ga | CCB2 | 0 | 1 | OTU-B | B | | mtB | NA | | Cape Cod Bay | 41.723891 | -70.236156 | 50.4 | NA | 39.1 | 22.7 | 24 | 0.3 - 1.2 | | |
| CGC_004 | solidissima | 2017 | Yarmouth, Chase Ga | CCB2 | 0 | 1 | OTU-B | B | | mtA | NA | | Cape Cod Bay | 41.723891 | -70.236156 | 72.2 | NA | 54.5 | 29.9 | 60 | 0.3 - 1.2 | | |
| CGC_005 | solidissima | 2017 | Yarmouth, Chase Ga | CCB2 | 0 | 1 | OTU-B | B | | mtB | NA | | Cape Cod Bay | 41.723891 | -70.236156 | 79.7 | NA | 62.9 | 37.2 | 93 | 0.3 - 1.2 | | |
| CGC_006 | solidissima | 2017 | Yarmouth, Chase Ga | CCB2 | 0.244 | 0.756 | OTU-Hyb | Hyb | majority B | mtB | NA | hyb | Cape Cod Bay | 41.723891 | -70.236156 | 92.6 | NA | 68 | 38.7 | 127 | 0.3 - 1.2 | | |
| CGC_007 | solidissima | 2017 | Yarmouth, Chase Ga | CCB2 | 0.703 | 0.297 | OTU-Hyb | Hyb | majority A | mtB | NA | hyb | Cape Cod Bay | 41.723891 | -70.236156 | 114 | NA | 82.6 | 47.8 | 245 | 0.3 - 1.2 | | |
| CGC_008 | solidissima | 2017 | Yarmouth, Chase Ga | CCB2 | 0.119 | 0.881 | OTU-Hyb | Hyb | majority B | mtB | NA | hyb | Cape Cod Bay | 41.723891 | -70.236156 | 105.6 | NA | 75.9 | 40.9 | 173.5 | 0.3 - 1.2 | | |
| CGC_009 | solidissima | 2017 | Yarmouth, Chase Ga | CCB2 | 0 | 1 | OTU-B | B | | mtB | NA | | Cape Cod Bay | 41.723891 | -70.236156 | 115.9 | NA | 83.3 | 48 | 250 | 0.3 - 1.2 | | |
| CGC_010 | solidissima | 2017 | Yarmouth, Chase Ga | CCB2 | 0 | 1 | OTU-B | B | | mtB | NA | | Cape Cod Bay | 41.723891 | -70.236156 | 111.7 | NA | 84.3 | NA | 177 | 0.3 - 1.2 | | broken shell |
| CGC_011 | solidissima | 2017 | Yarmouth, Chase Ga | CCB2 | 0.032 | 0.968 | OTU-B | Hyb | majority B | mtB | NA | | Cape Cod Bay | 41.723891 | -70.236156 | 120 | NA | 84.8 | 47.7 | 194 | 0.3 - 1.2 | | |
| BAR_001 | solidissima | 2017 | Barnstable, Barnstat | CCB3 | 0.444 | 0.556 | OTU-Hyb | Hyb | majority B | mtA | NA | hyb | Cape Cod Bay | 41.717159 | -70.280191 | 104.8 | NA | NA | 45.6 | NA | 0.6 - 1.2 | | |
| BAR_002 | solidissima | 2017 | Barnstable, Barnstat | CCB3 | 0.012 | 0.988 | OTU-B | Hyb | majority B | mtB | NA | | Cape Cod Bay | 41.717159 | -70.280191 | 127 | NA | 95 | 54.5 | 346 | 0.6 - 1.2 | | |
| BAR_003 | solidissima | 2017 | Barnstable, Barnstat | CCB3 | 0 | 1 | OTU-B | B | | mtB | NA | | Cape Cod Bay | 41.717159 | -70.280191 | 139.1 | NA | 103.8 | 59.6 | 507 | 0.6 - 1.2 | | |
| BAR_004 | solidissima | 2017 | Barnstable, Barnstat | CCB3 | 0.035 | 0.965 | OTU-B | Hyb | majority B | mtA | NA | | Cape Cod Bay | 41.717159 | -70.280191 | 57.6 | NA | 45 | 27.7 | 34 | 0.6 - 1.2 | | |
| BAR_005 | solidissima | 2017 | Barnstable, Barnstat | CCB3 | 0.014 | 0.986 | OTU-B | Hyb | majority B | mtB | NA | | Cape Cod Bay | 41.717159 | -70.280191 | 125.2 | NA | 89.6 | 53.6 | 303 | 0.6 - 1.2 | | |
| BAR_006 | solidissima | 2017 | Barnstable, Barnstat | CCB3 | 0 | 1 | OTU-B | B | | mtB | NA | | Cape Cod Bay | 41.717159 | -70.280191 | 118.4 | NA | 87.5 | 51.7 | 254 | 0.6 - 1.2 | | |
| BAR_007 | solidissima | 2017 | Barnstable, Barnstat | CCB3 | 0.003 | 0.997 | OTU-B | B | | mtB | NA | | Cape Cod Bay | 41.717159 | -70.280191 | 112.1 | NA | 84.1 | 49.1 | 226.5 | 0.6 - 1.2 | | |
| BAR_008 | solidissima | 2017 | Barnstable, Barnstat | CCB3 | 0 | 1 | OTU-B | B | | mtB | NA | | Cape Cod Bay | 41.717159 | -70.280191 | 145.7 | NA | 110.5 | 67 | 579 | 0.6 - 1.2 | | |
| BAR_009 | solidissima | 2017 | Barnstable, Barnstat | CCB3 | 0.001 | 0.999 | OTU-B | B | | mtB | NA | | Cape Cod Bay | 41.717159 | -70.280191 | 165 | NA | 108.3 | 67.6 | 577 | 0.6 - 1.2 | | |
| BAR_010 | solidissima | 2017 | Barnstable, Barnstat | CCB3 | 0 | 1 | OTU-B | B | | mtB | NA | | Cape Cod Bay | 41.717159 | -70.280191 | 96.9 | NA | 76.5 | 44.2 | 126.5 | 0.6 - 1.2 | | |
| PLY_001 | solidissima | 2017 | Plymouth, Plymouth Ba | CCB4 | 0.636 | 0.364 | OTU-Hyb | Hyb | majority A | mtA | NA | hyb | Cape Cod Bay | 41.983005 | -70.651725 | 114.8 | NA | 81.5 | 46.5 | 200 | 0.3 - 1.2 | | |
| PLY_002 | solidissima | 2017 | Plymouth, Plymouth Ba | CCB4 | 0.29 | 0.71 | OTU-Hyb | Hyb | majority B | mtB | NA | hyb | Cape Cod Bay | 41.983005 | -70.651725 | 101.3 | NA | 69.8 | 40.9 | 151 | 0.3 - 1.2 | | |
| PLY_003 | solidissima | 2017 | Plymouth, Plymouth Ba | CCB4 | 0 | 1 | OTU-B | B | | mtB | NA | | Cape Cod Bay | 41.983005 | -70.651725 | 141.7 | NA | 101.3 | 59.2 | 442 | 0.3 - 1.2 | | |
| PLY_004 | solidissima | 2017 | Plymouth, Plymouth Ba | CCB4 | 0.05 | 0.95 | OTU-B | Hyb | majority B | mtA | NA | | Cape Cod Bay | 41.983005 | -70.651725 | 139.2 | NA | 99.3 | 55.7 | 423 | 0.3 - 1.2 | | |
| PLY_005 | solidissima | 2017 | Plymouth, Plymouth Ba | CCB4 | 0.012 | 0.988 | OTU-B | Hyb | majority B | mtB | NA | | Cape Cod Bay | 41.983005 | -70.651725 | 148.4 | NA | 107.7 | 67.2 | 534 | 0.3 - 1.2 | | internal shell deformity |
| PLY_006 | solidissima | 2017 | Plymouth, Plymouth Ba | CCB4 | 0.054 | 0.946 | OTU-B | Hyb | majority B | mtB | NA | | Cape Cod Bay | 41.983005 | -70.651725 | 148.7 | NA | 105.2 | 62.1 | 413.5 | 0.3 - 1.2 | | |
| PLY_007 | solidissima | 2017 | Plymouth, Plymouth Ba | CCB4 | 0 | 1 | OTU-B | B | | mtA | NA | | Cape Cod Bay | 41.983005 | -70.651725 | 153.5 | NA | 104.5 | 63.1 | 584 | 0.3 - 1.2 | | |
| PLY_008 | solidissima | 2017 | Plymouth, Plymouth Ba | CCB4 | 0.002 | 0.998 | OTU-B | B | | mtA | NA | | Cape Cod Bay | 41.983005 | -70.651725 | 163 | NA | 111.1 | 71.9 | 886.5 | 0.3 - 1.2 | | |
| PLY_009 | solidissima | 2017 | Plymouth, Plymouth Ba | CCB4 | 0 | 1 | OTU-B | B | | mtB | NA | | Cape Cod Bay | 41.983005 | -70.651725 | 166 | NA | 113.8 | 69.2 | 804 | 0.3 - 1.2 | | |
| PLY_010 | solidissima | 2017 | Plymouth, Plymouth Ba | CCB4 | 0.81 | 0.19 | OTU-Hyb | Hyb | majority A | mtA | NA | hyb | Cape Cod Bay | 41.983005 | -70.651725 | 180 | NA | 115.1 | 66.3 | 588 | 0.3 - 1.2 | | broken shell |
| PT_003 | solidissima | 2012 | Provincetown, MA | CCB5 | 0.001 | 0.999 | OTU-B | B | | no amp | NA | B | Cape Cod Bay | 42.04 | -70.15 | 161 | NA | NA | NA | NA | NA | | |
| PT_006 | solidissima | 2012 | Provincetown, MA | CCB5 | 0 | 1 | OTU-B | B | | mtB | NA | B | Cape Cod Bay | 42.04 | -70.15 | 178 | NA | NA | NA | NA | NA | | |
| PT_008 | solidissima | 2012 | Provincetown, MA | CCB5 | 0.006 | 0.994 | OTU-B | B | | mtB | NA | B | Cape Cod Bay | 42.04 | -70.15 | 169 | NA | NA | NA | NA | NA | | |
| PT_009 | solidissima | 2012 | Provincetown, MA | CCB5 | 0 | 1 | OTU-B | B | | mtB | NA | B | Cape Cod Bay | 42.04 | -70.15 | 151 | NA | NA | NA | NA | NA | | |
| PT_010 | solidissima | 2012 | Provincetown, MA | CCB5 | 0.012 | 0.988 | OTU-B | Hyb | majority B | mtA | NA | B | Cape Cod Bay | 42.04 | -70.15 | 170 | NA | NA | NA | NA | NA | | |
| PT_011 | solidissima | 2012 | Provincetown, MA | CCB5 | 0.002 | 0.998 | OTU-B | B | | mtB | NA | B | Cape Cod Bay | 42.04 | -70.15 | 170 | NA | NA | NA | NA | NA | | |
| PT_012 | solidissima | 2012 | Provincetown, MA | CCB5 | 0 | 1 | OTU-B | B | | mtB | NA | B | Cape Cod Bay | 42.04 | -70.15 | 173 | NA | NA | NA | NA | NA | | |
| PT_013 | solidissima | 2012 | Provincetown, MA | CCB5 | 0 | 1 | OTU-B | B | | mtB | NA | B | Cape Cod Bay | 42.04 | -70.15 | 172 | NA | NA | NA | NA | NA | | |
| PT_014 | solidissima | 2012 | Provincetown, MA | CCB5 | 0 | 1 | OTU-B | B | | mtA | NA | B | Cape Cod Bay | 42.04 | -70.15 | 166 | NA | NA | NA | NA | NA | | |
| PT_016 | solidissima | 2012 | Provincetown, MA | CCB5 | 0.108 | 0.892 | OTU-Hyb | Hyb | majority B | mtB | NA | hyb | Cape Cod Bay | 42.04 | -70.15 | 167 | NA | NA | NA | NA | NA | | |
| PT_017 | solidissima | 2012 | Provincetown, MA | CCB5 | 0.007 | 0.993 | OTU-B | B | | mtB | NA | B | Cape Cod Bay | 42.04 | -70.15 | 171 | NA | NA | NA | NA | NA | | |
| PT_018 | solidissima | 2012 | Provincetown, MA | CCB5 | 0 | 1 | OTU-B | B | | mtB | NA | B | Cape Cod Bay | 42.04 | -70.15 | 156 | NA | NA | NA | NA | NA | | |
| PT_019 | solidissima | 2012 | Provincetown, MA | CCB5 | 0.069 | 0.931 | OTU-B | Hyb | majority B | mtA | NA | B | Cape Cod Bay | 42.04 | -70.15 | 160 | NA | NA | NA | NA | NA | | |
| PT_01n | solidissima | 2012 | Provincetown, MA | CCB5 | 0 | 1 | OTU-B | B | | no amp | NA | B | Cape Cod Bay | 42.04 | -70.15 | 167 | NA | NA | NA | NA | NA | | |
| PT_020 | solidissima | 2012 | Provincetown, MA | CCB5 | 0 | 1 | OTU-B | B | | mtB | NA | B | Cape Cod Bay | 42.04 | -70.15 | 165 | NA | NA | NA | NA | NA | | |
| PT_04n | solidissima | 2012 | Provincetown, MA | CCB5 | 0.134 | 0.866 | OTU-Hyb | Hyb | majority B | mtB | NA | B | Cape Cod Bay | 42.04 | -70.15 | NA | NA | NA | NA | NA | NA | | |
| PT_05n | solidissima | 2012 | Provincetown, MA | CCB5 | 0.027 | 0.973 | OTU-B | Hyb | majority B | mtB | NA | B | Cape Cod Bay | 42.04 | -70.15 | NA | NA | NA | NA | NA | NA | | |
| PT_07n | solidissima | 2012 | Provincetown, MA | CCB5 | 0.023 | 0.977 | OTU-B | Hyb | majority B | mtB | NA | B | Cape Cod Bay | 42.04 | -70.15 | 165 | NA | NA | NA | NA | NA | | |
| PVT_001 | solidissima | 2017 | Provincetown Harbor, M | CCB6 | 0.164 | 0.836 | OTU-Hyb | Hyb | majority B | mtB | NA | hyb | Cape Cod Bay | 42.060706 | -70.159156 | 104.1 | NA | 76.3 | 44.6 | 171 | 0 - 0.9 | | |
| PVT_002 | solidissima | 2017 | Provincetown Harbor, M | CCB6 | 0.001 | 0.999 | OTU-B | B | | no amp | NA | | Cape Cod Bay | 42.060706 | -70.159156 | 97.9 | NA | 72.1 | 45.4 | 159 | 0 - 0.9 | | |
| PVT_003 | solidissima | 2017 | Provincetown Harbor, M | CCB6 | 0 | 1 | OTU-B | B | | no amp | NA | | Cape Cod Bay | 42.060706 | -70.159156 | 109.3 | NA | 82.1 | 45.8 | 202 | 0 - 0.9 | | |
| PVT_004 | solidissima | 2017 | Provincetown Harbor, M | CCB6 | 0 | 1 | OTU-B | B | | no amp | NA | | Cape Cod Bay | 42.060706 | -70.159156 | 83.2 | NA | 65.9 | 38.5 | 110 | 0 - 0.9 | | |
| PVT_005 | solidissima | 2017 | Provincetown Harbor, M | CCB6 | 0.227 | 0.773 | OTU-Hyb | Hyb | majority B | mtB | NA | hyb | Cape Cod Bay | 42.060706 | -70.159156 | 99.7 | NA | 75.9 | 44.7 | 167 | 0 - 0.9 | | |
| PVT_006 | solidissima | 2017 | Provincetown Harbor, M | CCB6 | 0 | 1 | OTU-B | B | | no amp | NA | | Cape Cod Bay | 42.060706 | -70.159156 | 103 | NA | 76.2 | 46.7 | 187 | 0 - 0.9 | | |
| PVT_007 | solidissima | 2017 | Provincetown Harbor, M | CCB6 | 0 | 1 | OTU-B | B | | no amp | NA | | Cape Cod Bay | 42.060706 | -70.159156 | 85.3 | NA | 65.3 | 42.9 | 117 | 0 - 0.9 | | |
| PVT_008 | solidissima | 2017 | Provincetown Harbor, M | CCB6 | 0.014 | 0.986 | OTU-B | Hyb | majority B | no amp | NA | | Cape Cod Bay | 42.060706 | -70.159156 | 107.7 | NA | 78.7 | 47.7 | 185 | 0 - 0.9 | | |
| PVT_009 | solidissima | 2017 | Provincetown Harbor, M | CCB6 | 0.05 | 0.95 | OTU-B | Hyb | majority B | no amp | NA | | Cape Cod Bay | 42.060706 | -70.159156 | 103.5 | NA | 78.8 | 48.7 | 173 | 0 - 0.9 | | |
| PVT_010 | solidissima | 2017 | Provincetown Harbor, M | CCB6 | 0 | 1 | OTU-B | B | | no amp | NA | | Cape Cod Bay | 42.060706 | -70.159156 | 95.3 | NA | 70.9 | 43.8 | 149 | 0 - 0.9 | | |
| FNJ_001 | solidissima | 2020 | Off Fenwick Island, De | DEL1 | 0 | 1 | OTU-B | B | | mtA | sol | B | Delmarva | 38.46 | -74.9 | 128.9 | NA | 88.5 | NA | NA | NA | 37.15 | |
| FNJ_002 | solidissima | 2020 | Off Fenwick Island, De | DEL1 | 0 | 1 | OTU-B | B | | mtB | sol | B | Delmarva | 38.46 | -74.9 | 125.9 | NA | 87.2 | NA | NA | NA | | |
| FNJ_003 | solidissima | 2020 | Off Fenwick Island, De | DEL1 | 0.001 | 0.999 | OTU-B | B | | mtB | sol | B | Delmarva | 38.46 | -74.9 | 116.9 | NA | 84.1 | NA | NA | NA | | |
| FNJ_004 | solidissima | 2020 | Off Fenwick Island, De | DEL1 | 0.001 | 0.999 | OTU-B | B | | mtB | sol | | Delmarva | 38.46 | -74.9 | 120.4 | NA | 87.7 | NA | NA | NA | | |
| FNJ_005 | solidissima | 2020 | Off Fenwick Island, De | DEL1 | 0 | 1 | OTU-B | B | | mtB | sol | | Delmarva | 38.46 | -74.9 | 119.2 | NA | 88 | NA | NA | NA | | |
| FNJ_006 | solidissima | 2020 | Off Fenwick Island, De | DEL1 | 0 | 1 | OTU-B | B | | mtB | sol | | Delmarva | 38.46 | -74.9 | 126.2 | NA | 87.5 | NA | NA | NA | | |
| FNJ_007 | solidissima | 2020 | Off Fenwick Island, De | DEL1 | 0.031 | 0.969 | OTU-B | Hyb | majority B | mtB | sol | | Delmarva | 38.46 | -74.9 | 123.5 | NA | 85.8 | NA | NA | NA | | |
| FNJ_008 | solidissima | 2020 | Off Fenwick Island, De | DEL1 | 0 | 1 | OTU-B | B | | mtB | sol | | Delmarva | 38.46 | -74.9 | 120.8 | NA | 83.1 | NA | NA | NA | | |
| FNJ_009 | solidissima | 2020 | Off Fenwick Island, De | DEL1 | 0 | 1 | OTU-B | B | | mtB | sol | | Delmarva | 38.46 | -74.9 | 119.4 | NA | 87.3 | NA | NA | NA | | |
| FNJ_010 | solidissima | 2020 | Off Fenwick Island, De | DEL1 | 0 | 1 | OTU-B | B | | mtB | sol | | Delmarva | 38.46 | -74.9 | 125.8 | NA | 90.5 | NA | NA | NA | | |
| FNJ_011 | solidissima | 2020 | Off Fenwick Island, De | DEL1 | 0 | 1 | OTU-B | B | | mtB | sol | | Delmarva | 38.46 | -74.9 | 122.1 | NA | 86 | NA | NA | NA | | |
| FNJ_012 | solidissima | 2020 | Off Fenwick Island, De | DEL1 | 0 | 1 | OTU-B | B | | mtB | sol | | Delmarva | 38.46 | -74.9 | 123.2 | NA | 88.7 | NA | NA | NA | | |
| FNJ_013 | solidissima | 2020 | Off Fenwick Island, De | DEL1 | 0 | 1 | OTU-B | B | | mtB | sol | | Delmarva | 38.46 | -74.9 | 123.8 | NA | 86.2 | NA | NA | NA | | |
| FNJ_014 | solidissima | 2020 | Off Fenwick Island, De | DEL1 | 0 | 1 | OTU-B | B | | mtA | sol | | Delmarva | 38.46 | -74.9 | 123.7 | NA | 88.1 | NA | NA | NA | | |
| FNJ_015 | solidissima | 2020 | Off Fenwick Island, De | DEL1 | 0 | 1 | OTU-B | B | | mtA | sol | | Delmarva | 38.46 | -74.9 | 123.2 | NA | 90.3 | NA | NA | NA | | |
| FNJ_016 | solidissima | 2020 | Off Fenwick Island, De | DEL1 | 0 | 1 | OTU-B | B | | mtB | sol | | Delmarva | 38.46 | -74.9 | 119.2 | NA | 82.2 | NA | NA | NA | | |
| FNJ_017 | solidissima | 2020 | Off Fenwick Island, De | DEL1 | 0 | 1 | OTU-B | B | | mtA | sol | | Delmarva | 38.46 | -74.9 | 128.7 | NA | 86.2 | NA | NA | NA | | |
| FNJ_018 | solidissima | 2020 | Off Fenwick Island, De | DEL1 | 0 | 1 | OTU-B | B | | mtB | sol | | Delmarva | 38.46 | -74.9 | 113.7 | NA | 80.5 | NA | NA | NA | | |
| FNJ_019 | solidissima | 2020 | Off Fenwick Island, De | DEL1 | 0.012 | 0.988 | OTU-B | Hyb | majority B | mtB | sol | | Delmarva | 38.46 | -74.9 | 119.2 | NA | 83.3 | NA | NA | NA | | |
| FNJ_020 | solidissima | 2020 | Off Fenwick Island, De | DEL1 | 0 | 1 | OTU-B | B | | mtB | sol | | Delmarva | 38.46 | -74.9 | 114.6 | NA | 84.8 | NA | NA | NA | | |
| FNJ_021 | solidissima | 2020 | Off Fenwick Island, De | DEL1 | 0 | 1 | OTU-B | B | | mtB | sol | | Delmarva | 38.46 | -74.9 | 120.1 | NA | 83.7 | NA | NA | NA | | |
| FNJ_022 | solidissima | 2020 | Off Fenwick Island, De | DEL1 | 0 | 1 | OTU-B | B | | mtB | sol | | Delmarva | 38.46 | -74.9 | 127.3 | NA | 88.8 | NA | NA | NA | | |
| FNJ_023 | solidissima | 2020 | Off Fenwick Island, De | DEL1 | 0 | 1 | OTU-B | B | | mtB | sol | | Delmarva | 38.46 | -74.9 | 121.5 | NA | 83.7 | NA | NA | NA | | |
| FNJ_024 | solidissima | 2020 | Off Fenwick Island, De | DEL1 | 0 | 1 | OTU-B | B | | mtB | sol | | Delmarva | 38.46 | -74.9 | 125.8 | NA | 87.3 | NA | NA | NA | | |
| FNJ_025 | solidissima | 2020 | Off Fenwick Island, De | DEL1 | 0 | 1 | OTU-B | B | | mtB | sol | | Delmarva | 38.46 | -74.9 | 116.7 | NA | 84 | NA | NA | NA | | |
| FNJ_026 | solidissima | 2020 | Off Fenwick Island, De | DEL1 | 0 | 1 | OTU-B | B | | mtB | sol | | Delmarva | 38.46 | -74.9 | 121.2 | NA | 85.8 | NA | NA | NA | | |
| FNJ_027 | solidissima | 2020 | Off Fenwick Island, De | DEL1 | 0 | 1 | OTU-B | B | | mtB | sol | | Delmarva | 38.46 | -74.9 | 125.4 | NA | 84.3 | NA | NA | NA | | |
| FNJ_030 | solidissima | 2020 | Off Fenwick Island, De | DEL1 | 0.014 | 0.986 | OTU-B | Hyb | majority B | mtB | sol | | Delmarva | 38.46 | -74.9 | 120.3 | NA | 89.8 | NA | NA | NA | | |
| FNJ_031 | solidissima | 2020 | Off Fenwick Island, De | DEL1 | 0.048 | 0.952 | OTU-B | Hyb | majority B | mtB | sol | | Delmarva | 38.46 | -74.9 | 121.1 | NA | 87.4 | NA | NA | NA | | |
| FNJ_032 | solidissima | 2020 | Off Fenwick Island, De | DEL1 | 0 | 1 | OTU-B | B | | mtB | sol | | Delmarva | 38.46 | -74.9 | 129.5 | NA | 91.2 | NA | NA | NA | | |
| FNJ_033 | solidissima | 2020 | Off Fenwick Island, De | DEL1 | 0 | 1 | OTU-B | B | | mtB | sol | | Delmarva | 38.46 | -74.9 | 121.8 | NA | 83.6 | NA | NA | NA | | |
| FNJ_034 | solidissima | 2020 | Off Fenwick Island, De | DEL1 | 0 | 1 | OTU-B | B | | mtB | sol | | Delmarva | 38.46 | -74.9 | 117.9 | NA | 84.1 | NA | NA | NA | | |
| FNJ_035 | solidissima | 2020 | Off Fenwick Island, De | DEL1 | 0 | 1 | OTU-B | B | | mtB | sol | | Delmarva | 38.46 | -74.9 | 127.6 | NA | 90.5 | NA | NA | NA | | |
| FNJ_036 | solidissima | 2020 | Off Fenwick Island, De | DEL1 | 0.001 | 0.999 | OTU-B | B | | mtB | sol | | Delmarva | 38.46 | -74.9 | 122.5 | NA | 89.4 | NA | NA | NA | | |
| FNJ_037 | solidissima | 2020 | Off Fenwick Island, De | DEL1 | 0.001 | 0.999 | OTU-B | B | | mtB | sol | | Delmarva | 38.46 | -74.9 | 120.4 | NA | 83.5 | NA | NA | NA | | |
| FNJ_038 | solidissima | 2020 | Off Fenwick Island, De | DEL1 | 0 | 1 | OTU-B | B | | mtA | sol | | Delmarva | 38.46 | -74.9 | 115.1 | NA | 80.8 | NA | NA | NA | | |
| FNJ_039 | solidissima | 2020 | Off Fenwick Island, De | DEL1 | 0 | 1 | OTU-B | B | | mtB | sol | | Delmarva | 38.46 | -74.9 | 120.8 | NA | 86.2 | NA | NA | NA | | |
| FNJ_040 | solidissima | 2020 | Off Fenwick Island, De | DEL1 | 0 | 1 | OTU-B | B | | mtB | sol | | Delmarva | 38.46 | -74.9 | 124.4 | NA | 91.2 | NA | NA | NA | | |
| FNJ_041 | solidissima | 2020 | Off Fenwick Island, De | DEL1 | 0 | 1 | OTU-B | B | | mtB | sol | | Delmarva | 38.46 | -74.9 | 121.1 | NA | 85.9 | NA | NA | NA | | |
| FNJ_042 | solidissima | 2020 | Off Fenwick Island, De | DEL1 | 0 | 1 | OTU-B | B | | mtB | sol | | Delmarva | 38.46 | -74.9 | 121 | NA | 86.6 | NA | NA | NA | | |
| FNJ_043 | solidissima | 2020 | Off Fenwick Island, De | DEL1 | 0 | 1 | OTU-B | B | | mtB | sol | | Delmarva | 38.46 | -74.9 | 128.2 | NA | 92.7 | NA | NA | NA | | |
| FNJ_044 | solidissima | 2020 | Off Fenwick Island, De | DEL1 | 0 | 1 | OTU-B | B | | mtB | sol | | Delmarva | 38.46 | -74.9 | 120.3 | NA | 88.7 | NA | NA | NA | | |
| FNJ_045 | solidissima | 2020 | Off Fenwick Island, De | DEL1 | 0 | 1 | OTU-B | B | | mtB | sol | | Delmarva | 38.46 | -74.9 | 124.2 | NA | 88.4 | NA | NA | NA | | |
| FNJ_046 | solidissima | 2020 | Off Fenwick Island, De | DEL1 | 0 | 1 | OTU-B | B | | not done | NA | | Delmarva | 38.46 | -74.9 | 126.7 | NA | 86.1 | NA | NA | NA | | |
| FNJ_047 | solidissima | 2020 | Off Fenwick Island, De | DEL1 | 0 | 1 | OTU-B | B | | not done | NA | | Delmarva | 38.46 | -74.9 | 119.3 | NA | 86.3 | NA | NA | NA | | |
| FNJ_048 | solidissima | 2020 | Off Fenwick Island, De | DEL1 | 0 | 1 | OTU-B | B | | not done | NA | | Delmarva | 38.46 | -74.9 | 125.4 | NA | 90.6 | NA | NA | NA | | |
| FNJ_049 | solidissima | 2020 | Off Fenwick Island, De | DEL1 | 0.007 | 0.993 | OTU-B | B | | not done | NA | | Delmarva | 38.46 | -74.9 | 127.8 | NA | 91.6 | NA | NA | NA | | |
| FNJ_050 | solidissima | 2020 | Off Fenwick Island, De | DEL1 | 0 | 1 | OTU-B | B | | not done | NA | | Delmarva | 38.46 | -74.9 | 119 | NA | 86.8 | NA | NA | NA | | |
| FNJ_051 | solidissima | 2020 | Off Fenwick Island, De | DEL1 | 0 | 1 | OTU-B | B | | not done | NA | | Delmarva | 38.46 | -74.9 | 123.5 | NA | 88 | NA | NA | NA | | |
| 413_001 | solidissima | 1999 | 1999 Federal Survey | DEL2 | 0 | 1 | OTU-B | B | | mtB | NA | | Delmarva | 37.5023 | -75.1025 | 86 | 4 | NA | NA | NA | NA | | |
| 413_002 | solidissima | 1999 | 1999 Federal Survey | DEL2 | 0 | 1 | OTU-B | B | | mtB | NA | | Delmarva | 37.5023 | -75.1025 | 80 | 4 | NA | NA | NA | NA | | |
| 413_003 | solidissima | 1999 | 1999 Federal Survey | DEL2 | 0 | 1 | OTU-B | B | | mtA | sol | | Delmarva | 37.5023 | -75.1025 | 96 | 4 | NA | NA | NA | NA | | |
| 413_004 | solidissima | 1999 | 1999 Federal Survey | DEL2 | 0 | 1 | OTU-B | B | | NA | sol | | Delmarva | 37.5023 | -75.1025 | 90 | 4 | NA | NA | NA | NA | | |
| 413_005 | solidissima | 1999 | 1999 Federal Survey | DEL2 | 0 | 1 | OTU-B | B | | NA | sol | | Delmarva | 37.5023 | -75.1025 | 86 | 4 | NA | NA | NA | NA | | |
| 413_006 | solidissima | 1999 | 1999 Federal Survey | DEL2 | 0 | 1 | OTU-B | B | | NA | sol | | Delmarva | 37.5023 | -75.1025 | 78 | 4 | NA | NA | NA | NA | | |
| 413_007 | solidissima | 1999 | 1999 Federal Survey | DEL2 | 0 | 1 | OTU-B | B | | NA | sol | | Delmarva | 37.5023 | -75.1025 | 80 | 4 | NA | NA | NA | NA | | |
| 413_008 | solidissima | 1999 | 1999 Federal Survey | DEL2 | 0.001 | 0.999 | OTU-B | B | | mtB | sol | | Delmarva | 37.5023 | -75.1025 | 93 | 4 | NA | NA | NA | NA | | |
| 413_009 | solidissima | 1999 | 1999 Federal Survey | DEL2 | 0 | 1 | OTU-B | B | | mtA | sol | | Delmarva | 37.5023 | -75.1025 | 95 | 4 | NA | NA | NA | NA | | |
| 413_010 | solidissima | 1999 | 1999 Federal Survey | DEL2 | 0 | 1 | OTU-B | B | | mtB | sol | | Delmarva | 37.5023 | -75.1025 | 100 | 6 | NA | NA | NA | NA | | |
| 413_011 | solidissima | 1999 | 1999 Federal Survey | DEL2 | 0 | 1 | OTU-B | B | | mtB | sol | | Delmarva | 37.5023 | -75.1025 | 103 | 16 | NA | NA | NA | NA | | |
| 413_012 | solidissima | 1999 | 1999 Federal Survey | DEL2 | 0 | 1 | OTU-B | B | | mtB | sol | | Delmarva | 37.5023 | -75.1025 | 115 | 21 | NA | NA | NA | NA | | |
| 413_013 | solidissima | 1999 | 1999 Federal Survey | DEL2 | 0 | 1 | OTU-B | B | | NA | NA | | Delmarva | 37.5023 | -75.1025 | 110 | 20 | NA | NA | NA | NA | | |
| 413_014 | solidissima | 1999 | 1999 Federal Survey | DEL2 | 0 | 1 | OTU-B | B | | NA | NA | | Delmarva | 37.5023 | -75.1025 | 108 | 20 | NA | NA | NA | NA | | |
| 413_015 | solidissima | 1999 | 1999 Federal Survey | DEL2 | 0 | 1 | OTU-B | B | | NA | NA | | Delmarva | 37.5023 | -75.1025 | 108 | 20 | NA | NA | NA | NA | | |
| 413_016 | solidissima | 1999 | 1999 Federal Survey | DEL2 | 0 | 1 | OTU-B | B | | NA | NA | | Delmarva | 37.5023 | -75.1025 | 114 | 16 | NA | NA | NA | NA | | |
| 413_017 | solidissima | 1999 | 1999 Federal Survey | DEL2 | 0.001 | 0.999 | OTU-B | B | | mtA | sol | | Delmarva | 37.5023 | -75.1025 | 105 | 20 | NA | NA | NA | NA | | |
| 413_018 | solidissima | 1999 | 1999 Federal Survey | DEL2 | 0 | 1 | OTU-B | B | | mtA | sol | | Delmarva | 37.5023 | -75.1025 | 103 | 6 | NA | NA | NA | NA | | |
| 413_019 | solidissima | 1999 | 1999 Federal Survey | DEL2 | 0.031 | 0.969 | OTU-B | Hyb | majority B | mtB | sol | | Delmarva | 37.5023 | -75.1025 | 109 | 20 | NA | NA | NA | NA | | |
| 413_020 | solidissima | 1999 | 1999 Federal Survey | DEL2 | 0 | 1 | OTU-B | B | | mtA | sol | | Delmarva | 37.5023 | -75.1025 | 115 | 20 | NA | NA | NA | NA | | |
| 413_021 | solidissima | 1999 | 1999 Federal Survey | DEL2 | 0 | 1 | OTU-B | B | | mtB | sol | | Delmarva | 37.5023 | -75.1025 | 110 | 19 | NA | NA | NA | NA | | |
| 413_022 | solidissima | 1999 | 1999 Federal Survey | DEL2 | 0 | 1 | OTU-B | B | | mtB | sol | | Delmarva | 37.5023 | -75.1025 | 108 | 20 | NA | NA | NA | NA | | |
| 413_023 | solidissima | 1999 | 1999 Federal Survey | DEL2 | 0 | 1 | OTU-B | B | | mtB | sol | | Delmarva | 37.5023 | -75.1025 | 106 | 20 | NA | NA | NA | NA | | |
| 413_024 | solidissima | 1999 | 1999 Federal Survey | DEL2 | 0 | 1 | OTU-B | B | | mtA | sol | | Delmarva | 37.5023 | -75.1025 | 116 | 18 | NA | NA | NA | NA | | |
| 413_025 | solidissima | 1999 | 1999 Federal Survey | DEL2 | 0 | 1 | OTU-B | B | | mtB | sol | | Delmarva | 37.5023 | -75.1025 | 114 | 22 | NA | NA | NA | NA | | |
| 413_026 | solidissima | 1999 | 1999 Federal Survey | DEL2 | 0 | 1 | OTU-B | B | | mtB | sol | | Delmarva | 37.5023 | -75.1025 | 125 | 20 | NA | NA | NA | NA | | |
| 413_027 | solidissima | 1999 | 1999 Federal Survey | DEL2 | 0 | 1 | OTU-B | B | | mtA | sol | | Delmarva | 37.5023 | -75.1025 | 120 | 19 | NA | NA | NA | NA | | |
| 413_028 | solidissima | 1999 | 1999 Federal Survey | DEL2 | 0 | 1 | OTU-B | B | | mtA | sol | | Delmarva | 37.5023 | -75.1025 | 121 | 20 | NA | NA | NA | NA | | |
| 413_029 | solidissima | 1999 | 1999 Federal Survey | DEL2 | 0 | 1 | OTU-B | B | | mtB | sol | | Delmarva | 37.5023 | -75.1025 | 120 | 20 | NA | NA | NA | NA | | |
| 413_030 | solidissima | 1999 | 1999 Federal Survey | DEL2 | 0 | 1 | OTU-B | B | | mtA | sol | | Delmarva | 37.5023 | -75.1025 | 117 | 20 | NA | NA | NA | NA | | |
| 413_031 | solidissima | 1999 | 1999 Federal Survey | DEL2 | 0 | 1 | OTU-B | B | | mtB | sol | | Delmarva | 37.5023 | -75.1025 | 120 | 20 | NA | NA | NA | NA | | |
| 413_032 | solidissima | 1999 | 1999 Federal Survey | DEL2 | 0 | 1 | OTU-B | B | | mtA | sol | | Delmarva | 37.5023 | -75.1025 | 117 | 18 | NA | NA | NA | NA | | |
| 413_033 | solidissima | 1999 | 1999 Federal Survey | DEL2 | 0 | 1 | OTU-B | B | | mtB | sol | | Delmarva | 37.5023 | -75.1025 | 117 | 21 | NA | NA | NA | NA | | |
| 413_034 | solidissima | 1999 | 1999 Federal Survey | DEL2 | 0 | 1 | OTU-B | B | | mtB | sol | | Delmarva | 37.5023 | -75.1025 | 116 | 21 | NA | NA | NA | NA | | |
| 413_035 | solidissima | 1999 | 1999 Federal Survey | DEL2 | 0 | 1 | OTU-B | B | | mtB | sol | | Delmarva | 37.5023 | -75.1025 | 106 | 20 | NA | NA | NA | NA | | |
| 413_036 | solidissima | 1999 | 1999 Federal Survey | DEL2 | 0 | 1 | OTU-B | B | | mtB | sol | | Delmarva | 37.5023 | -75.1025 | 114 | 20 | NA | NA | NA | NA | | |
| 413_037 | solidissima | 1999 | 1999 Federal Survey | DEL2 | 0 | 1 | OTU-B | B | | mtB | sol | | Delmarva | 37.5023 | -75.1025 | 119 | 20 | NA | NA | NA | NA | | |
| 413_038 | solidissima | 1999 | 1999 Federal Survey | DEL2 | 0 | 1 | OTU-B | B | | mtB | sol | | Delmarva | 37.5023 | -75.1025 | 126 | 21 | NA | NA | NA | NA | | |
| 413_039 | solidissima | 1999 | 1999 Federal Survey | DEL2 | 0 | 1 | OTU-B | B | | mtB | sol | | Delmarva | 37.5023 | -75.1025 | 125 | 20 | NA | NA | NA | NA | | |
| 413_040 | solidissima | 1999 | 1999 Federal Survey | DEL2 | 0 | 1 | OTU-B | B | | mtB | sol | | Delmarva | 37.5023 | -75.1025 | 124 | 20 | NA | NA | NA | NA | | |
| 413_041 | solidissima | 1999 | 1999 Federal Survey | DEL2 | 0 | 1 | OTU-B | B | | mtB | sol | | Delmarva | 37.5023 | -75.1025 | 127 | 22 | NA | NA | NA | NA | | |
| 413_042 | solidissima | 1999 | 1999 Federal Survey | DEL2 | 0.017 | 0.983 | OTU-B | Hyb | majority B | mtB | sol | | Delmarva | 37.5023 | -75.1025 | 122 | 20 | NA | NA | NA | NA | | |
| 413_043 | solidissima | 1999 | 1999 Federal Survey | DEL2 | 0 | 1 | OTU-B | B | | mtB | sol | | Delmarva | 37.5023 | -75.1025 | 124 | 22 | NA | NA | NA | NA | | |
| 413_044 | solidissima | 1999 | 1999 Federal Survey | DEL2 | 0 | 1 | OTU-B | B | | mtB | sol | | Delmarva | 37.5023 | -75.1025 | 129 | 20 | NA | NA | NA | NA | | |
| 413_045 | solidissima | 1999 | 1999 Federal Survey | DEL2 | 0 | 1 | OTU-B | B | | mtB | sol | | Delmarva | 37.5023 | -75.1025 | 128 | 20 | NA | NA | NA | NA | | |
| 413_046 | solidissima | 1999 | 1999 Federal Survey | DEL2 | 0 | 1 | OTU-B | B | | mtA | sol | | Delmarva | 37.5023 | -75.1025 | 123 | 19 | NA | NA | NA | NA | | |
| 413_047 | solidissima | 1999 | 1999 Federal Survey | DEL2 | 0 | 1 | OTU-B | B | | mtA | sol | | Delmarva | 37.5023 | -75.1025 | 132 | 22 | NA | NA | NA | NA | | |
| 413_048 | solidissima | 1999 | 1999 Federal Survey | DEL2 | 0 | 1 | OTU-B | B | | mtB | sol | | Delmarva | 37.5023 | -75.1025 | 136 | 21 | NA | NA | NA | NA | | |
| 413_049 | solidissima | 1999 | 1999 Federal Survey | DEL2 | 0 | 1 | OTU-B | B | | mtB | sol | | Delmarva | 37.5023 | -75.1025 | 126 | 22 | NA | NA | NA | NA | | |
| 413_050 | solidissima | 1999 | 1999 Federal Survey | DEL2 | 0 | 1 | OTU-B | B | | mtB | sol | | Delmarva | 37.5023 | -75.1025 | 125 | 20 | NA | NA | NA | NA | | |
| MAB22_07926 | solidissima | 2022 | 2022 Federal Survey | DEL3 | 0 | 1 | OTU-B | B | | NA | sol | | Delmarva | 38.99685 | -73.874425 | 136.9 | 14 | NA | NA | NA | 35 | | Cruise ID 202260, station 46 |
| MAB22_08024 | solidissima | 2022 | 2022 Federal Survey | DEL3 | 0 | 1 | OTU-B | B | | NA | sol | | Delmarva | 38.99685 | -73.874425 | 118.7 | 6 | NA | NA | NA | 35 | | Cruise ID 202260, station 46 |
| MAB22_09577 | solidissima | 2022 | 2022 Federal Survey | DEL3 | 0 | 1 | OTU-B | B | | NA | sol | | Delmarva | 38.661359 | -73.865009 | 82 | 4 | NA | NA | NA | 46 | | Cruise ID 202260, station 52 |
| MAB22_09590 | solidissima | 2022 | 2022 Federal Survey | DEL3 | 0 | 1 | OTU-B | B | | NA | sol | | Delmarva | 38.661359 | -73.865009 | 115.1 | 11 | NA | NA | NA | 46 | | Cruise ID 202260, station 52 |
| MAB22_09611 | solidissima | 2022 | 2022 Federal Survey | DEL3 | 0.095 | 0.905 | OTU-B | Hyb | majority B | NA | sol | | Delmarva | 38.811642 | -74.153175 | 128.6 | 14 | NA | NA | NA | 43 | | Cruise ID 202260, station 54 |
| MAB22_09612 | solidissima | 2022 | 2022 Federal Survey | DEL3 | 0 | 1 | OTU-B | B | | NA | sol | | Delmarva | 38.811642 | -74.153175 | 136.2 | 13 | NA | NA | NA | 43 | | Cruise ID 202260, station 54 |
| MAB22_09678 | solidissima | 2022 | 2022 Federal Survey | DEL3 | 0 | 1 | OTU-B | B | | NA | sol | | Delmarva | 38.811642 | -74.153175 | 112.6 | 8 | NA | NA | NA | 43 | | Cruise ID 202260, station 54 |
| MAB22_10006 | solidissima | 2022 | 2022 Federal Survey | DEL3 | 0 | 1 | OTU-B | B | | NA | sol | | Delmarva | 38.628359 | -74.290284 | 134.7 | 13 | NA | NA | NA | 39 | | Cruise ID 202260, station 56 |
| MAB22_10035 | solidissima | 2022 | 2022 Federal Survey | DEL3 | 0 | 1 | OTU-B | B | | NA | sol | | Delmarva | 38.628359 | -74.290284 | 129.7 | 15 | NA | NA | NA | 39 | | Cruise ID 202260, station 56 |
| MAB22_10037 | solidissima | 2022 | 2022 Federal Survey | DEL3 | 0 | 1 | OTU-B | B | | NA | sol | | Delmarva | 38.628359 | -74.290284 | 153.6 | 26 | NA | NA | NA | 39 | | Cruise ID 202260, station 56 |
| MAB22_10040 | solidissima | 2022 | 2022 Federal Survey | DEL3 | 0 | 1 | OTU-B | B | | NA | sol | | Delmarva | 38.628359 | -74.290284 | 143 | 18 | NA | NA | NA | 39 | | Cruise ID 202260, station 56 |
| MAB22_12173 | solidissima | 2022 | 2022 Federal Survey | DEL3 | 0 | 1 | OTU-B | B | | NA | sol | | Delmarva | 38.132384 | -74.562259 | 130.7 | 16 | NA | NA | NA | 34 | | Cruise ID 202260, station 111 |
| MAB22_12174 | solidissima | 2022 | 2022 Federal Survey | DEL3 | 0 | 1 | OTU-B | B | | NA | sol | | Delmarva | 38.132384 | -74.562259 | 91.8 | 4 | NA | NA | NA | 34 | | Cruise ID 202260, station 111 |
| MAB22_12176 | solidissima | 2022 | 2022 Federal Survey | DEL3 | 0 | 1 | OTU-B | B | | NA | sol | | Delmarva | 38.132384 | -74.562259 | 119.7 | 17 | NA | NA | NA | 34 | | Cruise ID 202260, station 111 |
| MAB22_12212 | solidissima | 2022 | 2022 Federal Survey | DEL3 | 0.038 | 0.962 | OTU-B | Hyb | majority B | NA | sol | | Delmarva | 38.132384 | -74.562259 | 127.3 | 16 | NA | NA | NA | 34 | | Cruise ID 202260, station 111 |
| MAB22_12237 | solidissima | 2022 | 2022 Federal Survey | DEL3 | 0 | 1 | OTU-B | B | | NA | sol | | Delmarva | 38.132384 | -74.562259 | 107.2 | 5 | NA | NA | NA | 34 | | Cruise ID 202260, station 111 |
| MAB22_13195 | solidissima | 2022 | 2022 Federal Survey | DEL3 | 0 | 1 | OTU-B | B | | NA | sol | | Delmarva | 38.3358 | -74.737008 | 135 | 19 | NA | NA | NA | 26 | | Cruise ID 202260, station 120 |
| MAB22_13230 | solidissima | 2022 | 2022 Federal Survey | DEL3 | 0 | 1 | OTU-B | B | | NA | sol | | Delmarva | 38.3358 | -74.737008 | 84.7 | 4 | NA | NA | NA | 26 | | Cruise ID 202260, station 120 |
| MAB22_13233 | solidissima | 2022 | 2022 Federal Survey | DEL3 | 0 | 1 | OTU-B | B | | NA | sol | | Delmarva | 38.3358 | -74.737008 | 101.1 | 5 | NA | NA | NA | 26 | | Cruise ID 202260, station 120 |
| MAB22_13238 | solidissima | 2022 | 2022 Federal Survey | DEL3 | 0 | 1 | OTU-B | B | | NA | sol | | Delmarva | 38.5304 | -74.672709 | 127.3 | 17 | NA | NA | NA | 19 | | Cruise ID 202260, station 121 |
| MAB22_13245 | solidissima | 2022 | 2022 Federal Survey | DEL3 | 0 | 1 | OTU-B | B | | NA | sol | | Delmarva | 38.5304 | -74.672709 | 88.7 | 4 | NA | NA | NA | 19 | | Cruise ID 202260, station 121 |

| sample | subspecies | collection year | Collection location or source | Analytical Group Abbreviation | STRUCTURE Q (OUT-A) | STRUCTURE Q (OTU-B) | ddRAD taxon, 10% hybrid threshold | ddRAD taxon, 1% hybrid threshold | majority taxon in hybrid | mtCOI *Sph* I | mtCOI *Bsp* 1286I | GTseq | analytical group location | Latitude | Longitude | Length (mm) | Age (yrs) | Width (mm) | Depth (mm) | weight (g) | water depth (m) | Region Avg water depth (m) | Notes |
|---|---|---|---|---|---|---|---|---|---|---|---|---|---|---|---|---|---|---|---|---|---|---|---|
| MAB22_14131 | solidissima | 2022 | 2022 Federal Survey | DEL3 | 0 | 1 | OTU-B | B | | NA | sol | | Delmarva | 38.495492 | -74.161083 | 103.8 | 11 | NA | NA | NA | 49 | | Cruise ID 202260, station 126 |
| MAB22_14148 | solidissima | 2022 | 2022 Federal Survey | DEL3 | 0 | 1 | OTU-B | B | | NA | sol | | Delmarva | 38.495492 | -74.161083 | 120.4 | 12 | NA | NA | NA | 49 | | Cruise ID 202260, station 126 |
| MAB22_14151 | solidissima | 2022 | 2022 Federal Survey | DEL3 | 0 | 1 | OTU-B | B | | NA | sol | | Delmarva | 38.495492 | -74.161083 | 111.2 | 14 | NA | NA | NA | 49 | | Cruise ID 202260, station 126 |
| MAB22_14161 | solidissima | 2022 | 2022 Federal Survey | DEL3 | 0 | 1 | OTU-B | B | | NA | sol | | Delmarva | 38.495492 | -74.161083 | 96.4 | 10 | NA | NA | NA | 49 | | Cruise ID 202260, station 126 |
| MAB22_14229 | solidissima | 2022 | 2022 Federal Survey | DEL3 | 0 | 1 | OTU-B | B | | NA | sol | | Delmarva | 38.495492 | -74.161083 | 87.4 | 5 | NA | NA | NA | 49 | | Cruise ID 202260, station 126 |
| MAB22_14461 | solidissima | 2022 | 2022 Federal Survey | DEL3 | 0 | 1 | OTU-B | B | | NA | sol | | Delmarva | 38.495492 | -74.161083 | 78.4 | 10 | NA | NA | NA | 49 | | Cruise ID 202260, station 126 |
| MAB22_14509 | solidissima | 2022 | 2022 Federal Survey | DEL3 | 0 | 1 | OTU-B | B | | NA | sol | | Delmarva | 38.48105 | -74.294592 | 139.4 | 19 | NA | NA | NA | 44 | | Cruise ID 202260, station 127 |
| MAB22_14542 | solidissima | 2022 | 2022 Federal Survey | DEL3 | 0 | 1 | OTU-B | B | | NA | sol | | Delmarva | 38.48105 | -74.294592 | 150 | 21 | NA | NA | NA | 44 | | Cruise ID 202260, station 127 |
| MAB22_14543 | solidissima | 2022 | 2022 Federal Survey | DEL3 | 0 | 1 | OTU-B | B | | NA | sol | | Delmarva | 38.48105 | -74.294592 | 125.5 | 16 | NA | NA | NA | 44 | | Cruise ID 202260, station 127 |
| MAB22_14545 | solidissima | 2022 | 2022 Federal Survey | DEL3 | 0.001 | 0.999 | OTU-B | B | | NA | sol | | Delmarva | 38.48105 | -74.294592 | 108.5 | 12 | NA | NA | NA | 44 | | Cruise ID 202260, station 127 |
| MAB22_14547 | solidissima | 2022 | 2022 Federal Survey | DEL3 | 0 | 1 | OTU-B | B | | NA | sol | | Delmarva | 38.48105 | -74.294592 | 142.4 | 18 | NA | NA | NA | 44 | | Cruise ID 202260, station 127 |
| MAB22_14557 | solidissima | 2022 | 2022 Federal Survey | DEL3 | 0 | 1 | OTU-B | B | | NA | sol | | Delmarva | 38.48105 | -74.294592 | 119.2 | 13 | NA | NA | NA | 44 | | Cruise ID 202260, station 127 |
| MAB22_14561 | solidissima | 2022 | 2022 Federal Survey | DEL3 | 0 | 1 | OTU-B | B | | NA | sol | | Delmarva | 38.48105 | -74.294592 | 93.6 | 12 | NA | NA | NA | 44 | | Cruise ID 202260, station 127 |
| MAB22_14928 | solidissima | 2022 | 2022 Federal Survey | DEL3 | 0 | 1 | OTU-B | B | | NA | sol | | Delmarva | 38.680042 | -74.632425 | 131 | 16 | NA | NA | NA | 28 | | Cruise ID 202260, station 129 |
| MAB22_14941 | solidissima | 2022 | 2022 Federal Survey | DEL3 | 0 | 1 | OTU-B | B | | NA | sol | | Delmarva | 38.680042 | -74.632425 | 96.5 | 4 | NA | NA | NA | 28 | | Cruise ID 202260, station 129 |
| MAB22_14947 | solidissima | 2022 | 2022 Federal Survey | DEL3 | 0 | 1 | OTU-B | B | | NA | sol | | Delmarva | 38.680042 | -74.632425 | 128.5 | 15 | NA | NA | NA | 28 | | Cruise ID 202260, station 129 |
| MAB22_14954 | solidissima | 2022 | 2022 Federal Survey | DEL3 | 0 | 1 | OTU-B | B | | NA | sol | | Delmarva | 38.753517 | -74.609359 | 127.5 | 10 | NA | NA | NA | 25 | | Cruise ID 202260, station 131 |
| MAB22_14955 | solidissima | 2022 | 2022 Federal Survey | DEL3 | 0 | 1 | OTU-B | B | | NA | sol | | Delmarva | 38.753517 | -74.609359 | 133.4 | 16 | NA | NA | NA | 25 | | Cruise ID 202260, station 131 |
| MAB22_14958 | solidissima | 2022 | 2022 Federal Survey | DEL3 | 0.017 | 0.983 | OTU-B | Hyb | majority B | NA | sol | | Delmarva | 38.746825 | -74.400825 | 104.4 | 12 | NA | NA | NA | 32 | | Cruise ID 202260, station 132 |
| MAB22_11856 | solidissima | 2022 | 2022 Federal Survey | DEL4 | 0 | 1 | OTU-B | B | | NA | sol | | Delmarva | 37.154192 | -75.086992 | 115.6 | 4 | NA | NA | NA | 32 | | Cruise ID 202260, station 91 |
| MAB22_11858 | solidissima | 2022 | 2022 Federal Survey | DEL4 | 0 | 1 | OTU-B | B | | NA | sol | | Delmarva | 37.154192 | -75.086992 | 111.9 | 4 | NA | NA | NA | 32 | | Cruise ID 202260, station 91 |
| MAB22_11860 | solidissima | 2022 | 2022 Federal Survey | DEL4 | 0.037 | 0.963 | OTU-B | Hyb | majority B | NA | sol | | Delmarva | 37.154192 | -75.086992 | 140.7 | 5 | NA | NA | NA | 32 | | Cruise ID 202260, station 91 |
| MAB22_11861 | solidissima | 2022 | 2022 Federal Survey | DEL4 | 0.043 | 0.957 | OTU-B | Hyb | majority B | NA | sol | | Delmarva | 37.154192 | -75.086992 | 84.9 | 3 | NA | NA | NA | 32 | | Cruise ID 202260, station 91 |
| MAB22_11863 | solidissima | 2022 | 2022 Federal Survey | DEL4 | 0 | 1 | OTU-B | B | | NA | sol | | Delmarva | 37.154192 | -75.086992 | 138.9 | 5 | NA | NA | NA | 32 | | Cruise ID 202260, station 91 |
| MAB22_11865 | solidissima | 2022 | 2022 Federal Survey | DEL4 | 0 | 1 | OTU-B | B | | NA | sol | | Delmarva | 37.154192 | -75.086992 | 124.1 | 5 | NA | NA | NA | 32 | | Cruise ID 202260, station 91 |
| MAB22_11868 | solidissima | 2022 | 2022 Federal Survey | DEL4 | 0 | 1 | OTU-B | B | | NA | sol | | Delmarva | 37.154192 | -75.086992 | 103.8 | NA | NA | NA | NA | 32 | | Cruise ID 202260, station 91 |
| MAB22_12095 | solidissima | 2022 | 2022 Federal Survey | DEL4 | 0 | 1 | OTU-B | B | | NA | sol | | Delmarva | 37.13175 | -74.9985 | 138.3 | 5 | NA | NA | NA | 38 | | Cruise ID 202260, station 92 |
| MAB22_12096 | solidissima | 2022 | 2022 Federal Survey | DEL4 | 0 | 1 | OTU-B | B | | NA | sol | | Delmarva | 37.13175 | -74.9985 | 128.9 | 5 | NA | NA | NA | 38 | | Cruise ID 202260, station 92 |
| MAB22_12113 | solidissima | 2022 | 2022 Federal Survey | DEL4 | 0 | 1 | OTU-B | B | | NA | sol | | Delmarva | 37.26705 | -74.813725 | 119.9 | 5 | NA | NA | NA | 51 | | Cruise ID 202260, station 93 |
| MAB22_12116 | solidissima | 2022 | 2022 Federal Survey | DEL4 | 0 | 1 | OTU-B | B | | NA | sol | | Delmarva | 37.376242 | -74.783859 | 100.7 | 5 | NA | NA | NA | 45 | | Cruise ID 202260, station 94 |
| MAB22_12125 | solidissima | 2022 | 2022 Federal Survey | DEL4 | 1 | 0 | OTU-A | A | | NA | sol | | Delmarva | 37.376242 | -74.783859 | 98.7 | 4 | NA | NA | NA | 45 | | Not included in EEMS or SpaceMix OTU-A analyses. Cruise ID 202260, station 94 |
| MAB22_12127 | solidissima | 2022 | 2022 Federal Survey | DEL4 | 0 | 1 | OTU-B | B | | NA | sol | | Delmarva | 37.376242 | -74.783859 | 86.8 | 3 | NA | NA | NA | 45 | | Cruise ID 202260, station 94 |
| MAB22_12130 | solidissima | 2022 | 2022 Federal Survey | DEL4 | 0 | 1 | OTU-B | B | | NA | sol | | Delmarva | 37.384959 | -74.808984 | 93 | 4 | NA | NA | NA | 47 | | Cruise ID 202260, station 95 |
| MAB22_12134 | solidissima | 2022 | 2022 Federal Survey | DEL4 | 0 | 1 | OTU-B | B | | NA | sol | | Delmarva | 37.384959 | -74.808984 | 107.1 | 4 | NA | NA | NA | 47 | | Cruise ID 202260, station 95 |
| MAB22_12494 | solidissima | 2022 | 2022 Federal Survey | DEL4 | 0 | 1 | OTU-B | B | | NA | sol | | Delmarva | 37.384959 | -74.808984 | 94.4 | 5 | NA | NA | NA | 45 | | Cruise ID 202260, station 116 |
| MAB22_12537 | solidissima | 2022 | 2022 Federal Survey | DEL4 | 0 | 1 | OTU-B | B | | NA | sol | | Delmarva | 37.384959 | -74.808984 | 110.1 | 12 | NA | NA | NA | 45 | | Cruise ID 202260, station 116 |
| MAB22_12602 | solidissima | 2022 | 2022 Federal Survey | DEL4 | 0 | 1 | OTU-B | B | | NA | sol | | Delmarva | 37.384959 | -74.808984 | 104.3 | 12 | NA | NA | NA | 45 | | Cruise ID 202260, station 116 |
| MAB22_12697 | solidissima | 2022 | 2022 Federal Survey | DEL4 | 0 | 1 | OTU-B | B | | NA | sol | | Delmarva | 37.384959 | -74.808984 | 80.1 | 4 | NA | NA | NA | 45 | | Cruise ID 202260, station 116 |
| MAB22_12705 | solidissima | 2022 | 2022 Federal Survey | DEL4 | 0 | 1 | OTU-B | B | | NA | sol | | Delmarva | 37.384959 | -74.808984 | 120.4 | 19 | NA | NA | NA | 45 | | Cruise ID 202260, station 116 |
| MAB22_10200 | solidissima | 2022 | 2022 Federal Survey | DEL5 | 0 | 1 | OTU-B | B | | NA | sol | | Delmarva | 36.620733 | -75.086892 | 85.3 | 8 | NA | NA | NA | 27 | | Cruise ID 202260, station 82 |
| MAB22_10213 | solidissima | 2022 | 2022 Federal Survey | DEL5 | 0 | 1 | OTU-B | B | | NA | sol | | Delmarva | 36.620733 | -75.086892 | 98.1 | 10 | NA | NA | NA | 27 | | Cruise ID 202260, station 82 |
| MAB22_10219 | solidissima | 2022 | 2022 Federal Survey | DEL5 | 0 | 1 | OTU-B | B | | NA | sol | | Delmarva | 36.620733 | -75.086892 | 69.5 | 9 | NA | NA | NA | 27 | | Cruise ID 202260, station 82 |
| MAB22_10220 | solidissima | 2022 | 2022 Federal Survey | DEL5 | 0 | 1 | OTU-B | B | | NA | sol | | Delmarva | 36.620733 | -75.086892 | 74.2 | 9 | NA | NA | NA | 27 | | Cruise ID 202260, station 82 |
| MAB22_10762 | solidissima | 2022 | 2022 Federal Survey | DEL5 | 0 | 1 | OTU-B | B | | NA | sol | | Delmarva | 36.608 | -75.018534 | 101 | 9 | NA | NA | NA | 26 | | Cruise ID 202260, station 83 |
| MAB22_10763 | solidissima | 2022 | 2022 Federal Survey | DEL5 | 0 | 1 | OTU-B | B | | NA | sol | | Delmarva | 36.608 | -75.018534 | 94.9 | 9 | NA | NA | NA | 26 | | Cruise ID 202260, station 83 |
| MAB22_10764 | solidissima | 2022 | 2022 Federal Survey | DEL5 | 0 | 1 | OTU-B | B | | NA | sol | | Delmarva | 36.608 | -75.018534 | 89.7 | 9 | NA | NA | NA | 26 | | Cruise ID 202260, station 83 |
| MAB22_11194 | solidissima | 2022 | 2022 Federal Survey | DEL5 | 0 | 1 | OTU-B | B | | NA | sol | | Delmarva | 36.692884 | -74.932192 | 98.2 | 8 | NA | NA | NA | 32 | | Cruise ID 202260, station 84 |
| MAB22_11233 | solidissima | 2022 | 2022 Federal Survey | DEL5 | 0.003 | 0.997 | OTU-B | B | | NA | sol | | Delmarva | 36.692884 | -74.932192 | 116.4 | 7 | NA | NA | NA | 32 | | Cruise ID 202260, station 84 |
| MAB22_11256 | solidissima | 2022 | 2022 Federal Survey | DEL5 | 0 | 1 | OTU-B | B | | NA | sol | | Delmarva | 36.692884 | -74.932192 | 108.3 | 9 | NA | NA | NA | 32 | | Cruise ID 202260, station 84 |
| MAB22_11262 | solidissima | 2022 | 2022 Federal Survey | DEL5 | 0 | 1 | OTU-B | B | | NA | sol | | Delmarva | 36.692884 | -74.932192 | 84.8 | 8 | NA | NA | NA | 32 | | Cruise ID 202260, station 84 |
| MAB22_11310 | solidissima | 2022 | 2022 Federal Survey | DEL5 | 0 | 1 | OTU-B | B | | NA | sol | | Delmarva | 36.692884 | -74.932192 | 75.8 | 8 | NA | NA | NA | 32 | | Cruise ID 202260, station 84 |
| MAB22_11533 | solidissima | 2022 | 2022 Federal Survey | DEL5 | 0 | 1 | OTU-B | B | | NA | sol | | Delmarva | 36.789492 | -74.937092 | 102.2 | 7 | NA | NA | NA | 31 | | Cruise ID 202260, station 85 |
| MAB22_11537 | solidissima | 2022 | 2022 Federal Survey | DEL5 | 0 | 1 | OTU-B | B | | NA | sol | | Delmarva | 36.789492 | -74.937092 | 113.4 | 9 | NA | NA | NA | 31 | | Cruise ID 202260, station 85 |
| MAB22_11541 | solidissima | 2022 | 2022 Federal Survey | DEL5 | 0.094 | 0.906 | OTU-B | Hyb | majority B | NA | sol | | Delmarva | 36.789492 | -74.937092 | 99.5 | 4 | NA | NA | NA | 31 | | Cruise ID 202260, station 85 |
| MAB22_11546 | solidissima | 2022 | 2022 Federal Survey | DEL5 | 0 | 1 | OTU-B | B | | NA | sol | | Delmarva | 36.789492 | -74.937092 | 121.2 | 9 | NA | NA | NA | 31 | | Cruise ID 202260, station 85 |
| MAB22_11707 | solidissima | 2022 | 2022 Federal Survey | DEL5 | 0 | 1 | OTU-B | B | | NA | sol | | Delmarva | 36.789492 | -74.937092 | 131.6 | 9 | NA | NA | NA | 31 | | Cruise ID 202260, station 85 |
| MAB22_11835 | solidissima | 2022 | 2022 Federal Survey | DEL5 | 0 | 1 | OTU-B | B | | NA | sol | | Delmarva | 36.882467 | -74.818525 | 95.4 | 8 | NA | NA | NA | 55 | | Cruise ID 202260, station 89 |
| MAB22_11838 | solidissima | 2022 | 2022 Federal Survey | DEL5 | 0 | 1 | OTU-B | B | | NA | sol | | Delmarva | 36.882467 | -74.818525 | 84.8 | 5 | NA | NA | NA | 55 | | Cruise ID 202260, station 89 |
| MAB22_11840 | solidissima | 2022 | 2022 Federal Survey | DEL5 | 0 | 1 | OTU-B | B | | NA | sol | | Delmarva | 36.882467 | -74.818525 | 119.2 | 9 | NA | NA | NA | 55 | | Cruise ID 202260, station 89 |
| GBE_171 | solidissima | 2019 | 2019 Federal Survey | GBE1 | 0 | 1 | OTU-B | B | | no amp | NA | | George's Bank | 41.6 | -66.6 | 159.1 | 14 | NA | NA | NA | NA | 56.75 | |
| GBE_172 | solidissima | 2019 | 2019 Federal Survey | GBE1 | 0 | 1 | OTU-B | B | | unreadable | NA | | George's Bank | 41.6 | -66.6 | 165.7 | 14 | NA | NA | NA | NA | | |
| GBE_182 | solidissima | 2019 | 2019 Federal Survey | GBE1 | 0 | 1 | OTU-B | B | | unreadable | NA | | George's Bank | 41.6 | -66.6 | 160.4 | 11 | NA | NA | NA | NA | | |
| GBE_183 | solidissima | 2019 | 2019 Federal Survey | GBE1 | 0 | 1 | OTU-B | B | | mtB | NA | | George's Bank | 41.6 | -66.6 | 80.6 | 3 | NA | NA | NA | NA | | |
| GBE_184 | solidissima | 2019 | 2019 Federal Survey | GBE1 | 0 | 1 | OTU-B | B | | mtB | NA | | George's Bank | 41.6 | -66.6 | 102.9 | 5 | NA | NA | NA | NA | | |
| GBE_191 | solidissima | 2019 | 2019 Federal Survey | GBE1 | 0 | 1 | OTU-B | B | | mtA | NA | | George's Bank | 41.6 | -66.6 | 163.9 | 22 | NA | NA | NA | NA | | |
| GBE_192 | solidissima | 2019 | 2019 Federal Survey | GBE1 | 0 | 1 | OTU-B | B | | mtA | NA | | George's Bank | 41.6 | -66.6 | 159.7 | 16 | NA | NA | NA | NA | | |
| GBE_193 | solidissima | 2019 | 2019 Federal Survey | GBE1 | 0 | 1 | OTU-B | B | | no amp | NA | | George's Bank | 41.6 | -66.6 | 172.7 | 27 | NA | NA | NA | NA | | |
| GBE_194 | solidissima | 2019 | 2019 Federal Survey | GBE1 | 0 | 1 | OTU-B | B | | mtA | NA | | George's Bank | 41.6 | -66.6 | 145.5 | 15 | NA | NA | NA | NA | | |
| GBE_195 | solidissima | 2019 | 2019 Federal Survey | GBE1 | 0 | 1 | OTU-B | B | | unreadable | NA | | George's Bank | 41.6 | -66.6 | 139.7 | 7 | NA | NA | NA | NA | | |
| GBE_196 | solidissima | 2019 | 2019 Federal Survey | GBE1 | 0 | 1 | OTU-B | B | | mtB | NA | | George's Bank | 41.6 | -66.6 | 182.1 | 15 | NA | NA | NA | NA | | |
| GBE_201 | solidissima | 2019 | 2019 Federal Survey | GBE1 | 0 | 1 | OTU-B | B | | mtB | NA | | George's Bank | 41.6 | -66.6 | 177.7 | 22 | NA | NA | NA | NA | | |
| GBE_203 | solidissima | 2019 | 2019 Federal Survey | GBE1 | 0 | 1 | OTU-B | B | | mtB | NA | | George's Bank | 41.6 | -66.6 | 107 | 4 | NA | NA | NA | NA | | |
| GBE_261 | solidissima | 2019 | 2019 Federal Survey | GBE1 | 0 | 1 | OTU-B | B | | no amp | NA | | George's Bank | 41.6 | -66.6 | 174.9 | 26 | NA | NA | NA | NA | | |
| GBE_262 | solidissima | 2019 | 2019 Federal Survey | GBE1 | 0 | 1 | OTU-B | B | | unreadable | NA | | George's Bank | 41.6 | -66.6 | 171.4 | 26 | NA | NA | NA | NA | | |
| GBE_263 | solidissima | 2019 | 2019 Federal Survey | GBE1 | 0 | 1 | OTU-B | B | | mtB | NA | | George's Bank | 41.6 | -66.6 | 167 | 13 | NA | NA | NA | NA | | |
| GBE_291 | solidissima | 2019 | 2019 Federal Survey | GBE1 | 0 | 1 | OTU-B | B | | unreadable | NA | | George's Bank | 41.6 | -66.6 | 169.6 | 16 | NA | NA | NA | NA | | |
| GBE_332 | solidissima | 2019 | 2019 Federal Survey | GBE1 | 0 | 1 | OTU-B | B | | mtB | NA | | George's Bank | 41.6 | -66.6 | 137.6 | 16 | NA | NA | NA | NA | | |
| GBE_333 | solidissima | 2019 | 2019 Federal Survey | GBE1 | 0 | 1 | OTU-B | B | | unreadable | NA | | George's Bank | 41.6 | -66.6 | 86.9 | 4 | NA | NA | NA | NA | | |
| GBE_334 | solidissima | 2019 | 2019 Federal Survey | GBE1 | 0 | 1 | OTU-B | B | | mtB | NA | | George's Bank | 41.6 | -66.6 | 98.6 | 4 | NA | NA | NA | NA | | |
| GBE_335 | solidissima | 2019 | 2019 Federal Survey | GBE1 | 0 | 1 | OTU-B | B | | no amp | NA | | George's Bank | 41.6 | -66.6 | 129.6 | 13 | NA | NA | NA | NA | | |
| GBE_336 | solidissima | 2019 | 2019 Federal Survey | GBE1 | 0 | 1 | OTU-B | B | | mtB | NA | | George's Bank | 41.6 | -66.6 | 152.4 | 21 | NA | NA | NA | NA | | |
| GBE_371 | solidissima | 2019 | 2019 Federal Survey | GBE1 | 0 | 1 | OTU-B | B | | mtB | NA | | George's Bank | 41.6 | -66.6 | 146 | 8 | NA | NA | NA | NA | | |
| GBE_373 | solidissima | 2019 | 2019 Federal Survey | GBE1 | 0 | 1 | OTU-B | B | | mtB | NA | | George's Bank | 41.6 | -66.6 | 75.8 | 3 | NA | NA | NA | NA | | |
| GBE_381 | solidissima | 2019 | 2019 Federal Survey | GBE1 | 0 | 1 | OTU-B | B | | mtB | NA | | George's Bank | 41.6 | -66.6 | 127.3 | 22 | NA | NA | NA | NA | | |
| GBE_382 | solidissima | 2019 | 2019 Federal Survey | GBE1 | 0 | 1 | OTU-B | B | | no amp | NA | | George's Bank | 41.6 | -66.6 | 140.9 | 23 | NA | NA | NA | NA | | |
| GBE_383 | solidissima | 2019 | 2019 Federal Survey | GBE1 | 0 | 1 | OTU-B | B | | mtB | NA | | George's Bank | 41.6 | -66.6 | 135.1 | 25 | NA | NA | NA | NA | | |
| GBE_384 | solidissima | 2019 | 2019 Federal Survey | GBE1 | 0 | 1 | OTU-B | B | | no amp | NA | | George's Bank | 41.6 | -66.6 | 138.6 | 9 | NA | NA | NA | NA | | |
| GBE_385 | solidissima | 2019 | 2019 Federal Survey | GBE1 | 0 | 1 | OTU-B | B | | mtB | NA | | George's Bank | 41.6 | -66.6 | 115.8 | 6 | NA | NA | NA | NA | | |
| GBE_401 | solidissima | 2019 | 2019 Federal Survey | GBE1 | 0 | 1 | OTU-B | B | | mtA | NA | | George's Bank | 41.6 | -66.6 | 115.9 | 15 | NA | NA | NA | NA | | |
| GBE_402 | solidissima | 2019 | 2019 Federal Survey | GBE1 | 0 | 1 | OTU-B | B | | mtB | NA | | George's Bank | 41.6 | -66.6 | 109.8 | 15 | NA | NA | NA | NA | | |
| GBE_441 | solidissima | 2019 | 2019 Federal Survey | GBE1 | 0 | 1 | OTU-B | B | | mtB | NA | | George's Bank | 41.6 | -66.6 | 143.3 | 22 | NA | NA | NA | NA | | |
| GBE_442 | solidissima | 2019 | 2019 Federal Survey | GBE1 | 0 | 1 | OTU-B | B | | mtB | NA | | George's Bank | 41.6 | -66.6 | 139.7 | 17 | NA | NA | NA | NA | | |
| GBE_443 | solidissima | 2019 | 2019 Federal Survey | GBE1 | 0 | 1 | OTU-B | B | | mtB | NA | | George's Bank | 41.6 | -66.6 | 124.7 | 8 | NA | NA | NA | NA | | |
| GBE_471 | solidissima | 2019 | 2019 Federal Survey | GBE1 | 0 | 1 | OTU-B | B | | no amp | NA | | George's Bank | 41.6 | -66.6 | 147.6 | 9 | NA | NA | NA | NA | | |
| GBE_472 | solidissima | 2019 | 2019 Federal Survey | GBE1 | 0 | 1 | OTU-B | B | | no amp | NA | | George's Bank | 41.6 | -66.6 | 134.8 | 7 | NA | NA | NA | NA | | |
| GBE_473 | solidissima | 2019 | 2019 Federal Survey | GBE1 | 0 | 1 | OTU-B | B | | no amp | NA | | George's Bank | 41.6 | -66.6 | 150.6 | 9 | NA | NA | NA | NA | | |
| GBE_481 | solidissima | 2019 | 2019 Federal Survey | GBE1 | 0 | 1 | OTU-B | B | | no amp | NA | | George's Bank | 41.6 | -66.6 | 151.1 | 8 | NA | NA | NA | NA | | |
| GBE_483 | solidissima | 2019 | 2019 Federal Survey | GBE1 | 0 | 1 | OTU-B | B | | no amp | NA | | George's Bank | 41.6 | -66.6 | 163.9 | 23 | NA | NA | NA | NA | | |
| GBE_484 | solidissima | 2019 | 2019 Federal Survey | GBE1 | 0 | 1 | OTU-B | B | | no amp | NA | | George's Bank | 41.6 | -66.6 | 180.3 | 22 | NA | NA | NA | NA | | |
| GBE_521 | solidissima | 2019 | 2019 Federal Survey | GBE1 | 0.008 | 0.992 | OTU-B | B | | no amp | NA | | George's Bank | 41.6 | -66.6 | 196.5 | 21 | NA | NA | NA | NA | | |
| GBE_522 | solidissima | 2019 | 2019 Federal Survey | GBE1 | 0 | 1 | OTU-B | B | | mtB | NA | | George's Bank | 41.6 | -66.6 | 149 | 21 | NA | NA | NA | NA | | |
| GBE_523 | solidissima | 2019 | 2019 Federal Survey | GBE1 | 0 | 1 | OTU-B | B | | mtB | NA | | George's Bank | 41.6 | -66.6 | 152.4 | 8 | NA | NA | NA | NA | | |
| 536_001 | solidissima | 1999 | 1999 Federal Survey | GBE2 | 0 | 1 | OTU-B | B | | mtB | sol | | George's Bank | 41.5498 | -67.3732 | 134 | 6 | NA | NA | NA | NA | | |
| 536_003 | solidissima | 1999 | 1999 Federal Survey | GBE2 | 0 | 1 | OTU-B | B | | mtB | sol | | George's Bank | 41.5498 | -67.3732 | 109 | 5 | NA | NA | NA | NA | | |
| 536_004 | solidissima | 1999 | 1999 Federal Survey | GBE2 | 0 | 1 | OTU-B | B | | mtA | sol | | George's Bank | 41.5498 | -67.3732 | 132 | 4 | NA | NA | NA | NA | | |
| 536_005 | solidissima | 1999 | 1999 Federal Survey | GBE2 | 0 | 1 | OTU-B | B | | mtB | sol | | George's Bank | 41.5498 | -67.3732 | 114 | 3 | NA | NA | NA | NA | | |
| 536_006 | solidissima | 1999 | 1999 Federal Survey | GBE2 | 0 | 1 | OTU-B | B | | mtB | sol | | George's Bank | 41.5498 | -67.3732 | 133 | 5 | NA | NA | NA | NA | | |
| 536_007 | solidissima | 1999 | 1999 Federal Survey | GBE2 | 0 | 1 | OTU-B | B | | mtB | sol | | George's Bank | 41.5498 | -67.3732 | 119 | 4 | NA | NA | NA | NA | | |
| 536_008 | solidissima | 1999 | 1999 Federal Survey | GBE2 | 0 | 1 | OTU-B | B | | mtB | sol | | George's Bank | 41.5498 | -67.3732 | 109 | 5 | NA | NA | NA | NA | | |
| 536_009 | solidissima | 1999 | 1999 Federal Survey | GBE2 | 0 | 1 | OTU-B | B | | mtB | sol | | George's Bank | 41.5498 | -67.3732 | 119 | 4 | NA | NA | NA | NA | | |
| 536_010 | solidissima | 1999 | 1999 Federal Survey | GBE2 | 0 | 1 | OTU-B | B | | mtB | sol | | George's Bank | 41.5498 | -67.3732 | 122 | 5 | NA | NA | NA | NA | | |
| 536_011 | solidissima | 1999 | 1999 Federal Survey | GBE2 | 0 | 1 | OTU-B | B | | mtB | sol | | George's Bank | 41.5498 | -67.3732 | 133 | 6 | NA | NA | NA | NA | | |
| 536_012 | solidissima | 1999 | 1999 Federal Survey | GBE2 | 0 | 1 | OTU-B | B | | mtB | sol | | George's Bank | 41.5498 | -67.3732 | 103 | 3 | NA | NA | NA | NA | | |
| 536_013 | solidissima | 1999 | 1999 Federal Survey | GBE2 | 0 | 1 | OTU-B | B | | mtB | sol | | George's Bank | 41.5498 | -67.3732 | 122 | 3 | NA | NA | NA | NA | | |
| 536_014 | solidissima | 1999 | 1999 Federal Survey | GBE2 | 0 | 1 | OTU-B | B | | mtB | sol | | George's Bank | 41.5498 | -67.3732 | 104 | 3 | NA | NA | NA | NA | | |
| 536_015 | solidissima | 1999 | 1999 Federal Survey | GBE2 | 0 | 1 | OTU-B | B | | mtB | sol | | George's Bank | 41.5498 | -67.3732 | 133 | 5 | NA | NA | NA | NA | | |
| 536_016 | solidissima | 1999 | 1999 Federal Survey | GBE2 | 0 | 1 | OTU-B | B | | mtB | sol | | George's Bank | 41.5498 | -67.3732 | 128 | 5 | NA | NA | NA | NA | | |
| 536_017 | solidissima | 1999 | 1999 Federal Survey | GBE2 | 0 | 1 | OTU-B | B | | mtB | sol | | George's Bank | 41.5498 | -67.3732 | 89 | 3 | NA | NA | NA | NA | | |
| 536_018 | solidissima | 1999 | 1999 Federal Survey | GBE2 | 0 | 1 | OTU-B | B | | mtA | sol | | George's Bank | 41.5498 | -67.3732 | 91 | 3 | NA | NA | NA | NA | | |
| 536_019 | solidissima | 1999 | 1999 Federal Survey | GBE2 | 0 | 1 | OTU-B | B | | mtB | sol | | George's Bank | 41.5498 | -67.3732 | 117 | 5 | NA | NA | NA | NA | | |
| 536_020 | solidissima | 1999 | 1999 Federal Survey | GBE2 | 0 | 1 | OTU-B | B | | mtB | sol | | George's Bank | 41.5498 | -67.3732 | 146 | 6 | NA | NA | NA | NA | | |
| 536_021 | solidissima | 1999 | 1999 Federal Survey | GBE2 | 0 | 1 | OTU-B | B | | mtB | sol | | George's Bank | 41.5498 | -67.3732 | 142 | 6 | NA | NA | NA | NA | | |
| 536_022 | solidissima | 1999 | 1999 Federal Survey | GBE2 | 0 | 1 | OTU-B | B | | mtB | sol | | George's Bank | 41.5498 | -67.3732 | 143 | 6 | NA | NA | NA | NA | | |
| 536_024 | solidissima | 1999 | 1999 Federal Survey | GBE2 | 0 | 1 | OTU-B | B | | mtB | sol | | George's Bank | 41.5498 | -67.3732 | 92 | 3 | NA | NA | NA | NA | | |
| 536_025 | solidissima | 1999 | 1999 Federal Survey | GBE2 | 0 | 1 | OTU-B | B | | mtB | sol | | George's Bank | 41.5498 | -67.3732 | 143 | 6 | NA | NA | NA | NA | | |
| 536_026 | solidissima | 1999 | 1999 Federal Survey | GBE2 | 0 | 1 | OTU-B | B | | mtB | sol | | George's Bank | 41.5498 | -67.3732 | 113 | 4 | NA | NA | NA | NA | | |
| 536_028 | solidissima | 1999 | 1999 Federal Survey | GBE2 | 0 | 1 | OTU-B | B | | mtB | sol | | George's Bank | 41.5498 | -67.3732 | 77 | 3 | NA | NA | NA | NA | | |
| 536_029 | solidissima | 1999 | 1999 Federal Survey | GBE2 | 0 | 1 | OTU-B | B | | mtB | sol | | George's Bank | 41.5498 | -67.3732 | 90 | 3 | NA | NA | NA | NA | | |
| NAN_001 | solidissima | 2020 | 2020 Federal Sample | NAT | 0 | 1 | OTU-B | B | | mtB | sol | | Nantucket Shoals | 40.818333 | -70.056667 | 152 | NA | NA | NA | NA | 20 | 20 | |
| NAN_002 | solidissima | 2020 | 2020 Federal Sample | NAT | 0.008 | 0.992 | OTU-B | B | | mtB | sol | | Nantucket Shoals | 40.818333 | -70.056667 | 169 | NA | NA | NA | NA | 20 | | |
| NAN_003 | solidissima | 2020 | 2020 Federal Sample | NAT | 0 | 1 | OTU-B | B | | no amp | no amp | | Nantucket Shoals | 40.818333 | -70.056667 | 172 | NA | NA | NA | NA | 20 | | |
| NAN_007 | solidissima | 2020 | 2020 Federal Sample | NAT | 0 | 1 | OTU-B | B | | mtB | sol | | Nantucket Shoals | 40.818333 | -70.056667 | 171 | NA | NA | NA | NA | 20 | | |
| NAN_008 | solidissima | 2020 | 2020 Federal Sample | NAT | 0 | 1 | OTU-B | B | | mtB | sol | | Nantucket Shoals | 40.818333 | -70.056667 | 161 | NA | NA | NA | NA | 20 | | |
| NAN_009 | solidissima | 2020 | 2020 Federal Sample | NAT | 0 | 1 | OTU-B | B | | mtA | sol | | Nantucket Shoals | 40.818333 | -70.056667 | 147 | NA | NA | NA | NA | 20 | | |
| NAN_010 | solidissima | 2020 | 2020 Federal Sample | NAT | 0 | 1 | OTU-B | B | | mtB | sol | | Nantucket Shoals | 40.818333 | -70.056667 | 152 | NA | NA | NA | NA | 20 | | |
| NAN_012 | solidissima | 2020 | 2020 Federal Sample | NAT | 0 | 1 | OTU-B | B | | mtB | sol | | Nantucket Shoals | 40.818333 | -70.056667 | 164 | NA | NA | NA | NA | 20 | | |
| NAN_014 | solidissima | 2020 | 2020 Federal Sample | NAT | 0 | 1 | OTU-B | B | | mtB | sol | | Nantucket Shoals | 40.818333 | -70.056667 | 173 | NA | NA | NA | NA | 20 | | |
| NAN_019 | solidissima | 2020 | 2020 Federal Sample | NAT | 0 | 1 | OTU-B | B | | mtA | sol | | Nantucket Shoals | 40.818333 | -70.056667 | 175 | NA | NA | NA | NA | 20 | | |
| NAN_021 | solidissima | 2020 | 2020 Federal Sample | NAT | 0 | 1 | OTU-B | B | | no amp | no amp | | Nantucket Shoals | 40.818333 | -70.056667 | 160 | NA | NA | NA | NA | 20 | | |
| NAN_022 | solidissima | 2020 | 2020 Federal Sample | NAT | 0 | 1 | OTU-B | B | | no amp | no amp | | Nantucket Shoals | 40.818333 | -70.056667 | 163 | NA | NA | NA | NA | 20 | | |
| NAN_023 | solidissima | 2020 | 2020 Federal Sample | NAT | 0 | 1 | OTU-B | B | | mtB | sol | | Nantucket Shoals | 40.818333 | -70.056667 | 154 | NA | NA | NA | NA | 20 | | |
| NAN_024 | solidissima | 2020 | 2020 Federal Sample | NAT | 0 | 1 | OTU-B | B | | mtA | sol | | Nantucket Shoals | 40.818333 | -70.056667 | 165 | NA | NA | NA | NA | 20 | | |
| NAN_025 | solidissima | 2020 | 2020 Federal Sample | NAT | 0 | 1 | OTU-B | B | | mtB | sol | | Nantucket Shoals | 40.818333 | -70.056667 | 154 | NA | NA | NA | NA | 20 | | |
| NAN_026 | solidissima | 2020 | 2020 Federal Sample | NAT | 0 | 1 | OTU-B | B | | no amp | no amp | | Nantucket Shoals | 40.818333 | -70.056667 | 162 | NA | NA | NA | NA | 20 | | |
| NAN_028 | solidissima | 2020 | 2020 Federal Sample | NAT | 0 | 1 | OTU-B | B | | no amp | no amp | | Nantucket Shoals | 40.818333 | -70.056667 | 153 | NA | NA | NA | NA | 20 | | |
| NAN_031 | solidissima | 2020 | 2020 Federal Sample | NAT | 0.001 | 0.999 | OTU-B | B | | no amp | no amp | | Nantucket Shoals | 40.818333 | -70.056667 | 152 | NA | NA | NA | NA | 20 | | |
| NAN_033 | solidissima | 2020 | 2020 Federal Sample | NAT | 0 | 1 | OTU-B | B | | no amp | no amp | | Nantucket Shoals | 40.818333 | -70.056667 | 157 | NA | NA | NA | NA | 20 | | |
| NAN_034 | solidissima | 2020 | 2020 Federal Sample | NAT | 0 | 1 | OTU-B | B | | mtB | sol | | Nantucket Shoals | 40.818333 | -70.056667 | 163 | NA | NA | NA | NA | 20 | | |
| NAN_036 | solidissima | 2020 | 2020 Federal Sample | NAT | 0 | 1 | OTU-B | B | | no amp | no amp | | Nantucket Shoals | 40.818333 | -70.056667 | 158 | NA | NA | NA | NA | 20 | | |
| NAN_038 | solidissima | 2020 | 2020 Federal Sample | NAT | 0 | 1 | OTU-B | B | | no amp | no amp | | Nantucket Shoals | 40.818333 | -70.056667 | 161 | NA | NA | NA | NA | 20 | | |
| NAN_039 | solidissima | 2020 | 2020 Federal Sample | NAT | 0 | 1 | OTU-B | B | | no amp | no amp | | Nantucket Shoals | 40.818333 | -70.056667 | 162 | NA | NA | NA | NA | 20 | | |
| NAN_041 | solidissima | 2020 | 2020 Federal Sample | NAT | 0 | 1 | OTU-B | B | | no amp | no amp | | Nantucket Shoals | 40.818333 | -70.056667 | 160 | NA | NA | NA | NA | 20 | | |
| NAN_042 | solidissima | 2020 | 2020 Federal Sample | NAT | 0 | 1 | OTU-B | B | | no amp | no amp | | Nantucket Shoals | 40.818333 | -70.056667 | 170 | NA | NA | NA | NA | 20 | | |
| NAN_045 | solidissima | 2020 | 2020 Federal Sample | NAT | 0 | 1 | OTU-B | B | | mtB | sol | | Nantucket Shoals | 40.818333 | -70.056667 | 149 | NA | NA | NA | NA | 20 | | |
| NAN_046 | solidissima | 2020 | 2020 Federal Sample | NAT | 0 | 1 | OTU-B | B | | no amp | no amp | | Nantucket Shoals | 40.818333 | -70.056667 | 172 | NA | NA | NA | NA | 20 | | |
| NAN_048 | solidissima | 2020 | 2020 Federal Sample | NAT | 0 | 1 | OTU-B | B | | no amp | no amp | | Nantucket Shoals | 40.818333 | -70.056667 | 150 | NA | NA | NA | NA | 20 | | |
| NAN_049 | solidissima | 2020 | 2020 Federal Sample | NAT | 0 | 1 | OTU-B | B | | mtB | sol | | Nantucket Shoals | 40.818333 | -70.056667 | 149 | NA | NA | NA | NA | 20 | | |
| NAN_050 | solidissima | 2020 | 2020 Federal Sample | NAT | 0 | 1 | OTU-B | B | | mtB | sol | | Nantucket Shoals | 40.818333 | -70.056667 | 178 | NA | NA | NA | NA | 20 | | |
| MAB22_04410 | solidissima | 2022 | 2022 Federal Survey | NJ1 | 0 | 1 | OTU-B | B | | NA | sol | | New Jersey | 40.04935 | -73.821834 | 121.5 | 2 | NA | NA | NA | 25 | 32.4 | Cruise ID 202260, station 30 |
| MAB22_15806 | solidissima | 2022 | 2022 Federal Survey | NJ1 | 0 | 1 | OTU-B | B | | NA | sol | | New Jersey | 40.136817 | -73.262458 | 123.5 | 10 | NA | NA | NA | 39 | | Cruise ID 202260, station 160 |
| MAB22_17243 | solidissima | 2022 | 2022 Federal Survey | NJ1 | 0 | 1 | OTU-B | B | | NA | sol | | New Jersey | 40.270258 | -72.949092 | 122.1 | 13 | NA | NA | NA | 42 | | Cruise ID 202260, station 165 |
| MAB22_17246 | solidissima | 2022 | 2022 Federal Survey | NJ1 | 0 | 1 | OTU-B | B | | NA | sol | | New Jersey | 40.270258 | -72.949092 | 97.5 | 10 | NA | NA | NA | 42 | | Cruise ID 202260, station 165 |
| MAB22_17620 | solidissima | 2022 | 2022 Federal Survey | NJ1 | 0 | 1 | OTU-B | B | | NA | sol | | New Jersey | 40.389175 | -73.065584 | 116.3 | 11 | NA | NA | NA | 36 | | Cruise ID 202260, station 167 |
| MAB22_17626 | solidissima | 2022 | 2022 Federal Survey | NJ1 | 0 | 1 | OTU-B | B | | NA | sol | | New Jersey | 40.389175 | -73.065584 | 120.9 | 9 | NA | NA | NA | 36 | | Cruise ID 202260, station 167 |
| MAB22_17650 | solidissima | 2022 | 2022 Federal Survey | NJ1 | 0 | 1 | OTU-B | B | | NA | sol | | New Jersey | 40.389175 | -73.065584 | 134.5 | 9 | NA | NA | NA | 36 | | Cruise ID 202260, station 167 |
| MAB22_17672 | solidissima | 2022 | 2022 Federal Survey | NJ1 | 0 | 1 | OTU-B | B | | NA | sol | | New Jersey | 40.389175 | -73.065584 | 93.8 | 5 | NA | NA | NA | 36 | | Cruise ID 202260, station 167 |
| MAB22_18643 | solidissima | 2022 | 2022 Federal Survey | NJ1 | 0 | 1 | OTU-B | B | | NA | sol | | New Jersey | 40.475409 | -73.325275 | 177.2 | 21 | NA | NA | NA | 21 | | Cruise ID 202260, station 173 |
| MAB22_18704 | solidissima | 2022 | 2022 Federal Survey | NJ1 | 0 | 1 | OTU-B | B | | NA | sol | | New Jersey | 40.475409 | -73.325275 | 169.5 | 20 | NA | NA | NA | 21 | | Cruise ID 202260, station 173 |
| MAB22_18737 | solidissima | 2022 | 2022 Federal Survey | NJ1 | 0 | 1 | OTU-B | B | | NA | sol | | New Jersey | 40.475409 | -73.325275 | 139.4 | 10 | NA | NA | NA | 21 | | Cruise ID 202260, station 173 |
| MAB22_18760 | solidissima | 2022 | 2022 Federal Survey | NJ1 | 0 | 1 | OTU-B | B | | NA | sol | | New Jersey | 40.475409 | -73.325275 | 98.8 | 4 | NA | NA | NA | 21 | | Cruise ID 202260, station 173 |
| MAB22_18763 | solidissima | 2022 | 2022 Federal Survey | NJ1 | 0 | 1 | OTU-B | B | | NA | sol | | New Jersey | 40.475409 | -73.325275 | 118.9 | 5 | NA | NA | NA | 21 | | Cruise ID 202260, station 173 |
| MAB22_18774 | solidissima | 2022 | 2022 Federal Survey | NJ1 | 0 | 1 | OTU-B | B | | NA | sol | | New Jersey | 40.475409 | -73.325275 | 120.4 | 6 | NA | NA | NA | 21 | | Cruise ID 202260, station 173 |
| MAB22_18793 | solidissima | 2022 | 2022 Federal Survey | NJ1 | 0 | 1 | OTU-B | B | | NA | sol | | New Jersey | 40.475409 | -73.325275 | 108.2 | 4 | NA | NA | NA | 21 | | Cruise ID 202260, station 173 |
| MAB22_02638 | solidissima | 2022 | 2022 Federal Survey | NJ2 | 0 | 1 | OTU-B | B | | NA | sol | | New Jersey | 39.295 | -73.420425 | 111.6 | 8 | NA | NA | NA | 44 | | Cruise ID 202260, station 20 |
| MAB22_02757 | solidissima | 2022 | 2022 Federal Survey | NJ2 | 0 | 1 | OTU-B | B | | NA | sol | | New Jersey | 39.295 | -73.420425 | 76.9 | 6 | NA | NA | NA | 44 | | Cruise ID 202260, station 20 |
| MAB22_02822 | solidissima | 2022 | 2022 Federal Survey | NJ2 | 0 | 1 | OTU-B | B | | NA | sol | | New Jersey | 39.295 | -73.420425 | 99.5 | NA | NA | NA | NA | 44 | | Cruise ID 202260, station 20 |
| MAB22_02889 | solidissima | 2022 | 2022 Federal Survey | NJ2 | 0 | 1 | OTU-B | B | | NA | sol | | New Jersey | 39.295 | -73.420425 | 83.8 | 4 | NA | NA | NA | 44 | | Cruise ID 202260, station 20 |
| MAB22_02957 | solidissima | 2022 | 2022 Federal Survey | NJ2 | 0 | 1 | OTU-B | B | | NA | sol | | New Jersey | 39.295 | -73.420425 | 125.6 | 7 | NA | NA | NA | 44 | | Cruise ID 202260, station 20 |
| MAB22_02959 | solidissima | 2022 | 2022 Federal Survey | NJ2 | 0 | 1 | OTU-B | B | | NA | sol | | New Jersey | 39.295 | -73.420425 | 133.8 | 14 | NA | NA | NA | 44 | | Cruise ID 202260, station 20 |
| MAB22_03373 | solidissima | 2022 | 2022 Federal Survey | NJ2 | 0 | 1 | OTU-B | B | | NA | sol | | New Jersey | 39.525883 | -73.780033 | 152.5 | 26 | NA | NA | NA | 24 | | Cruise ID 202260, station 23 |
| MAB22_03483 | solidissima | 2022 | 2022 Federal Survey | NJ2 | 0.057 | 0.943 | OTU-B | Hyb | majority B | NA | sol | | New Jersey | 39.525883 | -73.780033 | 136.2 | 7 | NA | NA | NA | 24 | | Cruise ID 202260, station 23 |
| MAB22_03484 | solidissima | 2022 | 2022 Federal Survey | NJ2 | 0 | 1 | OTU-B | B | | NA | sol | | New Jersey | 39.525883 | -73.780033 | 108.2 | 4 | NA | NA | NA | 24 | | Cruise ID 202260, station 23 |
| MAB22_03512 | solidissima | 2022 | 2022 Federal Survey | NJ2 | 0 | 1 | OTU-B | B | | NA | sol | | New Jersey | 39.564858 | -73.758 | 147.3 | 11 | NA | NA | NA | 24 | | Cruise ID 202260, station 24 |
| MAB22_03590 | solidissima | 2022 | 2022 Federal Survey | NJ2 | 0 | 1 | OTU-B | B | | NA | sol | | New Jersey | 39.564858 | -73.758 | 160 | 12 | NA | NA | NA | 24 | | Cruise ID 202260, station 24 |
| MAB22_03597 | solidissima | 2022 | 2022 Federal Survey | NJ2 | 0.016 | 0.984 | OTU-B | Hyb | majority B | NA | sol | | New Jersey | 39.564858 | -73.758 | 123.5 | 4 | NA | NA | NA | 24 | | Cruise ID 202260, station 24 |
| MAB22_03604 | solidissima | 2022 | 2022 Federal Survey | NJ2 | 0 | 1 | OTU-B | B | | NA | sol | | New Jersey | 39.538483 | -73.703284 | 152.5 | 28 | NA | NA | NA | 26 | | Cruise ID 202260, station 25 |
| MAB22_03687 | solidissima | 2022 | 2022 Federal Survey | NJ2 | 0 | 1 | OTU-B | B | | NA | sol | | New Jersey | 39.538483 | -73.703284 | 129.9 | 5 | NA | NA | NA | 26 | | Cruise ID 202260, station 25 |
| MAB22_03707 | solidissima | 2022 | 2022 Federal Survey | NJ2 | 0 | 1 | OTU-B | B | | NA | sol | | New Jersey | 39.538483 | -73.703284 | 162.9 | 13 | NA | NA | NA | 26 | | Cruise ID 202260, station 25 |
| MAB22_03736 | solidissima | 2022 | 2022 Federal Survey | NJ2 | 0 | 1 | OTU-B | B | | NA | sol | | New Jersey | 39.558375 | -73.67875 | 162 | 18 | NA | NA | NA | 28 | | Cruise ID 202260, station 26 |
| MAB22_03765 | solidissima | 2022 | 2022 Federal Survey | NJ2 | 0 | 1 | OTU-B | B | | NA | sol | | New Jersey | 39.558375 | -73.67875 | 147.1 | 25 | NA | NA | NA | 28 | | Cruise ID 202260, station 26 |
| MAB22_03824 | solidissima | 2022 | 2022 Federal Survey | NJ2 | 0.001 | 0.999 | OTU-B | B | | NA | sol | | New Jersey | 39.558375 | -73.67875 | 130.1 | 8 | NA | NA | NA | 28 | | Cruise ID 202260, station 26 |
| MAB22_04427 | solidissima | 2022 | 2022 Federal Survey | NJ2 | 0 | 1 | OTU-B | B | | NA | sol | | New Jersey | 39.997242 | -73.534817 | 137 | 24 | NA | NA | NA | 37 | | Cruise ID 202260, station 32 |
| MAB22_04444 | solidissima | 2022 | 2022 Federal Survey | NJ2 | 0 | 1 | OTU-B | B | | NA | sol | | New Jersey | 39.997242 | -73.534817 | 103.6 | 8 | NA | NA | NA | 37 | | Cruise ID 202260, station 32 |
| MAB22_04450 | solidissima | 2022 | 2022 Federal Survey | NJ2 | 0.005 | 0.995 | OTU-B | B | | NA | sol | | New Jersey | 39.997242 | -73.534817 | 116 | 12 | NA | NA | NA | 37 | | Cruise ID 202260, station 32 |
| MAB22_04536 | solidissima | 2022 | 2022 Federal Survey | NJ2 | 0 | 1 | OTU-B | B | | NA | sol | | New Jersey | 39.997242 | -73.534817 | 79.1 | 5 | NA | NA | NA | 37 | | Cruise ID 202260, station 32 |
| MAB22_04559 | solidissima | 2022 | 2022 Federal Survey | NJ2 | 0 | 1 | OTU-B | B | | NA | sol | | New Jersey | 39.927875 | -73.56035 | 136.3 | 24 | NA | NA | NA | 29 | | Cruise ID 202260, station 33 |
| MAB22_04563 | solidissima | 2022 | 2022 Federal Survey | NJ2 | 0 | 1 | OTU-B | B | | NA | sol | | New Jersey | 39.927875 | -73.56035 | 115.1 | 9 | NA | NA | NA | 29 | | Cruise ID 202260, station 33 |
| MAB22_04603 | solidissima | 2022 | 2022 Federal Survey | NJ2 | 0 | 1 | OTU-B | B | | NA | sol | | New Jersey | 39.927875 | -73.56035 | 128.8 | 17 | NA | NA | NA | 29 | | Cruise ID 202260, station 33 |
| MAB22_04648 | solidissima | 2022 | 2022 Federal Survey | NJ2 | 0.001 | 0.999 | OTU-B | B | | NA | sol | | New Jersey | 39.927875 | -73.56035 | 91.3 | 6 | NA | NA | NA | 29 | | Cruise ID 202260, station 33 |
| MAB22_04655 | solidissima | 2022 | 2022 Federal Survey | NJ2 | 0 | 1 | OTU-B | B | | NA | sol | | New Jersey | 39.927875 | -73.56035 | 106.6 | 6 | NA | NA | NA | 29 | | Cruise ID 202260, station 33 |
| MAB22_05392 | solidissima | 2022 | 2022 Federal Survey | NJ2 | 0 | 1 | OTU-B | B | | NA | sol | | New Jersey | 39.644167 | -73.612067 | 157 | 21 | NA | NA | NA | 32 | | Cruise ID 202260, station 36 |
| MAB22_05397 | solidissima | 2022 | 2022 Federal Survey | NJ2 | 0 | 1 | OTU-B | B | | NA | sol | | New Jersey | 39.644167 | -73.612067 | 97.6 | 4 | NA | NA | NA | 32 | | Cruise ID 202260, station 36 |
| MAB22_05467 | solidissima | 2022 | 2022 Federal Survey | NJ2 | 0 | 1 | OTU-B | B | | NA | sol | | New Jersey | 39.644167 | -73.612067 | 168.4 | 23 | NA | NA | NA | 32 | | Cruise ID 202260, station 36 |
| MAB22_05482 | solidissima | 2022 | 2022 Federal Survey | NJ2 | 0 | 1 | OTU-B | B | | NA | sol | | New Jersey | 39.644167 | -73.612067 | 114.5 | 5 | NA | NA | NA | 32 | | Cruise ID 202260, station 36 |
| MAB22_05490 | solidissima | 2022 | 2022 Federal Survey | NJ2 | 0 | 1 | OTU-B | B | | NA | sol | | New Jersey | 39.644167 | -73.612067 | 89 | 3 | NA | NA | NA | 32 | | Cruise ID 202260, station 36 |
| MAB22_05661 | solidissima | 2022 | 2022 Federal Survey | NJ2 | 0 | 1 | OTU-B | B | | NA | sol | | New Jersey | 39.454159 | -73.609625 | 150.6 | 15 | NA | NA | NA | 31 | | Cruise ID 202260, station 38 |
| MAB22_05763 | solidissima | 2022 | 2022 Federal Survey | NJ2 | 0 | 1 | OTU-B | B | | NA | sol | | New Jersey | 39.454159 | -73.609625 | 127.3 | 6 | NA | NA | NA | 31 | | Cruise ID 202260, station 38 |
| MAB22_05788 | solidissima | 2022 | 2022 Federal Survey | NJ2 | 0 | 1 | OTU-B | B | | NA | sol | | New Jersey | 39.454159 | -73.609625 | 133.7 | 13 | NA | NA | NA | 31 | | Cruise ID 202260, station 38 |
| MAB22_05839 | solidissima | 2022 | 2022 Federal Survey | NJ2 | 0 | 1 | OTU-B | B | | NA | sol | | New Jersey | 39.363934 | -73.635333 | 153 | NA | NA | NA | NA | 36 | | Cruise ID 202260, station 39 |
| MAB22_05860 | solidissima | 2022 | 2022 Federal Survey | NJ2 | 0 | 1 | OTU-B | B | | NA | sol | | New Jersey | 39.363934 | -73.635333 | 148.7 | 23 | NA | NA | NA | 36 | | Cruise ID 202260, station 39 |
| MAB22_05917 | solidissima | 2022 | 2022 Federal Survey | NJ2 | 0 | 1 | OTU-B | B | | NA | sol | | New Jersey | 39.363934 | -73.635333 | 129.8 | 4 | NA | NA | NA | 36 | | Cruise ID 202260, station 39 |
| MAB22_05952 | solidissima | 2022 | 2022 Federal Survey | NJ2 | 0 | 1 | OTU-B | B | | NA | sol | | New Jersey | 39.363934 | -73.635333 | 104.6 | 11 | NA | NA | NA | 36 | | Cruise ID 202260, station 39 |
| MAB22_06029 | solidissima | 2022 | 2022 Federal Survey | NJ2 | 0 | 1 | OTU-B | B | | NA | sol | | New Jersey | 39.363934 | -73.635333 | 84.9 | 4 | NA | NA | NA | 36 | | Cruise ID 202260, station 39 |
| MAB22_06571 | solidissima | 2022 | 2022 Federal Survey | NJ2 | 0 | 1 | OTU-B | B | | NA | sol | | New Jersey | 39.189125 | -73.76575 | 135.9 | 12 | NA | NA | NA | 39 | | Cruise ID 202260, station 42 |
| MAB22_06578 | solidissima | 2022 | 2022 Federal Survey | NJ2 | 0 | 1 | OTU-B | B | | NA | sol | | New Jersey | 39.189125 | -73.76575 | 144.8 | 15 | NA | NA | NA | 39 | | Cruise ID 202260, station 42 |
| MAB22_06647 | solidissima | 2022 | 2022 Federal Survey | NJ2 | 0 | 1 | OTU-B | B | | NA | sol | | New Jersey | 39.189125 | -73.76575 | 94.8 | 6 | NA | NA | NA | 39 | | Cruise ID 202260, station 42 |
| MAB22_06691 | solidissima | 2022 | 2022 Federal Survey | NJ2 | 0 | 1 | OTU-B | B | | NA | sol | | New Jersey | 39.189125 | -73.76575 | 126.7 | 13 | NA | NA | NA | 39 | | Cruise ID 202260, station 42 |

| sample | subspecies | collection year | Collection location or source | Analytical Group Abbreviation | STRUCTURE Q (OUT-A) | STRUCTURE Q (OTU-B) | ddRAD taxon, 10% hybrid threshold | ddRAD taxon, 1% hybrid threshold | majority taxon in hybrid | mtCOI *Sph* I | mtCOI *Bsp* 1286I | GTseq | analytical group location | Latitude | Longitude | Length (mm) | Age (yrs) | Width (mm) | Depth (mm) | weight (g) | water depth (m) | Region Avg water depth (m) | Notes |
|---|---|---|---|---|---|---|---|---|---|---|---|---|---|---|---|---|---|---|---|---|---|---|---|
| MAB22_06710 | solidissima | 2022 | 2022 Federal Survey | NJ2 | 0 | 1 | OTU-B | B | | NA | sol | | New Jersey | 39.189125 | -73.76575 | 117.2 | 7 | NA | NA | NA | 39 | | Cruise ID 202260, station 42 |
| MAB22_06747 | solidissima | 2022 | 2022 Federal Survey | NJ2 | 0 | 1 | OTU-B | B | | NA | sol | | New Jersey | 39.189125 | -73.76575 | 78.1 | 3 | NA | NA | NA | 39 | | Cruise ID 202260, station 42 |
| MAB22_06779 | solidissima | 2022 | 2022 Federal Survey | NJ2 | 0 | 1 | OTU-B | B | | NA | sol | | New Jersey | 39.189125 | -73.76575 | 88.6 | 4 | NA | NA | NA | 39 | | Cruise ID 202260, station 42 |
| MAB22_07043 | solidissima | 2022 | 2022 Federal Survey | NJ2 | 0 | 1 | OTU-B | B | | NA | sol | | New Jersey | 39.093709 | -73.744009 | 145.3 | 16 | NA | NA | NA | 40 | | Cruise ID 202260, station 44 |
| MAB22_07049 | solidissima | 2022 | 2022 Federal Survey | NJ2 | 0 | 1 | OTU-B | B | | NA | sol | | New Jersey | 39.093709 | -73.744009 | 117.9 | 9 | NA | NA | NA | 40 | | Cruise ID 202260, station 44 |
| MAB22_07056 | solidissima | 2022 | 2022 Federal Survey | NJ2 | 0 | 1 | OTU-B | B | | NA | sol | | New Jersey | 39.093709 | -73.744009 | 124.4 | 7 | NA | NA | NA | 40 | | Cruise ID 202260, station 44 |
| MAB22_07092 | solidissima | 2022 | 2022 Federal Survey | NJ2 | 0 | 1 | OTU-B | B | | NA | sol | | New Jersey | 39.093709 | -73.744009 | 131.2 | 13 | NA | NA | NA | 40 | | Cruise ID 202260, station 44 |
| MAB22_15450 | solidissima | 2022 | 2022 Federal Survey | NJ2 | 0 | 1 | OTU-B | B | | NA | sol | | New Jersey | 39.311358 | -74.119075 | 158.8 | 21 | NA | NA | NA | 25 | | Cruise ID 202260, station 146 |
| MAB22_15455 | solidissima | 2022 | 2022 Federal Survey | NJ2 | 0 | 1 | OTU-B | B | | NA | sol | | New Jersey | 39.311358 | -74.119075 | 141 | 20 | NA | NA | NA | 25 | | Cruise ID 202260, station 146 |
| MAB22_15457 | solidissima | 2022 | 2022 Federal Survey | NJ2 | 0 | 1 | OTU-B | B | | NA | sol | | New Jersey | 39.311358 | -74.119075 | 162.6 | 23 | NA | NA | NA | 25 | | Cruise ID 202260, station 146 |
| MAB22_15459 | solidissima | 2022 | 2022 Federal Survey | NJ2 | 0 | 1 | OTU-B | B | | NA | sol | | New Jersey | 39.311358 | -74.119075 | 90.5 | 3 | NA | NA | NA | 25 | | Cruise ID 202260, station 146 |
| WV1_001 | solidissima | 2017 | Westport, Buzzards Ba | SCC1 | 1 | 0 | OTU-A | A | | mtA | NA | | Southern Coast of Cape Cod | 41.49073 | -71.05343 | 117.4 | NA | 85.5 | 50.1 | 287.5 | 8.8 | 6.27 | |
| WV1_002 | solidissima | 2017 | Westport, Buzzards Ba | SCC1 | 1 | 0 | OTU-A | A | | no amp | NA | | Southern Coast of Cape Cod | 41.49073 | -71.05343 | 105.4 | NA | 80 | 46.9 | 199 | 8.8 | | |
| WV1_003 | solidissima | 2017 | Westport, Buzzards Ba | SCC1 | 1 | 0 | OTU-A | A | | mtA | NA | | Southern Coast of Cape Cod | 41.49073 | -71.05343 | 121 | NA | 87.5 | 50.4 | 291 | 8.8 | | |
| WV1_004 | solidissima | 2017 | Westport, Buzzards Ba | SCC1 | 1 | 0 | OTU-A | A | | mtA | NA | | Southern Coast of Cape Cod | 41.49073 | -71.05343 | 115.1 | NA | 82.2 | 48.7 | 277 | 8.8 | | |
| WV1_005 | solidissima | 2017 | Westport, Buzzards Ba | SCC1 | 1 | 0 | OTU-A | A | | mtA | NA | | Southern Coast of Cape Cod | 41.49073 | -71.05343 | 115.2 | NA | 85.2 | 48.9 | 264.5 | 8.8 | | |
| WV1_006 | solidissima | 2017 | Westport, Buzzards Ba | SCC1 | 1 | 0 | OTU-A | A | | mtA | NA | | Southern Coast of Cape Cod | 41.49073 | -71.05343 | 105.6 | NA | 75.9 | 42.6 | 199.5 | 8.8 | | |
| WV1_007 | solidissima | 2017 | Westport, Buzzards Ba | SCC1 | 1 | 0 | OTU-A | A | | mtA | NA | | Southern Coast of Cape Cod | 41.49073 | -71.05343 | 131.4 | NA | 95.5 | 56.1 | 390 | 8.8 | | |
| WV1_008 | solidissima | 2017 | Westport, Buzzards Ba | SCC1 | 1 | 0 | OTU-A | A | | mtA | NA | | Southern Coast of Cape Cod | 41.49073 | -71.05343 | 116.5 | NA | 86.2 | 50.2 | 287 | 8.8 | | |
| WV1_009 | solidissima | 2017 | Westport, Buzzards Ba | SCC1 | 1 | 0 | OTU-A | A | | no amp | NA | | Southern Coast of Cape Cod | 41.49073 | -71.05343 | 116.8 | NA | 79.9 | 46.2 | 269 | 8.8 | | |
| WV1_010 | solidissima | 2017 | Westport, Buzzards Ba | SCC1 | 0.986 | 0.014 | OTU-A | Hyb | majority A | mtA | NA | | Southern Coast of Cape Cod | 41.49073 | -71.05343 | 117.3 | NA | 88.2 | 52.3 | 316.5 | 8.8 | | |
| WV5_001 | solidissima | 2017 | Westport, Buzzards Ba | SCC1 | 1 | 0 | OTU-A | A | | mtA | NA | | Southern Coast of Cape Cod | 41.50455 | -71.0619 | 110 | NA | 75.5 | 45.1 | 226.5 | 4.3 | | |
| WV5_002 | solidissima | 2017 | Westport, Buzzards Ba | SCC1 | 1 | 0 | OTU-A | A | | no amp | NA | | Southern Coast of Cape Cod | 41.50455 | -71.0619 | 109.6 | NA | 77.9 | 43.8 | 212 | 4.3 | | |
| WV5_003 | solidissima | 2017 | Westport, Buzzards Ba | SCC1 | 1 | 0 | OTU-A | A | | mtA | NA | | Southern Coast of Cape Cod | 41.50455 | -71.0619 | 120.8 | NA | 91.4 | 54.6 | 374.5 | 4.3 | | |
| WV5_004 | solidissima | 2017 | Westport, Buzzards Ba | SCC1 | 1 | 0 | OTU-A | A | | no amp | NA | | Southern Coast of Cape Cod | 41.50455 | -71.0619 | 115.1 | NA | 82.8 | 50.4 | 266 | 4.3 | | |
| WV5_005 | solidissima | 2017 | Westport, Buzzards Ba | SCC1 | 1 | 0 | OTU-A | A | | mtA | NA | | Southern Coast of Cape Cod | 41.50455 | -71.0619 | 125.7 | NA | 90.3 | 52 | 310 | 4.3 | | |
| WV5_006 | solidissima | 2017 | Westport, Buzzards Ba | SCC1 | 1 | 0 | OTU-A | A | | mtA | NA | | Southern Coast of Cape Cod | 41.50455 | -71.0619 | 123.8 | NA | 88.7 | 53.2 | 346.5 | 4.3 | | |
| WV5_007 | solidissima | 2017 | Westport, Buzzards Ba | SCC1 | 1 | 0 | OTU-A | A | | mtA | NA | | Southern Coast of Cape Cod | 41.50455 | -71.0619 | 106.3 | NA | 72.8 | 44.3 | 204 | 4.3 | | |
| WV5_008 | solidissima | 2017 | Westport, Buzzards Ba | SCC1 | 1 | 0 | OTU-A | A | | mtA | NA | | Southern Coast of Cape Cod | 41.50455 | -71.0619 | 118.5 | NA | 83.7 | 52.6 | 305.5 | 4.3 | | |
| WV5_009 | solidissima | 2017 | Westport, Buzzards Ba | SCC1 | 1 | 0 | OTU-A | A | | mtA | NA | | Southern Coast of Cape Cod | 41.50455 | -71.0619 | 117.1 | NA | 83 | 47 | 246 | 4.3 | | |
| WV5_010 | solidissima | 2017 | Westport, Buzzards Ba | SCC1 | 1 | 0 | OTU-A | A | | mtA | NA | | Southern Coast of Cape Cod | 41.50455 | -71.0619 | 115.1 | NA | 82.2 | 43.7 | 224.5 | 4.3 | | |
| CTM_008 | solidissima | 2017 | Chatham, MA | SCC2 | 0.019 | 0.981 | OTU-B | Hyb | majority B | mtB | NA | | Southern Coast of Cape Cod | 41.670604 | -70.016384 | 44.5 | NA | 34.8 | 20.8 | 18.5 | 0.3 - 1.2 | | |
| MW_001 | solidissima | 2012 | Washburn Island, MA | SCC2 | 0 | 1 | OTU-B | B | | mtB | NA | | Southern Coast of Cape Cod | 41.552158 | -70.546503 | NA | NA | NA | NA | NA | NA | | |
| MW_002 | solidissima | 2012 | Washburn Island, MA | SCC2 | 0 | 1 | OTU-B | B | | mtB | NA | | Southern Coast of Cape Cod | 41.552158 | -70.546503 | NA | NA | NA | NA | NA | NA | | |
| MW_016 | solidissima | 2012 | Washburn Island, MA | SCC2 | 0 | 1 | OTU-B | B | | mtB | NA | | Southern Coast of Cape Cod | 41.552158 | -70.546503 | 120 | NA | NA | NA | NA | NA | | |
| MCX_015 | solidissima | 2019 | Mecox Bay, NY | SLI1 | 1 | 0 | OTU-A | A | | mtA | NA | | Southern Long Island - East | 40.887954 | -72.328834 | 65 | NA | NA | NA | NA | NA | 8.0 | |
| MCX_016 | solidissima | 2019 | Mecox Bay, NY | SLI1 | 1 | 0 | OTU-A | A | | mtA | NA | | Southern Long Island - East | 40.887954 | -72.328834 | 68 | NA | NA | NA | NA | 4.9 | | |
| MCX_017 | solidissima | 2019 | Mecox Bay, NY | SLI1 | 1 | 0 | OTU-A | A | | mtA | NA | | Southern Long Island - East | 40.887954 | -72.328834 | 83 | 3 | NA | NA | NA | 4.6 | | |
| MCX_018 | solidissima | 2019 | Mecox Bay, NY | SLI1 | 1 | 0 | OTU-A | A | | mtA | NA | | Southern Long Island - East | 40.887954 | -72.328834 | 65 | NA | NA | NA | NA | 4.6 | | |
| MCX_019 | solidissima | 2019 | Mecox Bay, NY | SLI1 | 1 | 0 | OTU-A | A | | mtA | NA | | Southern Long Island - East | 40.887954 | -72.328834 | 69 | 3 | NA | NA | NA | 4.6 | | |
| MCX_020 | solidissima | 2019 | Mecox Bay, NY | SLI1 | 1 | 0 | OTU-A | A | | mtA | NA | | Southern Long Island - East | 40.887954 | -72.328834 | 76 | NA | NA | NA | NA | 4.6 | | |
| MCX_021 | solidissima | 2019 | Mecox Bay, NY | SLI1 | 0.98 | 0.02 | OTU-A | Hyb | majority A | mtA | NA | | Southern Long Island - East | 40.887954 | -72.328834 | 76 | NA | NA | NA | NA | NA | | |
| MCX_027 | solidissima | 2019 | Mecox Bay, NY | SLI1 | 1 | 0 | OTU-A | A | | mtA | NA | | Southern Long Island - East | 40.887954 | -72.328834 | 64 | NA | NA | NA | NA | 4.9 | | |
| MCX_029 | solidissima | 2019 | Mecox Bay, NY | SLI1 | 1 | 0 | OTU-A | A | | mtA | NA | | Southern Long Island - East | 40.887954 | -72.328834 | 86 | 5 | NA | NA | NA | 5.2 | | |
| MCX_030 | solidissima | 2019 | Mecox Bay, NY | SLI1 | 1 | 0 | OTU-A | A | | mtA | NA | | Southern Long Island - East | 40.887954 | -72.328834 | 76 | NA | NA | NA | NA | 5.2 | | |
| MCX_031 | solidissima | 2019 | Mecox Bay, NY | SLI1 | 1 | 0 | OTU-A | A | | mtA | NA | | Southern Long Island - East | 40.887954 | -72.328834 | 75 | NA | NA | NA | NA | 5.2 | | |
| MCX_032 | solidissima | 2019 | Mecox Bay, NY | SLI1 | 1 | 0 | OTU-A | A | | mtA | NA | | Southern Long Island - East | 40.887954 | -72.328834 | 65 | NA | NA | NA | NA | 5.2 | | |
| MCX_033 | solidissima | 2019 | Mecox Bay, NY | SLI1 | 1 | 0 | OTU-A | A | | mtA | NA | | Southern Long Island - East | 40.887954 | -72.328834 | 60 | NA | NA | NA | NA | 5.2 | | |
| MCX_034 | solidissima | 2019 | Mecox Bay, NY | SLI1 | 1 | 0 | OTU-A | A | | mtA | NA | A | Southern Long Island - East | 40.887954 | -72.328834 | 59 | 2 | NA | NA | NA | 5.2 | | |
| MCX_035 | solidissima | 2019 | Mecox Bay, NY | SLI1 | 0.977 | 0.023 | OTU-A | Hyb | majority A | mtA | NA | | Southern Long Island - East | 40.887954 | -72.328834 | 65 | NA | NA | NA | NA | 5.2 | | |
| MCX_038 | solidissima | 2019 | Mecox Bay, NY | SLI1 | 0.921 | 0.079 | OTU-A | Hyb | majority A | mtA | NA | hyb | Southern Long Island - East | 40.887954 | -72.328834 | 68 | NA | NA | NA | NA | 5.2 | | |
| MCX_039 | solidissima | 2019 | Mecox Bay, NY | SLI1 | 1 | 0 | OTU-A | A | | mtA | NA | | Southern Long Island - East | 40.887954 | -72.328834 | 75 | NA | NA | NA | NA | 5.2 | | |
| MCX_046 | solidissima | 2019 | Mecox Bay, NY | SLI1 | 1 | 0 | OTU-A | A | | mtA | NA | | Southern Long Island - East | 40.887954 | -72.328834 | 67 | NA | NA | NA | NA | 5.5 | | |
| MCX_047 | solidissima | 2019 | Mecox Bay, NY | SLI1 | 1 | 0 | OTU-A | A | | mtA | NA | | Southern Long Island - East | 40.887954 | -72.328834 | 66 | NA | NA | NA | NA | 5.5 | | |
| MCX_055 | solidissima | 2019 | Mecox Bay, NY | SLI1 | 1 | 0 | OTU-A | A | | mtA | NA | | Southern Long Island - East | 40.887954 | -72.328834 | 62 | NA | NA | NA | NA | 6.4 | | |
| MCX_061 | solidissima | 2019 | Mecox Bay, NY | SLI1 | 1 | 0 | OTU-A | A | | mtA | NA | | Southern Long Island - East | 40.887954 | -72.328834 | 66 | NA | NA | NA | NA | 6.4 | | |
| MCX_062 | solidissima | 2019 | Mecox Bay, NY | SLI1 | 1 | 0 | OTU-A | A | | mtA | NA | | Southern Long Island - East | 40.887954 | -72.328834 | 66 | NA | NA | NA | NA | 6.4 | | |
| MCX_066 | solidissima | 2019 | Mecox Bay, NY | SLI1 | 1 | 0 | OTU-A | A | | mtA | NA | | Southern Long Island - East | 40.887954 | -72.328834 | 62 | 3 | NA | NA | NA | NA | | |
| MCX_067 | solidissima | 2019 | Mecox Bay, NY | SLI1 | 1 | 0 | OTU-A | A | | mtA | NA | | Southern Long Island - East | 40.887954 | -72.328834 | 66 | NA | NA | NA | NA | NA | | |
| MCX_068 | solidissima | 2019 | Mecox Bay, NY | SLI1 | 1 | 0 | OTU-A | A | | mtA | NA | | Southern Long Island - East | 40.887954 | -72.328834 | 65 | NA | NA | NA | NA | NA | | |
| MCX_069 | solidissima | 2019 | Mecox Bay, NY | SLI1 | 1 | 0 | OTU-A | A | | mtA | NA | | Southern Long Island - East | 40.887954 | -72.328834 | 60 | NA | NA | NA | NA | NA | | |
| MCX_079 | solidissima | 2019 | Mecox Bay, NY | SLI1 | 1 | 0 | OTU-A | A | | mtA | NA | | Southern Long Island - East | 40.887954 | -72.328834 | 64 | NA | NA | NA | NA | 6.1 | | |
| MCX_080 | solidissima | 2019 | Mecox Bay, NY | SLI1 | 1 | 0 | OTU-A | A | | mtA | NA | | Southern Long Island - East | 40.887954 | -72.328834 | 65.3 | NA | NA | NA | NA | 6.1 | | |
| MCX_084 | solidissima | 2019 | Mecox Bay, NY | SLI1 | 1 | 0 | OTU-A | A | | mtA | NA | | Southern Long Island - East | 40.887954 | -72.328834 | 76 | NA | NA | NA | NA | 6.1 | | |
| MCX_089 | solidissima | 2019 | Mecox Bay, NY | SLI1 | 1 | 0 | OTU-A | A | | mtA | NA | | Southern Long Island - East | 40.887954 | -72.328834 | 61.3 | NA | NA | NA | NA | 5.5 | | |
| MCX_090 | solidissima | 2019 | Mecox Bay, NY | SLI1 | 1 | 0 | OTU-A | A | | mtA | NA | | Southern Long Island - East | 40.887954 | -72.328834 | 61.1 | NA | NA | NA | NA | 5.5 | | |
| MCX_091 | solidissima | 2019 | Mecox Bay, NY | SLI1 | 1 | 0 | OTU-A | A | | mtA | NA | | Southern Long Island - East | 40.887954 | -72.328834 | 73.1 | NA | NA | NA | NA | 5.5 | | |
| MCX_093 | solidissima | 2019 | Mecox Bay, NY | SLI1 | 1 | 0 | OTU-A | A | | mtA | NA | | Southern Long Island - East | 40.887954 | -72.328834 | 72.1 | 4 | NA | NA | NA | 5.5 | | |
| MCX_096 | solidissima | 2019 | Mecox Bay, NY | SLI1 | 1 | 0 | OTU-A | A | | mtA | NA | | Southern Long Island - East | 40.887954 | -72.328834 | 58.7 | 2 | NA | NA | NA | 6.4 | | |
| SsLI4109_1 | solidissima | 2012 | 2012 NY Survey | SLI2 | 0.975 | 0.025 | OTU-A | Hyb | majority A | mtA | NA | | Southern Long Island - East | 40.827803 | -72.482281 | 114 | 4 | NA | NA | NA | 13 | | |
| SsLI4109_3n | solidissima | 2012 | 2012 NY Survey | SLI2 | 1 | 0 | OTU-A | A | | mtA | NA | | Southern Long Island - East | 40.827803 | -72.482281 | 150 | 17 | NA | NA | NA | 13 | | |
| SsLI4129_1 | solidissima | 2012 | 2012 NY Survey | SLI2 | 1 | 0 | OTU-A | A | | mtA | NA | A | Southern Long Island - East | 40.825859 | -72.472207 | 159 | 24 | NA | NA | NA | 16 | | |
| SsLI4129_2 | solidissima | 2012 | 2012 NY Survey | SLI2 | 0 | 1 | OTU-B | B | | mtB | NA | B | Southern Long Island - East | 40.825859 | -72.472207 | 129 | 6 | NA | NA | NA | 16 | | |
| SsLI4129_3 | solidissima | 2012 | 2012 NY Survey | SLI2 | 1 | 0 | OTU-A | A | | mtA | NA | A | Southern Long Island - East | 40.825859 | -72.472207 | 133 | 8 | NA | NA | NA | 16 | | |
| SsLI4139_1 | solidissima | 2012 | 2012 NY Survey | SLI2 | 0 | 1 | OTU-B | B | | mtB | NA | | Southern Long Island - East | 40.835173 | -72.447195 | 167 | 11 | NA | NA | NA | 14 | | |
| MAB22_21224 | solidissima | 2022 | 2022 Federal Survey | SLI2 | 0 | 1 | OTU-B | B | | NA | sol | | Southern Long Island - East | 40.803784 | -72.412242 | 116.8 | 8 | NA | NA | NA | NA | | Is separated from SLI pie diagram in fig. 5 but lumped with SLI2 for spacemix and other analyses. This |
| SHN_102 | solidissima | 2019 | Shinnicock, NY | SLI2 | 1 | 0 | OTU-A | A | | unreadable | NA | | Southern Long Island - East | 40.838522 | -72.464345 | 66.5 | 2 | NA | NA | NA | 6.1 | | |
| SHN_103 | solidissima | 2019 | Shinnicock, NY | SLI2 | 0.985 | 0.015 | OTU-A | Hyb | majority A | unreadable | NA | | Southern Long Island - East | 40.838522 | -72.464345 | 75 | 2 | NA | NA | NA | 6.1 | | |
| SHN_104 | solidissima | 2019 | Shinnicock, NY | SLI2 | 0 | 1 | OTU-B | B | | unreadable | NA | | Southern Long Island - East | 40.838522 | -72.464345 | 88 | 4 | NA | NA | NA | 6.1 | | |
| SHN_151 | solidissima | 2019 | Shinnicock, NY | SLI2 | 1 | 0 | OTU-A | A | | mtA | NA | | Southern Long Island - East | 40.838522 | -72.464345 | 67 | NA | NA | NA | NA | 4.6 | | |
| SHN_152 | solidissima | 2019 | Shinnicock, NY | SLI2 | 1 | 0 | OTU-A | A | | mtA | NA | | Southern Long Island - East | 40.838522 | -72.464345 | 60 | NA | NA | NA | NA | 4.6 | | |
| SsLI4106_1 | solidissima | 2012 | 2012 NY Survey | SLI3 | 0.002 | 0.998 | OTU-B | B | | mtB | NA | | Southern Long Island - East | 40.772146 | -72.662375 | 127 | 6 | NA | NA | NA | 16 | | |
| SsLI4106_2 | solidissima | 2012 | 2012 NY Survey | SLI3 | 0.001 | 0.999 | OTU-B | B | | mtB | NA | B | Southern Long Island - East | 40.772146 | -72.662375 | 159 | 9 | NA | NA | NA | 16 | | |
| SsLI4106_3 | solidissima | 2012 | 2012 NY Survey | SLI3 | 0.687 | 0.313 | OTU-Hyb | Hyb | majority A | mtA | NA | hyb? | Southern Long Island - East | 40.772146 | -72.662375 | 135 | 9 | NA | NA | NA | 16 | | |
| SsLI4106_4 | solidissima | 2012 | 2012 NY Survey | SLI3 | 0.02 | 0.98 | OTU-B | Hyb | majority B | mtB | NA | | Southern Long Island - East | 40.772146 | -72.662375 | 133 | 5 | NA | NA | NA | 16 | | |
| SsLI4106_5 | solidissima | 2012 | 2012 NY Survey | SLI3 | 0 | 1 | OTU-B | B | | no amp | NA | hyb | Southern Long Island - East | 40.772146 | -72.662375 | 127 | 6 | NA | NA | NA | 16 | | |
| SsLI4121_2 | solidissima | 2012 | 2012 NY Survey | SLI3 | 1 | 0 | OTU-A | A | | mtA | NA | A | Southern Long Island - East | 40.808448 | -72.572455 | 142 | 9 | NA | NA | NA | 5 | | |
| SsLI4121_3 | solidissima | 2012 | 2012 NY Survey | SLI3 | 1 | 0 | OTU-A | A | | mtA | NA | | Southern Long Island - East | 40.808448 | -72.572455 | 141 | 20 | NA | NA | NA | 5 | | |
| SsLI4121_4 | solidissima | 2012 | 2012 NY Survey | SLI3 | 1 | 0 | OTU-A | A | | mtA | NA | A | Southern Long Island - East | 40.808448 | -72.572455 | 130 | 20 | NA | NA | NA | 5 | | |
| SsLI4121_5 | solidissima | 2012 | 2012 NY Survey | SLI3 | 1 | 0 | OTU-A | A | | mtA | NA | A | Southern Long Island - East | 40.808448 | -72.572455 | 148 | 19 | NA | NA | NA | 5 | | |
| SsLI4146_1 | solidissima | 2012 | 2012 NY Survey | SLI3 | 1 | 0 | OTU-A | A | | mtA | NA | A | Southern Long Island - East | 40.789088 | -72.63187 | 154 | 22 | NA | NA | NA | NA | | |
| SsLI4146_2 | solidissima | 2012 | 2012 NY Survey | SLI3 | 1 | 0 | OTU-A | A | | mtA | NA | | Southern Long Island - East | 40.789088 | -72.63187 | 139 | 20 | NA | NA | NA | NA | | |
| SsLI3109_1 | solidissima | 2012 | 2012 NY Survey | SLI4 | 1 | 0 | OTU-A | A | | mtA | NA | | Southern Long Island - East | 40.752945 | -72.771589 | 113 | 4 | NA | NA | NA | 8 | | |
| SsLI3109_2 | solidissima | 2012 | 2012 NY Survey | SLI4 | 1 | 0 | OTU-A | A | | mtA | NA | | Southern Long Island - East | 40.752945 | -72.771589 | 156 | 22 | NA | NA | NA | 8 | | |
| SsLI3109_3 | solidissima | 2012 | 2012 NY Survey | SLI4 | 1 | 0 | OTU-A | A | | mtA | NA | A | Southern Long Island - East | 40.752945 | -72.771589 | 111 | 5 | NA | NA | NA | 8 | | |
| SsLI3109_4 | solidissima | 2012 | 2012 NY Survey | SLI4 | 0.985 | 0.015 | OTU-A | Hyb | majority A | mtA | NA | A | Southern Long Island - East | 40.752945 | -72.771589 | 142 | 7 | NA | NA | NA | 8 | | |
| SsLI3109_5 | solidissima | 2012 | 2012 NY Survey | SLI4 | 1 | 0 | OTU-A | A | | mtA | NA | A | Southern Long Island - East | 40.752945 | -72.771589 | 157 | 25 | NA | NA | NA | 8 | | |
| BLP_179n | solidissima | 2019 | Bellport, NY | SLI4 | 1 | 0 | OTU-A | A | | mtA | NA | | Southern Long Island - East | 40.71853 | -72.892526 | 44 | NA | NA | NA | NA | 6.2 | | |
| BLP_190 | solidissima | 2019 | Bellport, NY | SLI4 | 1 | 0 | OTU-A | A | | mtA | NA | | Southern Long Island - East | 40.71853 | -72.892526 | 62 | NA | NA | NA | NA | 5.5 | | |
| BLP_191 | solidissima | 2019 | Bellport, NY | SLI4 | 1 | 0 | OTU-A | A | | mtA | NA | | Southern Long Island - East | 40.71853 | -72.892526 | NA | NA | NA | NA | NA | 5.5 | | |
| BLP_192 | solidissima | 2019 | Bellport, NY | SLI4 | 1 | 0 | OTU-A | A | | mtA | NA | | Southern Long Island - East | 40.71853 | -72.892526 | NA | NA | NA | NA | NA | 5.5 | | |
| BLP_193 | solidissima | 2019 | Bellport, NY | SLI4 | 1 | 0 | OTU-A | A | | mtA | NA | | Southern Long Island - East | 40.71853 | -72.892526 | NA | NA | NA | NA | NA | 5.5 | | |
| BLP_194 | solidissima | 2019 | Bellport, NY | SLI4 | 1 | 0 | OTU-A | A | | mtA | NA | | Southern Long Island - East | 40.71853 | -72.892526 | NA | NA | NA | NA | NA | 5.5 | | |
| BLP_195 | solidissima | 2019 | Bellport, NY | SLI4 | 0.999 | 0.001 | OTU-A | A | | mtA | NA | | Southern Long Island - East | 40.71853 | -72.892526 | NA | NA | NA | NA | NA | 5.5 | | |
| BLP_198n | solidissima | 2019 | Bellport, NY | SLI4 | 0.91 | 0.09 | OTU-A | Hyb | majority A | mtA | NA | A | Southern Long Island - East | 40.71853 | -72.892526 | NA | NA | NA | NA | NA | 4.6 | | |
| BLP_199n | solidissima | 2019 | Bellport, NY | SLI4 | 1 | 0 | OTU-A | A | | mtA | NA | | Southern Long Island - East | 40.71853 | -72.892526 | NA | NA | NA | NA | NA | 4.6 | | |
| BLP_212 | solidissima | 2019 | Bellport, NY | SLI4 | 1 | 0 | OTU-A | A | | mtA | NA | | Southern Long Island - East | 40.71853 | -72.892526 | 66.5 | 2 | NA | NA | NA | 4.6 | | |
| BLP_213 | solidissima | 2019 | Bellport, NY | SLI4 | 0 | 1 | OTU-B | B | | unreadable | NA | | Southern Long Island - East | 40.71853 | -72.892526 | 61.7 | 2 | NA | NA | NA | 4.6 | | |
| BLP_231 | solidissima | 2019 | Bellport, NY | SLI4 | 1 | 0 | OTU-A | A | | unreadable | NA | | Southern Long Island - East | 40.71853 | -72.892526 | 74 | NA | NA | NA | NA | 6.4 | | |
| BLP_233 | solidissima | 2019 | Bellport, NY | SLI4 | 0.755 | 0.245 | OTU-Hyb | Hyb | majority A | unreadable | NA | | Southern Long Island - East | 40.71853 | -72.892526 | 63 | 4 | NA | NA | NA | NA | | |
| BLP_234 | solidissima | 2019 | Bellport, NY | SLI4 | 0.994 | 0.006 | OTU-A | A | | unreadable | NA | | Southern Long Island - East | 40.71853 | -72.892526 | 78 | 2 | NA | NA | NA | NA | | |
| BLP_237 | solidissima | 2019 | Bellport, NY | SLI4 | 0.993 | 0.007 | OTU-A | A | | unreadable | NA | | Southern Long Island - East | 40.71853 | -72.892526 | 75 | 3 | NA | NA | NA | 4.0 | | |
| SsLI2101_7 | solidissima | 2012 | 2012 NY Survey | SLI5 | 0 | 1 | OTU-B | B | | mtA | NA | | Southern Long Island - West | 40.609683 | -73.308927 | NA | NA | NA | NA | NA | 14 | | |
| SsLI2109_2 | solidissima | 2012 | 2012 NY Survey | SLI5 | 1 | 0 | OTU-A | A | | mtA | NA | | Southern Long Island - West | 40.607649 | -73.314017 | 116 | 14 | NA | NA | NA | 14 | | |
| SsLI2109_3 | solidissima | 2012 | 2012 NY Survey | SLI5 | 0.985 | 0.015 | OTU-A | Hyb | majority A | mtA | NA | | Southern Long Island - West | 40.607649 | -73.314017 | 138 | 16 | NA | NA | NA | 14 | | |
| SsLI2109_4 | solidissima | 2012 | 2012 NY Survey | SLI5 | 1 | 0 | OTU-A | A | | mtA | NA | A | Southern Long Island - West | 40.607649 | -73.314017 | 146 | 21 | NA | NA | NA | 14 | | |
| SsLI3105_2 | solidissima | 2012 | 2012 NY Survey | SLI5 | 1 | 0 | OTU-A | A | | mtA | NA | | Southern Long Island - West | 40.609979 | -73.273337 | 141 | 24 | NA | NA | NA | 13 | | |
| SsLI3105_3 | solidissima | 2012 | 2012 NY Survey | SLI5 | 1 | 0 | OTU-A | A | | mtA | NA | | Southern Long Island - West | 40.609979 | -73.273337 | 147 | 16 | NA | NA | NA | 13 | | |
| SsLI3105_5 | solidissima | 2012 | 2012 NY Survey | SLI5 | 1 | 0 | OTU-A | A | | mtA | NA | | Southern Long Island - West | 40.609979 | -73.273337 | 157 | 21 | NA | NA | NA | 13 | | |
| SsLI1101_1 | solidissima | 2012 | 2012 NY Survey | SLI6 | 1 | 0 | OTU-A | A | | mtA | NA | | Southern Long Island - West | 40.565793 | -73.817567 | 98 | 3 | NA | NA | NA | 9 | | |
| SsLI1102_1 | solidissima | 2012 | 2012 NY Survey | SLI6 | 1 | 0 | OTU-A | A | | mtA | NA | | Southern Long Island - West | 40.573978 | -73.601822 | 103 | 4 | NA | NA | NA | 6 | | |
| SsLI1102_2 | solidissima | 2012 | 2012 NY Survey | SLI6 | 1 | 0 | OTU-A | A | | mtA | NA | | Southern Long Island - West | 40.573978 | -73.601822 | 114 | 8 | NA | NA | NA | 6 | | |
| SsLI1102_3 | solidissima | 2012 | 2012 NY Survey | SLI6 | 1 | 0 | OTU-A | A | | mtA | NA | | Southern Long Island - West | 40.573978 | -73.601822 | 148 | 15 | NA | NA | NA | 6 | | |
| SsLI1102_4 | solidissima | 2012 | 2012 NY Survey | SLI6 | 1 | 0 | OTU-A | A | | mtA | NA | | Southern Long Island - West | 40.573978 | -73.601822 | 145 | 8 | NA | NA | NA | 6 | | |
| SsLI1107_1 | solidissima | 2012 | 2012 NY Survey | SLI6 | 0.992 | 0.008 | OTU-A | A | | mtA | NA | A | Southern Long Island - West | 40.570486 | -73.742964 | 125 | 13 | NA | NA | NA | 11 | | |
| SsLI1107_2 | solidissima | 2012 | 2012 NY Survey | SLI6 | 1 | 0 | OTU-A | A | | mtA | NA | A | Southern Long Island - West | 40.570486 | -73.742964 | 128 | 13 | NA | NA | NA | 11 | | |
| SsLI1107_3 | solidissima | 2012 | 2012 NY Survey | SLI6 | 1 | 0 | OTU-A | A | | mtA | NA | A | Southern Long Island - West | 40.570486 | -73.742964 | 149 | 13 | NA | NA | NA | 11 | | |
| SsLI1112_1 | solidissima | 2012 | 2012 NY Survey | SLI6 | 0.889 | 0.111 | OTU-Hyb | Hyb | majority A | mtA | NA | hyb | Southern Long Island - West | 40.574498 | -73.772801 | 115 | 23 | NA | NA | NA | 9 | | |
| SsLI1112_2 | solidissima | 2012 | 2012 NY Survey | SLI6 | 1 | 0 | OTU-A | A | | mtA | NA | A | Southern Long Island - West | 40.574498 | -73.772801 | 148 | 19 | NA | NA | NA | 9 | | |
| SsLI1113_4 | solidissima | 2012 | 2012 NY Survey | SLI6 | 0.854 | 0.146 | OTU-Hyb | Hyb | majority A | mtA | NA | hyb | Southern Long Island - West | 40.571067 | -73.599508 | 130 | 8 | NA | NA | NA | 7 | | |
| SsLI1113_5 | solidissima | 2012 | 2012 NY Survey | SLI6 | 0.992 | 0.008 | OTU-A | A | | mtA | NA | A | Southern Long Island - West | 40.571067 | -73.599508 | 148 | 14 | NA | NA | NA | 7 | | |
| SsLI1116_2 | solidissima | 2012 | 2012 NY Survey | SLI6 | 0.999 | 0.001 | OTU-A | A | | mtA | NA | | Southern Long Island - West | 40.572322 | -73.79214 | 120 | 7 | NA | NA | NA | 9 | | |
| SsLI1117_2 | solidissima | 2012 | 2012 NY Survey | SLI6 | 0 | 1 | OTU-B | B | | mtB | NA | | Southern Long Island - West | 40.572977 | -73.822306 | 102 | 4 | NA | NA | NA | 7 | | |
| SsLI1117_4 | solidissima | 2012 | 2012 NY Survey | SLI6 | 1 | 0 | OTU-A | A | | mtA | NA | | Southern Long Island - West | 40.572977 | -73.822306 | 100 | 3 | NA | NA | NA | 7 | | |
| SsLI2107_1 | solidissima | 2012 | 2012 NY Survey | SLI6 | 1 | 0 | OTU-A | A | | mtA | NA | A | Southern Long Island - West | 40.56614 | -73.565544 | 150 | 8 | NA | NA | NA | 9 | | |
| SsLI2107_2 | solidissima | 2012 | 2012 NY Survey | SLI6 | 1 | 0 | OTU-A | A | | mtA | NA | | Southern Long Island - West | 40.56614 | -73.565544 | 128 | 7 | NA | NA | NA | 9 | | |
| SsLI2108_1 | solidissima | 2012 | 2012 NY Survey | SLI6 | 1 | 0 | OTU-A | A | | mtA | NA | A | Southern Long Island - West | 40.562769 | -73.568494 | 123 | 8 | NA | NA | NA | 11 | | |
| SsLI2108_2n | solidissima | 2012 | 2012 NY Survey | SLI6 | 0.982 | 0.018 | OTU-A | Hyb | majority A | mtA | NA | A | Southern Long Island - West | 40.562769 | -73.568494 | 124 | 8 | NA | NA | NA | 11 | | |
| SsLI2108_3 | solidissima | 2012 | 2012 NY Survey | SLI6 | 0.951 | 0.049 | OTU-A | Hyb | majority A | mtA | NA | A | Southern Long Island - West | 40.562769 | -73.568494 | 137 | 8 | NA | NA | NA | 11 | | |
| MAB22_10193 | similis | 2022 | 2022 Federal Survey | NA | NA | NA | SIM | SIM | | NA | sim | | NA | 36.668542 | -75.423092 | 101.1 | 5 | NA | NA | NA | 20 | | Cruise ID 202260, station 80 |
| MAB22_10194 | similis | 2022 | 2022 Federal Survey | NA | NA | NA | SIM | SIM | | NA | sim | sim | NA | 36.668542 | -75.423092 | 89.3 | 4 | NA | NA | NA | 20 | | Cruise ID 202260, station 80 |
| MAB22_10773 | similis | 2022 | 2022 Federal Survey | NA | NA | NA | SIM | SIM | | NA | sim | | NA | 36.608 | -75.018534 | 75.8 | 5 | NA | NA | NA | 26 | | Cruise ID 202260, station 83 |
| MAB22_11137 | similis | 2022 | 2022 Federal Survey | NA | NA | NA | SIM | SIM | | NA | sim | | NA | 36.608 | -75.018534 | 66.7 | 5 | NA | NA | NA | 26 | | Cruise ID 202260, station 83 |
| MAB22_11535 | similis | 2022 | 2022 Federal Survey | NA | NA | NA | SIM | SIM | | NA | sim | | NA | 36.789492 | -74.937092 | 85.7 | 5 | NA | NA | NA | 31 | | Cruise ID 202260, station 85 |
| MAB22_15461 | similis | 2022 | 2022 Federal Survey | NA | NA | NA | SIM | SIM | | NA | sim | | NA | 39.731842 | -74.038892 | 95 | 3 | NA | NA | NA | 14 | | Cruise ID 202260, station 148 |
| MAB22_15465 | similis | 2022 | 2022 Federal Survey | NA | NA | NA | SIM | SIM | | NA | sim | | NA | 39.731167 | -73.966625 | 83.8 | 3 | NA | NA | NA | 20 | | Cruise ID 202260, station 149 |
| MAB22_10201 | similis | 2022 | 2022 Federal Survey | NA | NA | NA | SIM-SOL-Hyb | SIM-SOL-Hyb | 50/50 F1 | NA | sim | SIM-SOL-Hyb | NA | 36.620733 | -75.086892 | 102.4 | 9 | NA | NA | NA | 27 | | Cruise ID 202260, station 82 |
| ELP_001 | similis (reference) | 2017 | Eel Pond, Washburn Is. | NA | NA | NA | SIM | SIM | | NA | NA | | NA | 41.554 | -70.546 | 94.9 | NA | 65.1 | 39.3 | 109.5 | 0.9 - 1.2 | | |
| ELP_002 | similis (reference) | 2017 | Eel Pond, Washburn Is. | NA | NA | NA | SIM | SIM | | NA | NA | | NA | 41.554 | -70.546 | 83 | NA | 59.1 | 37.1 | 72.4 | 0.9 - 1.2 | | |
| ELP_003 | similis (reference) | 2017 | Eel Pond, Washburn Is. | NA | NA | NA | SIM | SIM | | NA | NA | | NA | 41.554 | -70.546 | 82.4 | NA | 60.4 | 35.6 | 77.4 | 0.9 - 1.2 | | |
| ELP_004 | similis (reference) | 2017 | Eel Pond, Washburn Is. | NA | NA | NA | SIM | SIM | | NA | NA | | NA | 41.554 | -70.546 | 99.1 | NA | 69.1 | 41.9 | 128.6 | 0.9 - 1.2 | | |
| ELP_005 | similis (reference) | 2017 | Eel Pond, Washburn Is. | NA | NA | NA | SIM | SIM | | NA | NA | | NA | 41.554 | -70.546 | 88.8 | NA | 64.2 | 39.1 | 87.4 | 0.9 - 1.2 | | |
| ELP_006 | similis (reference) | 2017 | Eel Pond, Washburn Is. | NA | NA | NA | SIM | SIM | | NA | NA | | NA | 41.554 | -70.546 | 101.4 | NA | 70.3 | 42.9 | 132.6 | 0.9 - 1.2 | | |
| ELP_007 | similis (reference) | 2017 | Eel Pond, Washburn Is. | NA | NA | NA | SIM | SIM | | NA | NA | | NA | 41.554 | -70.546 | 60.3 | NA | 43.4 | 24.8 | 27 | 0.9 - 1.2 | | |
| ELP_008 | similis (reference) | 2017 | Eel Pond, Washburn Is. | NA | NA | NA | SIM | SIM | | NA | NA | | NA | 41.554 | -70.546 | 81.8 | NA | 59.6 | 33.8 | 72.3 | 0.9 - 1.2 | | |
| ELP_009 | similis (reference) | 2017 | Eel Pond, Washburn Is. | NA | NA | NA | SIM | SIM | | NA | NA | | NA | 41.554 | -70.546 | 82 | NA | 57.8 | 36.3 | 78 | 0.9 - 1.2 | | |
| ELP_010 | similis (reference) | 2017 | Eel Pond, Washburn Is. | NA | NA | NA | SIM | SIM | | NA | NA | | NA | 41.554 | -70.546 | 85.5 | NA | 58.2 | 36.2 | 76.3 | 0.9 - 1.2 | | |
| MW003 | similis (reference) | 2012 | Washburn Is., MA | NA | NA | NA | SIM | NA | | NA | NA | sim | NA | 41.552 | -70.546 | NA | NA | NA | NA | NA | 0.9 - 1.2 | | |
| MW037 | similis (reference) | 2012 | Washburn Is., MA | NA | NA | NA | SIM | NA | | NA | NA | sim | NA | 41.552 | -70.546 | NA | NA | NA | NA | NA | 0.9 - 1.2 | | |
| MW038 | similis (reference) | 2012 | Washburn Is., MA | NA | NA | NA | SIM | NA | | NA | NA | sim | NA | 41.552 | -70.546 | NA | NA | NA | NA | NA | 0.9 - 1.2 | | |
| MW039 | similis (reference) | 2012 | Washburn Is., MA | NA | NA | NA | SIM | NA | | NA | NA | sim | NA | 41.552 | -70.546 | NA | NA | NA | NA | NA | 0.9 - 1.2 | | |
| MW040 | similis (reference) | 2012 | Washburn Is., MA | NA | NA | NA | SIM | NA | | NA | NA | sim | NA | 41.552 | -70.546 | NA | NA | NA | NA | NA | 0.9 - 1.2 | | |
| MW041 | similis (reference) | 2012 | Washburn Is., MA | NA | NA | NA | SIM | NA | | NA | NA | sim | NA | 41.552 | -70.546 | NA | NA | NA | NA | NA | 0.9 - 1.2 | | |
| MW042 | similis (reference) | 2012 | Washburn Is., MA | NA | NA | NA | SIM | NA | | NA | NA | sim | NA | 41.552 | -70.546 | NA | NA | NA | NA | NA | 0.9 - 1.2 | | |
| MW043 | similis (reference) | 2012 | Washburn Is., MA | NA | NA | NA | SIM | NA | | NA | NA | sim | NA | 41.552 | -70.546 | NA | NA | NA | NA | NA | 0.9 - 1.2 | | |
| MW044 | similis (reference) | 2012 | Washburn Is., MA | NA | NA | NA | SIM | NA | | NA | NA | sim | NA | 41.552 | -70.546 | NA | NA | NA | NA | NA | 0.9 - 1.2 | | |
| MW045 | similis (reference) | 2012 | Washburn Is., MA | NA | NA | NA | SIM | NA | | NA | NA | sim | NA | 41.552 | -70.546 | NA | NA | NA | NA | NA | 0.9 - 1.2 | | |
| MW046 | similis (reference) | 2012 | Washburn Is., MA | NA | NA | NA | SIM | NA | | NA | NA | sim | NA | 41.552 | -70.546 | NA | NA | NA | NA | NA | 0.9 - 1.2 | | |
| MW047 | similis (reference) | 2012 | Washburn Is., MA | NA | NA | NA | SIM | NA | | NA | NA | sim | NA | 41.552 | -70.546 | NA | NA | NA | NA | NA | 0.9 - 1.2 | | |
| MW049 | similis (reference) | 2012 | Washburn Is., MA | NA | NA | NA | SIM | NA | | NA | NA | sim | NA | 41.552 | -70.546 | NA | NA | NA | NA | NA | 0.9 - 1.2 | | |
| MW050 | similis (reference) | 2012 | Washburn Is., MA | NA | NA | NA | SIM | NA | | NA | NA | sim | NA | 41.552 | -70.546 | NA | NA | NA | NA | NA | 0.9 - 1.2 | | |
| MW051 | similis (reference) | 2012 | Washburn Is., MA | NA | NA | NA | SIM | NA | | NA | NA | sim | NA | 41.552 | -70.546 | NA | NA | NA | NA | NA | 0.9 - 1.2 | | |
| MW052 | similis (reference) | 2012 | Washburn Is., MA | NA | NA | NA | SIM | NA | | NA | NA | sim | NA | 41.552 | -70.546 | NA | NA | NA | NA | NA | 0.9 - 1.2 | | |
| MW053 | similis (reference) | 2012 | Washburn Is., MA | NA | NA | NA | SIM | NA | | NA | NA | sim | NA | 41.552 | -70.546 | NA | NA | NA | NA | NA | 0.9 - 1.2 | | |
| MW054 | similis (reference) | 2012 | Washburn Is., MA | NA | NA | NA | SIM | NA | | NA | NA | sim | NA | 41.552 | -70.546 | NA | NA | NA | NA | NA | 0.9 - 1.2 | | |
| MW055 | similis (reference) | 2012 | Washburn Is., MA | NA | NA | NA | SIM | NA | | NA | NA | sim | NA | 41.552 | -70.546 | NA | NA | NA | NA | NA | 0.9 - 1.2 | | |
| MW056 | similis (reference) | 2012 | Washburn Is., MA | NA | NA | NA | SIM | NA | | NA | NA | sim | NA | 41.552 | -70.546 | NA | NA | NA | NA | NA | 0.9 - 1.2 | | |
| MW057 | similis (reference) | 2012 | Washburn Is., MA | NA | NA | NA | SIM | NA | | NA | NA | sim | NA | 41.552 | -70.546 | NA | NA | NA | NA | NA | 0.9 - 1.2 | | |
| MW058 | similis (reference) | 2012 | Washburn Is., MA | NA | NA | NA | SIM | NA | | NA | NA | sim | NA | 41.552 | -70.546 | NA | NA | NA | NA | NA | 0.9 - 1.2 | | |
| MW059 | similis (reference) | 2012 | Washburn Is., MA | NA | NA | NA | SIM | NA | | NA | NA | sim | NA | 41.552 | -70.546 | NA | NA | NA | NA | NA | 0.9 - 1.2 | | |
| MW060 | similis (reference) | 2012 | Washburn Is., MA | NA | NA | NA | SIM | NA | | NA | NA | sim | NA | 41.552 | -70.546 | NA | NA | NA | NA | NA | 0.9 - 1.2 | | |
| MW061 | similis (reference) | 2012 | Washburn Is., MA | NA | NA | NA | SIM | NA | | NA | NA | sim | NA | 41.552 | -70.546 | NA | NA | NA | NA | NA | 0.9 - 1.2 | | |
| MW062 | similis (reference) | 2012 | Washburn Is., MA | NA | NA | NA | SIM | NA | | NA | NA | sim | NA | 41.552 | -70.546 | NA | NA | NA | NA | NA | 0.9 - 1.2 | | |
| MW065 | similis (reference) | 2012 | Washburn Is., MA | NA | NA | NA | SIM | NA | | NA | NA | sim | NA | 41.552 | -70.546 | NA | NA | NA | NA | NA | 0.9 - 1.2 | | |
| MW066 | similis (reference) | 2012 | Washburn Is., MA | NA | NA | NA | SIM | NA | | NA | NA | sim | NA | 41.552 | -70.546 | NA | NA | NA | NA | NA | 0.9 - 1.2 | | |

*Table S.2: Sequence quality control filtering steps*

| **Criteria for filtering out** | **All samples plus 10 reference *S.s. similis* individuals (n=558)** | ***S.s. solidissima* only (n=540)** |
|---|---|---|
| Quality < 30<br>Minimum Depth < 3 | 703,029 loci | 696,245 loci |
| Missingness > 15%<br>Mean Depth < 20<br>MAF < 0.05 | 32,118 loci | 38,776 loci |
| Missingness < 10% from any regional population | 32,115 loci | 38,760 loci |
| Default recommendations of Puritz's SNP Filtering Tutorial (see details in Method) | 24,407 loci | 29,512 loci |
| Recode as SNPs and remove indels | 25,065 SNPs | 30,552 SNPs |
| Population-specific deviations from Hardy-Weinberg Equilibrium ($p < 0.001$) | 22,683 SNPs | 30,137 SNPs |
| Filter to a single SNP with the highest minor allele frequency per contig | 8,681 SNPs | 9,739 SNPs |
| LD pruning to remove one contig from each pair of contigs having $r^2 > 0.8$ | 8,628 SNPs<br>(8,585 biallelic SNPs) | 9,634 SNPs<br>(9,597 biallelic SNPs) |

### Table S.3: Primers, probe and amplicon sequence for 113 GTseq panel loci

| "113" locus set for GTseq | primer1 | primer2 RC | Probe seq | Probe notes | original seq with [polymorphism] |
|---|---|---|---|---|---|
| dDocent_Contig_10060_227 | ACGTACGTATTTGCATGGTGA | CCGTTGTCACAAACTGAGCA | ACGTACGTATTTGCATGGTGA | Seq | ACGTACGTATTTGCATGGTGAACTTTATATAATCACAACATACCTTGTTGTTTCATAAAA[A/C]TGTCCAGATTTACAAAGTAGGCGTTACCGTTGTCACAAACTGAGCA |
| dDocent_Contig_10178_50 | TGCAGCTGTAAACTAGTTTCATTGA | TGAAGCATTAACCACATTGTGTCA | TGCAGCTGTAAACTAGTTTCATTGA | Seq | TGCAGCTGTAAACTAGTTTCATTGAAACTTGAGGATATTTGTACTACA[C/T]TTTTCTAATTGTACTGAGTACATTATATACATGTATGATGTTATTTTTGTTATCATGAAGCATTAACCACATTGTGTCA |
| dDocent_Contig_10354_46 | GCAGACGTTAACCGCTACCT | AATACATACCACGTGCGGGC | GCAGACGTTAACCGCTACCT | Seq | GCAGACGTTAACCGCTACCTGAGCCGCTTCATAATGCTCTACT[G/T]ATGGAGAGCTAATTTCAATACTTCAAACGATCCAGTATTTTATATATCTATCATCTGTGAAAATACATACCACGTGCGGGC |
| dDocent_Contig_10374_79 | TTGTGTAGGCGCAATCAAGT | TGGCATATCTTCAAAGGTCGGT | ACCGACCTTTGAAGATATGCCA | Seq_reverse_complemented | ACCGACCTTTGAAGATATGCCAAAATAACTGAAAAAAGTTGGTTTGTTTTTGATAACA[C/T]ATGTTTTGTAGTATATATGTTTTCAACACAAATACTCAAAGGCGTATATACTTGATTGCGCCTACACAA |
| dDocent_Contig_10680_212 | GCACAGCGGGCGAGATATAA | TCTGGTGTGACTCAAATCCTCC | GCACAGCGGGCGAGATATAA | Seq | GCACAGCGGGCGAGATATAAGATTCAGTGACATGATACCAAACATTAGTTT[G/C]CTTGTATCGTAGAAAACTGGCATACATTCTGGTGTGACTCAAATCCTCC |
| dDocent_Contig_11824_163 | CCTCAGACATTAGAAAGGCAAGC | TAACTGACTCCAGGGAGCAC | CCTCAGACATTAGAAAGGCAAGC | Seq | CCTCAGACATTAGAAAGGCAAGCAATAAAGGANNNNNNNNNNNATTATGTTT[G/T]TGTCTAATGAAATTGCTGTTTAACTGACTCCAGGGAGCAC |
| dDocent_Contig_12150_168 | TCTCCCTCTCGAGCTGTCAA | AGGTGATGAATGTAAGTCCTGGT | ACCAGGACTTACATTCATCACCT | Seq_reverse_complemented | ACCAGGACTTACATTCATCACCTCCAGAATCTGGCTT[A/T]CTTGTGCAAATCAANNNNNNNNNNNCAATTGCAGAAATGACGTGTATTTTGTATAATGTAACTTTATGTTGACAGCTCGAGAGGGAGA |
| dDocent_Contig_12333_56 | TGCAGAAGCTGGATTTAAGTGG | GGTGATCCATCTACGTAAGGTATTT | TGCAGAAGCTGGATTTAAGTGG | Seq | TGCAGAAGCTGGATTTAAGTGGAATGAACTTATTATCAACAAATAGTCCCCATT[G/T]AAGTCGAATCATGTTTTGACTCTCGAAATAAGTGTTTTGGTGATCCATCTACGTAAGGTATTT |
| dDocent_Contig_12594_34 | TGCAGTAAGTATCCAGTCAAGTGA | GCTTAGATCGGGTGGCTGTA | TGCAGTAAGTATCCAGTCAAGTGA | Seq | TGCAGTAAGTATCCAGTCAAGTGATCAGTCAA[A/C]TGTGACTGCTTAGATCGGGTGGCTGTA |
| dDocent_Contig_12904_64 | TGCAGATTGTGTCAGTCAGAGA | TTGGCGGCATACAGTTCTGT | TGCAGATTGTGTCAGTCAGAGA | Seq | TGCAGATTGTGTCAGTCAGAGATTGCTATTGCAAAGTAATTGATCTAACGCCACAATAAGTT[A/T]ACAATCTATGGCCAGTATTCTTCTTAGATAGTGTATGATGCCCATTTGGCGGCATACAGTTCTGT |
| dDocent_Contig_13123_190 | AGTCTACAAACGCACCACCA | AACTGTGCACACTGACTCGA | AGTCTACAAACGCACCACCA | Seq | AGTCTACAAACGCACCACCATG[T/C]ACCACAAATGTATTATTTGTACAATGAGTTTTTAAATTAATTCGAAACTGTGCACACTGACTCGA |
| dDocent_Contig_13675_172 | ACCCCGCGTGGCTTTATTTA | GTAACTCGGCACAACAGGGA | TCCCTGTTGTGCCGAGTTAC | Seq_reverse_complemented | TCCCTGTTGTGCCGAGTTACTGAGATCTTCT[G/C]CAATGTMCTCAACAAATGNNNNNNNNNNNTTGTCCATTAATTAATATTTGCATAAGAATGAACATTAAAAATGCTAGAATAAATAAAGCCACGCGGGGT |
| dDocent_Contig_13735_158 | TCGTTGTTTCGTGTTTCACCA | TACTTCTTTGCAGCGATGGC | TCGTTGTTTCGTGTTTCACCA | Seq | TCGTTGTTTCGTGTTTCACCATGANNNNNNNNNNNTTTT[A/C]ACCATCGTACATTCAAGTTCGTGGTTTCGTGTTTTTGCCATCGTAGTTTCGTATTTTCGTTGTTTCAAGTTACTTCTTTGCAGCGATGGC |
| dDocent_Contig_13896_82 | GCCATGACAACCGATGTGTG | GGTTCATTCGAGAAACATCGGG | GCCATGACAACCGATGTGTG | Seq | GCCATGACAACCGATGTGTGTGTCACGTGATCAGAAAAAAACACCGACATGTC[A/G]AGTACTTAAGGTTCATTCGAGAAACATCGGG |
| dDocent_Contig_14380_67 | AGCAGTTTCGACCAACCAGT | CATTCAAAGCCGCTGTCTGG | AGCAGTTTCGACCAACCAGT | Seq | AGCAGTTTCGACCAACCAGTTACATTATCTGCGTGTGGAATAATGTACTCGTCCATATCAAT[A/G]TTAACAATAAATTCTGACGATAACTTACTCCTAAAAAGACAATCATTCAAAGCCGCTGTCTGG |
| dDocent_Contig_15313_112 | TGGCACGTAATGACATGAACA | GCAGCTATATCTGTGTTAGAAGTGC | GCACTTCTAACACAGATATAGCTGC | Seq_reverse_complemented | GCACTTCTAACACAGATATAGCTGCA[T/A]ACTAGTTAACTCCGTTTTTATAACGAATTCCTTGAACATTTACCTAGTGAAACATACATTGCATAAAATAAATTGTTCATGTCATTACGTGCCA |
| dDocent_Contig_15353_232 | AATCGCCTCTCGCTGGAAAA | AAGAACTTAGTTTTATTGCATGCGT | AATCGCCTCTCGCTGGAAAA | Seq | AATCGCCTCTCGCTGGAAAAATTAAGCTGTTGTAAATTTAATTAAAAATCGCATTTTAAAGATC[C/A]CAAAGAACTTAGTTTTATTGCATGCGT |
| dDocent_Contig_15742_241 | ATTGGTAACCGTTTCTTTATGCG | AGAAATCAGCAAGCAAACTTATTCA | ATTGGTAACCGTTTCTTTATGCG | Seq | ATTGGTAACCGTTTCTTTATGCGGTTGGTTTGAAT[C/A]AAAGAAATCAGCAAGCAAACTTATTCA |
| dDocent_Contig_15767_86 | TCACAGATGGCTGACAGTGT | ACCCTTGGTGTATGTTATTTTGTCA | TCACAGATGGCTGACAGTGT | Seq | TCACAGATGGCTGACAGTGTAAGTATTAACATTAAATATAAGCATTTATTTTTG[A/C]ATTTTCTAAGTGTCACCCTTGGTGTATGTTATTTTGTCA |
| dDocent_Contig_15805_248 | GCAAGACAAACGCCGATACC | GACGCCGTACTAACCAGCTA | GCAAGACAAACGCCGATACC | Seq | GCAAGACAAACGCCGATACCGATATTGAACGCCATATTTAGGGTTAGAATA[G/A]CACCCAACGCGACGCCGTACTAACCAGCTA |
| dDocent_Contig_15918_232 | TCCCTTACTGAGAAACAATTCGGT | TATGTGCGGCACTGAGTTGT | TCCCTTACTGAGAAACAATTCGGT | Seq | TCCCTTACTGAGAAACAATTCGGTTTATTAACGGTATTTGAAATGATACGTCATATGGTATAAMTTTT[G/A]ATAACCAGAACACATTTCAATATGTGCGGCACTGAGTTGT |
| dDocent_Contig_15946_143 | ACGAGTATTTACGATCTGACCTGT | TTTTCAGATGGCCACTAACACA | ACGAGTATTTACGATCTGACCTGT | Seq | ACGAGTATTTACGATCTGACCTGTTTTAGATAGCCATTCAATTGATACATCAA[C/A]NNNNNNNNNNNAATGATAGAACTTGCTTAAGACAGATGATTGCTTTTTTCAGATGGCCACTAACACA |
| dDocent_Contig_16056_191 | ACTCAGACAAACGTTTACCATTTCT | ACTACAGTTGCACGAATACATGT | ACTCAGACAAACGTTTACCATTTCT | Seq | ACTCAGACAAACGTTTACCATTTCTATT[T/C]AATACTACAGTTGCACGAATACATGT |
| dDocent_Contig_16256_188 | TGCTGTATCTTGCATTCCGATG | CCCGTCACAAATGCTCATTCA | TGCTGTATCTTGCATTCCGATG | Seq | TGCTGTATCTTGCATTCCGATGCTATGC[A/G]ATTCGGCCACACCCGTCACAAATGCTCATTCA |
| dDocent_Contig_16521_72 | TCTTCCTTTCAGAACCACGCT | TCGGTTCTGTGTTTCTAGCGT | TCTTCCTTTCAGAACCACGCT | Seq | TCTTCCTTTCAGAACCACGCTTTAAAAGTGGGTGTAAG[G/C]AAACGATTGTTAAAGTTATATAATTTGCACATTTATTTGTCGGTTCTGTGTTTCTAGCGT |
| dDocent_Contig_16522_214 | TGAAATAGACATTCACATGTGAGCA | AGGATCAAGGTGTCGATATTGTCA | TGAAATAGACATTCACATGTGAGCA | Seq | TGAAATAGACATTCACATGTGAGCATGTGTTACATGCATATA[C/T]AAGGATCAAGGTGTCGATATTGTCA |
| dDocent_Contig_17195_211 | CAACAGTCGCTAAATGGATACGA | CCCACCGCCATTATTCCAGT | CAACAGTCGCTAAATGGATACGA | Seq | CAACAGTCGCTAAATGGATACGACTTAT[A/G]TAATATTTCATTCTTAATCCCCACCGCCATTATTCCAGT |
| dDocent_Contig_17558_82 | TGCCATCCATTCCCTAGTGG | TGTGCATCACTGTAAAGTTGTCT | TGCCATCCATTCCCTAGTGG | Seq | TGCCATCCATTCCCTAGTGGTCTTCTAKATGCTATATGGTATTTTAGTCTTGGAAACATT[T/C]CGTTAACACTTATATTTATGTGCATCACTGTAAAGTTGTCT |
| dDocent_Contig_17910_84 | CTCGGTTTTCGCGTTGTTGA | TTGGTGCTGGTCCTGGAATA | CTCGGTTTTCGCGTTGTTGA | Seq | CTCGGTTTTCGCGTTGTTGATTTGAACAATAGCCATGTTGATCTGGTGGTCAGTAGGGTCC[A/C]TTGGTGCTGGTCCTGGAATA |
| dDocent_Contig_18503_242 | GCTTTTAGATGGCAGGCAGC | GGAGCAGTGGCCTAGAGGTA | GCTTTTAGATGGCAGGCAGC | Seq | GCTTTTAGATGGCAGGCAGCATGATGTGGTGCTACATAAATCGTAACCGTATCTC[T/C]GTGGGGAGCAGTGGCCTAGAGGTA |
| dDocent_Contig_18847_119 | AGGCAGAAGAAATCAAAGCAGT | GCCATGTTCAGACAGGGTCC | GGACCCTGTCTGAACATGGC | Seq_reverse_complemented | GGACCCTGTCTGAACATGGCAAC[T/A]ATCAGTGCTATCAATTCATTCTTCTACATAAATGCTCTCAATTTATCTCAGAAATGTATGCAAATCATTAGTGACTGCTTTGATTTCTTCTGCCT |
| dDocent_Contig_19058_194 | TCACAGACACTTTATATACGGTCCA | TCTCTGTAGTAGTTGGTTGGTTGT | TCACAGACACTTTATATACGGTCCA | Seq | TCACAGACACTTTATATACGGTCCATGATGGATA[C/T]ATGTACTTCAAAATTTAGAACAAACCTTTTAGTCCACCTAGTTCTTAAAACTAGGGTCTAGTTCTCTGTAGTAGTTGGTTGGTTGT |
| dDocent_Contig_19231_268 | CTGGCTTGCTTCAAAGGTCAC | TGTGCTAGCGTCAAGCCTG | CAGGCTTGACGCTAGCACA | Seq_reverse_complemented | CAGGCTTGACGCTAGCACA[A/G]RCATGTCCTCCGAAATCCTAACATGTAATTTGCCTATGCTATGAATAAAGCACCTTCATATGTGACCTTTGAAGCAAGCCAG |
| dDocent_Contig_19332_44 | TGCAGAAATGAGAGGTTGCATG | GGACTGACAGACACAGTGACA | TGCAGAAATGAGAGGTTGCATG | Seq | TGCAGAAATGAGAGGTTGCATGTACGATCTTCGACGAATGCG[T/C]CAGGACTGACAGACACAGTGACA |
| dDocent_Contig_19754_248 | TGGAAATCTGGCGTTCAATGC | AGTGTTGAAGTATTTTAGAGACGCA | TGGAAATCTGGCGTTCAATGC | Seq | TGGAAATCTGGCGTTCAATGCATTTACTCTTATCGAGTTTTAATT[C/T]ACGTTAAAGTGTTGAAGTATTTTAGAGACGCA |
| dDocent_Contig_20347_246 | ACACCCTGCATTAGAGTCACA | ACAATGATGCCATCTTTAATGAAGC | ACACCCTGCATTAGAGTCACA | Seq | ACACCCTGCATTAGAGTCACATTTATAATATGAGTTGAGTCACATTTATAATATGAGTTAGAGGA[C/T]CAACAATGATGCCATCTTTAATGAAGC |
| dDocent_Contig_20531_247 | ATGCACCTTTTCGGGTAGCA | TGATTGTCTCCACAGAAACGGT | ATGCACCTTTTCGGGTAGCA | Seq | ATGCACCTTTTCGGGTAGCAGATTGCAAAGCAGGCTTCTAATGACTAATACT[G/A]CTGATTGTCTCCACAGAAACGGT |
| dDocent_Contig_20911_243 | CAGTCAGCCTTTCTGTATCGGT | AGTGTCTCAATAGCACGGCG | CAGTCAGCCTTTCTGTATCGGT | Seq | CAGTCAGCCTTTCTGTATCGGTAATGTGTAT[A/G]AGGAAGCGCGCTACACAAGTGTCTCAATAGCACGGCG |
| dDocent_Contig_20927_242 | ACCCTTACAGTTAAATTGGAGCA | ACGATTAATCAACACAATATCAGCG | ACCCTTACAGTTAAATTGGAGCA | Seq | ACCCTTACAGTTAAATTGGAGCATTTATAAGC[A/C]ATGCATCACAATAGAAATTATATGGTATGACGATTAATCAACACAATATCAGCG |
| dDocent_Contig_21201_191 | AGAGGTGAGGAAGGTACGTGA | GGACACAATGAGAGCTTTGCA | TGCAAAGCTCTCATTGTGTCC | Seq_reverse_complemented | TGCAAAGCTCTCATTGTGTCCT[T/G]TAATCTGATATATCTCTCTTCTATCCTTGCTTTTTCCTNNNNNNNNNNNTACTCGTGTTGTTTGTGTTGTTCTGATTCACGTACCTTCCTCACCTCT |
| dDocent_Contig_21241_201 | ACAATGTACAACAATTAGCCGCA | ATGTGTGAATTTGGGCAATTTAGAA | ACAATGTACAACAATTAGCCGCA | Seq | ACAATGTACAACAATTAGCCGCAAATTATAAGTATCACTGTCATTTAGAATAAG[C/T]TTATGTGTGAATTTGGGCAATTTAGAA |
| dDocent_Contig_21272_238 | ATGTCCGCGACTCTAGTTGG | CGCTTAACGGACGAGCTTTG | ATGTCCGCGACTCTAGTTGG | Seq | ATGTCCGCGACTCTAGTTGGTATATCGGGACAGATTTG[G/A]AAAATAAATCAACCATATCAACTGCGCTTAACGGACGAGCTTTG |
| dDocent_Contig_21523_227 | GAAGTGGTCGAAGGCACAGA | TGTTACCTGTAAGGTTCCAAAAGGA | GAAGTGGTCGAAGGCACAGA | Seq | GAAGTGGTCGAAGGCACAGAAGATTGTAAAATTCGTTAGACCAGTATACAAGCCACTAAA[T/A]TCGATAATGTTACCTGTAAGGTTCCAAAAGGA |
| dDocent_Contig_21596_105 | GCCCTTTGACAGACATGTGC | TTGCTGTGAGTCGGTGTGAG | GCCCTTTGACAGACATGTGC | Seq | GCCCTTTGACAGACATGTGCCCAGAAAGTGTCAGTATTARA[C/T]CCAGCAAACAACCAGATAATATACCTCACTACACAGTCNNNNNNNNNNNATGGTGGTATATTGCTGTGAGTCGGTGTGAG |
| dDocent_Contig_22038_208 | TGTAGGCTAAAGAGTTCTCAAGCT | TCCGACAAACTATTCAACATGTGT | TGTAGGCTAAAGAGTTCTCAAGCT | Seq | TGTAGGCTAAAGAGTTCTCAAGCTAACAATC[G/A]AACATGGTTAGGTCTTCCGACAAACTATTCAACATGTGT |
| dDocent_Contig_22140_158 | TGTGTCAATTCGGTCGACTCA | ACCCAGATGTATATAACCGCTACG | CGTAGCGGTTATATACATCTGGGT | Seq_reverse_complemented | CGTAGCGGTTATATACATCTGGGTTC[T/C]ATAGNNNNNNNNNNNAAAGCCAGAGAAAGAGTGTGGAAGTATCCGTGTATTTTCGTGGTAGGATACCTATGAGTCGACCGAATTGACACA |
| dDocent_Contig_22611_33 | GCAGATCAAAATGTTCATACAATGC | TTGTTAATGGTCGGTAGTCTTTGA | GCAGATCAAAATGTTCATACAATGC | Seq | GCAGATCAAAATGTTCATACAATGCTCTKG[A/G]GGTCTCTTTTAATAGGTTAACTTTCATTGTTAATGGTCGGTAGTCTTTGA |
| dDocent_Contig_22680_236 | TGATCTGCGCATGAATTTGTCT | TGTCTCGATAGCCAAACGGT | TGATCTGCGCATGAATTTGTCT | Seq | TGATCTGCGCATGAATTTGTCTTACTGTATGTTTTGGTCCAGACTGTTTWC[G/T]CAAAAATATGTCTCGATAGCCAAACGGT |
| dDocent_Contig_23105_165 | ACTGGAATCTGTGGAGTTGCT | GCATGCATAGAATCCATCCAACA | TGTTGGATGGATTCTATGCATGC | Seq_reverse_complemented | TGTTGGATGGATTCTATGCATGCATGT[G/A]TACATATCTTCNNNNNNNNNNNATGGAAGATGGATTAATGAACAATGATGTATAATGCGAGAACGTGATTTATTCTAGCAACTCCACAGATTCCAGT |
| dDocent_Contig_23428_154 | TCCTTGTTGATCCAGTTTGTCTT | ACTGCGTTACAGGTCACCTG | TCCTTGTTGATCCAGTTTGTCTT | Seq | TCCTTGTTGATCCAGTTTGTCTTACTGACATACAATACTGAAATTCNNNNNNNNNNN[A/G]AAGTGTATTTATAAAATTGTTATATATGTTTTGTATACAGTCAAAACTGCGTTACAGGTCACCTG |
| dDocent_Contig_23972_178 | TCTTGTGTGACAGAGATCAGACC | TGTGCCAAAATCGCATGTCC | TCTTGTGTGACAGAGATCAGACC | Seq | TCTTGTGTGACAGAGATCAGACCNNNNNNNNNNNCCGTTTGTTACTAGGTCACAGACT[T/G]GGTCTTATGATGTACATTTTACTGATTGTGCCAAAATCGCATGTCC |
| dDocent_Contig_24207_58 | AACACACGGATTCGGTGACA | GCGAGCAAATAGGTGAATGTGA | AACACACGGATTCGGTGACA | Seq | AACACACGGATTCGGTGACATCTAACTTGAACACAATGTATAAATAACCC[T/G]CTACGTTTAAATACAGCAGTTGTAAAAAATGTTGAGTCTTGAGTAAATCTAGCGAGCAAATAGGTGAATGTGA |
| dDocent_Contig_2527_113 | AGGGTACAACATACATGGCCC | GCAAAGACCACCAGCTGTTG | AGGGTACAACATACATGGCCC | Seq | AGGGTACAACATACATGGCCCCTTCTAATATAAGTAAAGTGAACGAAAC[T/C]CCTAAACATTCGAGTGGAAGTCAAAGTAGTNNNNNNNNNNNATATAATGATTCATGAGCAAAGACCACCAGCTGTTG |
| dDocent_Contig_26506_167 | ACATGGAGTCATTCACTGTGAC | GGACACGGGCATTTAAAGCG | ACATGGAGTCATTCACTGTGAC | Seq | ACATGGAGTCATTCACTGTGACGTAGTGCGTATNNNNNNNNNNNCTGACATCAGATA[C/A]AGGACACGGGCATTTAAAGCG |
| dDocent_Contig_26826_189 | ACATGGGCAGTGAAAATCGC | TCGGAAGGTTTCAATACAGGCA | ACATGGGCAGTGAAAATCGC | Seq | ACATGGGCAGTGAAAATCGCTGTGTAAAT[C/T]GCAAACTAATGTTAAACAGATCAGGAATAACTGTCACTTAACCAACTACTACTTTTGTTCGGAAGGTTTCAATACAGGCA |
| dDocent_Contig_27463_92 | TGCAGTACACCGTACAACAGA | TCTTTTGAGTTCGGTTGGTGTG | CACACCAACCGAACTCAAAAGA | Seq_reverse_complemented | CACACCAACCGAACTCAAAAGATAAAATATCGT[T/C]GCTGAACTTACACTCGTATGTAGTCCAGCTTCGCAAAAATGACGATATTACTCAAATTTGCTCGCAATATCTGTTGTACGGTGTACTGCA |
| dDocent_Contig_2788_227 | AGACCTTCTGAACAGTGGCG | AGCCGTCGAAGAAGATTCGT | AGACCTTCTGAACAGTGGCG | Seq | AGACCTTCTGAACAGTGGCGTCATGAGCGATGTTGCCTTGCAAAGAGG[A/G]CTTAAAGAAGCCGTCGAAGAAGATTCGT |
| dDocent_Contig_27920_173 | TGGTTTGTTGCATCGTGCTG | TCCTATGTTGCGCTTGAGCA | TGGTTTGTTGCATCGTGCTG | Seq | TGGTTTGTTGCATCGTGCTGCATGTGGTANNNNNNNNNNNNTACAGGAGTTTATTTAGAC[G/A]GAAGGTCATAGTTGTCTCTTACTGACGCATTAGAAGTCCTATGTTGCGCTTGAGCA |
| dDocent_Contig_28233_173 | TGCCGTGTAGATGACATATTCGA | TGTGCTGAGGACAAAGGAAGA | TCTTCCTTTGTCCTCAGCACA | Seq_reverse_complemented | TCTTCCTTTGTCCTCAGCACATACACTGTGTT[T/A]ATATGGGACAAAATATGGTNNNNNNNNNNNTATGTGGTCGAACCATTTCGAATATGTCATCTACACGGCA |
| dDocent_Contig_32417_56 | AGGACCCGAAATGTAAAACTTGT | ACAACTTTAGGGTGGCACGG | AGGACCCGAAATGTAAAACTTGT | Seq | AGGACCCGAAATGTAAAACTTGTCACAGCAGTACATAA[T/C]TGTAGGACCCGAAATGTAAAAAATGTCATAGCAGTTCATAATTGTAGGACCCGAAATGTACAACTTTAGGGTGGCACGG |
| dDocent_Contig_34362_156 | ACTTATGAAGAGTATGTGACCAGTG | TATCCATCGCCACGAGGGTA | TACCCTCGTGGCGATGGATA | Seq_reverse_complemented | TACCCTCGTGGCGATGGATACCACCAGGAATGAG[T/G]TGNNNNNNNNNNNCACTTGTTTCACAATCGACCTTAAATAGATTAGTATAAACTAAACTATGGTAATTATCACTGGTCACATACTCTTCATAAGT |
| dDocent_Contig_35122_84 | GAAAGTTTTGGGCCTGGTCG | TCAGATGCGGACAACTTTGC | GAAAGTTTTGGGCCTGGTCG | Seq | GAAAGTTTTGGGCCTGGTCGTTAGCCTTAAGAACATAGCAGCAATA[C/T]CAAATTTCAGATGCGGACAACTTTGC |
| dDocent_Contig_36757_24 | GCAGGACGCAATAGACAGGT | AAAGATGACGTAGTGAGTCGAAAA | GCAGGACGCAATAGACAGGT | Seq | GCAGGACGCAATAGACAGGTG[C/T]TCTCTGGAAAAACTTAAAGATGACGTAGTGAGTCGAAAA |
| dDocent_Contig_37208_99 | ACGCTTGGGTTTCTAACAGACA | TCTCATTAGGTGTAGCGGAGAA | TTCTCCGCTACACCTAATGAGA | Seq_reverse_complemented | TTCTCCGCTACACCTAATGAGATAATAACTGAACACG[T/A]TATCTCAGAGCCTTGTTATGAGAATATTCGAACGTCACAACGGTTCCATAATTTGTATAATGTCTGTTAGAAACCCAAGCGT |
| dDocent_Contig_3811_74 | TGTCTCGGGCAAGGTCAAAG | TGCATCGCTTATAACGCAGT | TGTCTCGGGCAAGGTCAAAG | Seq | TGTCTCGGGCAAGGTCAAAGGGTGAACTACCATCATA[G/C]GCACGTACGTTGATATTGCATCGCTTATAACGCAGT |
| dDocent_Contig_3976_80 | TTCTCTGGCCTCGTGTTCAC | AGGCTGAATACTTTTATACGGGCT | AGCCCGTATAAAAGTATTCAGCCT | Seq_reverse_complemented | AGCCCGTATAAAAGTATTCAGCCTTACATTCGATACAGTCTCTGAATT[T/C]GACAGAAAACCCATGTTAGCTTGATATTCATCCTGAAAACTAATCCTTTGTGAACACGAGGCCAGAGAA |
| dDocent_Contig_4097_180 | ACAGATCTTAGGTGGAGCCG | GTGTGTGTCACTCTGTTTATGACA | TGTCATAAACAGAGTGACACACAC | Seq_reverse_complemented | TGTCATAAACAGAGTGACACACACTGCATGTTG[G/A]CTGATCAGAGCATGTCTCTATTCCACCNNNNNNNNNNNTTGTTACCAGGAGTTTCGGCTCCACCTAAGATCTGT |
| dDocent_Contig_4278_228 | TCCATGATAACACTGCGATACGT | TGTCGACAATCATGGTGTAACTG | TCCATGATAACACTGCGATACGT | Seq | TCCATGATAACACTGCGATACGTTCTGTAATTATTCTATAATCGGTACTTTGAG[G/C]CAAAGATCAGGACATTCTATGTCGACAATCATGGTGTAACTG |
| dDocent_Contig_4296_196 | CGTACCAAACGTTACGTCATTGA | AGGTGAGTGTATCTTGGTTCAGA | CGTACCAAACGTTACGTCATTGA | Seq | CGTACCAAACGTTACGTCATTGACGTTTTAGRTATGCGTGGC[T/C]GTTTGCTGTGTCTGTAAATATCCTGTTTACGTTTAGTAGAAAATAGTGAAAAGGTGAGTGTATCTTGGTTCAGA |
| dDocent_Contig_4355_71 | GCAGTTCGCTTGTGTATGAGG | TGAACCAACCACGATTAACACA | GCAGTTCGCTTGTGTATGAGG | Seq | GCAGTTCGCTTGTGTATGAGGATTGTTCAAAATATTTATGTTTAACAACACACTATTTTATATGAATA[T/A]ATATAGAGATAAATACTGAACCAACCACGATTAACACA |
| dDocent_Contig_46792_119 | AGGTCAGATAAACCAACATTCCC | TCCGTTACTTCGACAGCTTAACA | AGGTCAGATAAACCAACATTCCC | Seq | AGGTCAGATAAACCAACATTCCC[C/T]GTATTCCATACTCATCAATCAGTTNNNNNNNNNNNCCGTTACTTCGACAGCTTAACA |
| dDocent_Contig_5391_237 | TCATCAGCCGAGTACAGTGTG | TCCTATTGGAACCTATAGAGGAAGT | TCATCAGCCGAGTACAGTGTG | Seq | TCATCAGCCGAGTACAGTGTG[A/T]GTATATGTGTCTCGAGTTTTTTCCTATTGGAACCTATAGAGGAAGT |
| dDocent_Contig_5443_205 | CGTCACTATCCAGTCTCTCGT | ACCTCAACTGTTTCATGTTGCT | CGTCACTATCCAGTCTCTCGT | Seq | CGTCACTATCCAGTCTCTCGTCTTCTCT[A/T]GCATTCGCCAAATCAGTTACCTCAACTGTTTCATGTTGCT |
| dDocent_Contig_5450_166 | CGTCTGCTGTCCAATCACCT | GAGGTCATCAAGTTTTGCAGGA | TCCTGCAAAACTTGATGACCTC | Seq_reverse_complemented | TCCTGCAAAACTTGATGACCTCTTCA[A/G]CACTGCTTCATTNNNNNNNNNNNCTCACTATGTCCATCTGTCTTTGTTAGTTGAATAGAAACATCAGAAACCACAGGTGATTGGACAGCAGACG |
| dDocent_Contig_5451_204 | ACTTTTGCACAGGTGTGTACAAA | GGCCGAAAAGGGTCACTTTG | ACTTTTGCACAGGTGTGTACAAA | Seq | ACTTTTGCACAGGTGTGTACAAATCATTATTTCA[A/G]TTAGCGGCCGAAAAGGGTCACTTTG |
| dDocent_Contig_5480_80 | TGCAGTCGCACAAAGAGATT | TTGTGGAAAGTACAGATGTGGAAA | TTTCCACATCTGTACTTTCCACAA | Seq_reverse_complemented | TTTCCACATCTGTACTTTCCACAA[A/T]ATTAAGTATGTTGAACTGTTATATGTGTGTGTAATGAATTATAGAAACTCTGTTTTGAAATCTCTTTGTGCGACTGCA |
| dDocent_Contig_5689_32 | TGCAGAGTCACTTCGAATAACAG | CGCCAGTTGGTGGTAGACAT | TGCAGAGTCACTTCGAATAACAG | Seq | TGCAGAGTCACTTCGAATAACAGTTGGTAA[C/T]GATCAGTTCCGAGATCATCCGCCAGTTGGTGGTAGACAT |
| dDocent_Contig_5729_214 | GAGCAAAATCCTATCGCGGAC | CCATGACCACGAAGAACCCA | GAGCAAAATCCTATCGCGGAC | Seq | GAGCAAAATCCTATCGCGGACAAAATAGATGAATGAAATCATATTAACACYATGAAATAA[T/A]TACCATGACCACGAAGAACCCA |
| dDocent_Contig_5746_182 | AGAATACGTCACACAGCCGT | GGCCGTTCCTCTTGAAGATGA | TCATCTTCAAGAGGAACGGCC | Seq_reverse_complemented | TCATCTTCAAGAGGAACGGCC[T/A]GCRGCTGCTCTTCAGCAACCGATTTGTCNNNNNNNNNNNAATTTAGGACCGTAAATACACTATGTATACGGCTGTGTGACGTATTCT |
| dDocent_Contig_5802_131 | CAATTGACGAATGACGAGAACCA | ATTAGCCCAATCAAGCGCCT | CAATTGACGAATGACGAGAACCA | Seq | CAATTGACGAATGACGAGAACCAATATCAATAATGTATATAAGGAATAATAGGG[C/G]ATCTAATATTGANNNNNNNNNNNTGAGAAATGATAAAGATGATTAGCCCAATCAAGCGCCT |
| dDocent_Contig_5841_203 | TGCCCACTTACCGTTTAACCT | AGTTGGACAGACTATGTTCATTGT | ACAATGAACATAGTCTGTCCAACT | Seq_reverse_complemented | ACAATGAACATAGTCTGTCCAACT[T/A]AACAATGTAAGACGTTTTAAGATGACTACATGTATAATGGACCTACATTNNNNNNNNNNNTACATTACAGGATAGGTTAAACGGTAAGTGGGCA |
| dDocent_Contig_5941_141 | GCAGTTGCAATTGCTGTCGT | GATTCCCTCCGAAACCACGA | GCAGTTGCAATTGCTGTCGT | Seq | GCAGTTGCAATTGCTGTCGTAATAATACTGATCAGTGGTTTTTACTGGACAGCCATCGAA[A/C]AANNNNNNNNNNNCTATTTTCACAGGCAGCTACATTTGTAAATCTGTTTGGATTCCCTCCGAAACCACGA |
| dDocent_Contig_5994_87 | CTTAGCAACCACAATAGCGCC | CTACGGGCGCTTGAATGAAA | CTTAGCAACCACAATAGCGCC | Seq | CTTAGCAACCACAATAGCGCCGCGATAATCGTCGTAACGTTAATCTAAGC[G/C]CTACGGGCGCTTGAATGAAA |
| dDocent_Contig_6266_243 | TCACATTTTGCTTTCACCAGACA | ACTGTGTCTATATCACCATGTACCA | TCACATTTTGCTTTCACCAGACA | Seq | TCACATTTTGCTTTCACCAGACAGAAGATAAATATGGTCAACTATACCTTATGTTCCTCCACCA[G/A]GTTGTTGACTGTGTCTATATCACCATGTACCA |
| dDocent_Contig_6289_81 | ACTTCCCTGGGACAAAACTGG | CTCCACTATTGAGTTAGCCCGT | ACTTCCCTGGGACAAAACTGG | Seq | ACTTCCCTGGGACAAAACTGGAACAATTCCCGCCAAACGAYCAATTATAAAGGCAGACATAGTTCC[T/G]TAGTTATTAACTCCACTATTGAGTTAGCCCGT |
| dDocent_Contig_6430_156 | GCCCTTCATAGACCGTGTGT | CGATCTAACTTCGTGCTATGCA | GCCCTTCATAGACCGTGTGT | Seq | GCCCTTCATAGACCGTGTGTAGGCCTCGTTATTCGTCACAGNNNNNNNNNNNTG[G/A]CGGACATCTAACGATCTAACTTCGTGCTATGCA |
| dDocent_Contig_6481_134 | CGACCACAGCATCGTATGGA | CCAAAATCAAGGCAGGAATCTCC | CGACCACAGCATCGTATGGA | Seq | CGACCACAGCATCGTATGGATAGATGGATAGGACAACTGCTCGCT[T/G]TGAGAGTCCNNNNNNNNNNNACTATGAGATGATATACTGGCCAAAATCAAGGCAGGAATCTCC |
| dDocent_Contig_6604_221 | TCGAGACCCTGTGTACACCT | GGCAAAACATCCAACTCTAGGG | TCGAGACCCTGTGTACACCT | Seq | TCGAGACCCTGTGTACACCTTTGCACTGAGAAATAGGAAAGAGCTTGCTATTTCAGCGGAM[T/A]ATTGAAGTCTAAAACACTTCACAGAGAAGAAGAATTGGCAAAACATCCAACTCTAGGG |
| dDocent_Contig_6684_142 | TCCATGTGTGGTTGCTGACA | ACAGTTAATACCCTAAGGCATTCAC | TCCATGTGTGGTTGCTGACA | Seq | TCCATGTGTGGTTGCTGACATCTCATGGTTGATTAAGTGTATCTCATC[C/G]GNNNNNNNNNNNACATTAGGCATGTTGATATGTAATTTGACAGTTAATACCCTAAGGCATTCAC |
| dDocent_Contig_7035_200 | ATAACCATCAAGCGCAGGCG | AGGAAAGTGGCGTGGATGAG | ATAACCATCAAGCGCAGGCG | Seq | ATAACCATCAAGCGCAGGCGT[G/A]TCAGATGTCAAATAAAAATTATATAGAGAATAATAGGAAAGTGGCGTGGATGAG |
| dDocent_Contig_7302_67 | GAGGGACCAACTGTGAGCTG | GGCGCGAAATCGGTGTTTAT | GAGGGACCAACTGTGAGCTG | Seq | GAGGGACCAACTGTGAGCTGATCCACACCAGAACCGAACGATA[T/C]GTTTCCTGGCGCGAAATCGGTGTTTAT |
| dDocent_Contig_7367_36 | TGCAGGCATCATACACGCAT | TGTAGTGCAACATCTGGATGAGA | TGCAGGCATCATACACGCAT | Seq | TGCAGGCATCATACACGCATTGKTGGTRAWAKTG[T/A]TCTACTTGTAGTGCAACATCTGGATGAGA |
| dDocent_Contig_7554_92 | AGCAGCGCACTCTAGACATG | AGCAAACTTAGGACCAAAATGCA | TGCATTTTGGTCCTAAGTTTGCT | Seq_reverse_complemented | TGCATTTTGGTCCTAAGTTTGCTAAAAAGTAATTTACT[G/A]TAATTATATATATTTCTTCCAGATGTACTGATACAACATAGAAGATTCTACCAAGTTTGCATGTCTAGAGTGCGCTGCT |
| dDocent_Contig_7735_142 | CCAATTTCGGCTGCCTACC | TCCAGGGTTTATTTCACTACAAGAG | CCAATTTCGGCTGCCTACC | Seq | CCAATTTCGGCTGCCTACCG[T/A]GNNNNNNNNNNNTCAACTTCGGATAATGAATGGATTATCCAGGGTTTATTTCACTACAAGAG |
| dDocent_Contig_7820_197 | TGTGGCTTCATCAGGGCAAT | TCCTTCTCTTGATACATCTTTGCCA | TGTGGCTTCATCAGGGCAAT | Seq | TGTGGCTTCATCAGGGCAATTCACTGCTT[C/G]ATCTACATTTGATGTTAATGCTTTTTCTTCAGGCATTTTAATATCCTTCTCTTGATACATCTTTGCCA |
| dDocent_Contig_7891_76 | ACAGACTCAAGCTCCTCCGA | CGACCCCTACCAGCTATACC | ACAGACTCAAGCTCCTCCGA | Seq | ACAGACTCAAGCTCCTCCGACCCCTTCAAAAATGTTAAA[T/C]CGACCCCTACCAGCTATACC |
| dDocent_Contig_7916_246 | CCCTGCGTCATACCTGGAAC | TGTATGATGAAATGTTCTGTGCTGT | ACAGCACAGAACATTTCATCATACA | Seq_reverse_complemented | ACAGCACAGAACATTTCATCATACAGTAGTTAACCAAGA[G/A]CAACACCAGTTAAGGCATTCGTCAATATTTCAGTTCTAATAYAACTAAATGTTTTATCCTGGTCGTTCCAGGTATGACGCAGGG |
| dDocent_Contig_8149_31 | TGCAGAAACACTGGGAAACTG | AAGATTTGGCCGCTGATGGA | TGCAGAAACACTGGGAAACTG | Seq | TGCAGAAACACTGGGAAACTGAAAATGAA[A/T]ACTGGTATATTTCATTGATGTGCATTATCATGTGTTCAGGCTGCCTCAAAAGATTTGGCCGCTGATGGA |
| dDocent_Contig_8379_234 | ACTCACGCGGAATAACACCT | TGTTCAACTAAGGACGCAACG | ACTCACGCGGAATAACACCT | Seq | ACTCACGCGGAATAACACCTTGGAAAT[C/A]GGCATCACTGATGTTCAACTAAGGACGCAACG |
| dDocent_Contig_8451_206 | ACCTAATGGCTTCACTGGGC | ACCAGTCTCGTCAGTGTTTCC | ACCTAATGGCTTCACTGGGC | Seq | ACCTAATGGCTTCACTGGGCAATGTTGTAGTRGAGAGYAC[G/A]TCACAGTCTCAGCACATAACCAGTCTCGTCAGTGTTTCC |
| dDocent_Contig_8795_50 | GCAGTGAGTGATAATTAGTGAATGC | TGCAACAAAATGGTTTCCTCCA | GCAGTGAGTGATAATTAGTGAATGC | Seq | GCAGTGAGTGATAATTAGTGAATGCTATAGGATTTATTTCACCGATA[G/C]AACTAAGTATCTCATTAGATAGCATAAAATAAATGTTATGATCAGAAAAAGAGAATGCAACAAAATGGTTTCCTCCA |
| dDocent_Contig_8834_135 | AGGGATAAAAGGGTCAATTAAAGGG | ACAAGGGCAATCAACTGGACA | AGGGATAAAAGGGTCAATTAAAGGG | Seq | AGGGATAAAAGGGTCAATTAAAGGGACAAAT[G/A]AGGACCCCNNNNNNNNNNNTGACATAAATATTGACTACAAGGGCAATCAACTGGACA |
| dDocent_Contig_8899_245 | GTTCGATTGAAAACCAGGTCCA | TTCCTCAGCGTATTCACCCT | AGGGTGAATACGCTGAGGAA | Seq_reverse_complemented | AGGGTGAATACGCTGAGGAATCTCCTTTTAATGATGCTGT[T/A]TACACAGTTAAATTCGAGGATGGATACGACAAATCAATACCCGCGAAAAGTATKATTGGACCTGGTTTTCAATCGAAC |
| dDocent_Contig_8948_104 | CAGGCTGCACTACTGGGAAA | TCTAACCTGTGGTCCACTGG | CCAGTGGACCACAGGTTAGA | Seq_reverse_complemented | CCAGTGGACCACAGGTTAGATTGAA[G/C]ATATTACTTGACTGATCAATATAAATATGATGACTGAATGSTCTCAGGTTTTCATTTCATCATAATATTTTATGAGATAATTTCCCAGTAGTGCAGCCTG |
| dDocent_Contig_8990_231 | TCCCTGTTATCACACTGATGACA | ACACCTCTATCTTGCATTTCACG | TCCCTGTTATCACACTGATGACA | Seq | TCCCTGTTATCACACTGATGACAGCATA[C/A]ATCTTATCTACACCTCTATCTTGCATTTCACG |
| dDocent_Contig_8992_245 | AGTGTAGAATCTAGGGAATGGGT | ACCAGTGTTCTAAATGATGCTTCT | AGTGTAGAATCTAGGGAATGGGT | Seq | AGTGTAGAATCTAGGGAATGGGTATATTATATCACGGTATTAAACAATGGTAAG[G/A]CGTACATTACCTATAACCAGTGTTCTAAATGATGCTTCT |
| dDocent_Contig_9531_217 | CTGGATCCTGTGCTTGAGGT | GTGCTCTCGGACTCTGATGA | CTGGATCCTGTGCTTGAGGT | Seq | CTGGATCCTGTGCTTGAGGTATCAGAGAGTGACTCGTCACTACTGCTCATTCCAGA[T/C]GTGCTCTCGGACTCTGATGA |
| dDocent_Contig_9558_204 | GTTGCTGATTTCATGGTGGTGA | ATGAGGTGCTGTGCCTACAC | GTTGCTGATTTCATGGTGGTGA | Seq | GTTGCTGATTTCATGGTGGTGATTTGGTCATGTACTTCTGTTTCATCATC[A/G]CTATATTGAGGAATGAGGTGCTGTGCCTACAC |
| dDocent_Contig_9635_159 | GCCATCTTGTCGTTGTGCAT | TGTGTTGACGGGTTTTATGTTGG | CCAACATAAAACCCGTCAACACA | Seq_reverse_complemented | CCAACATAAAACCCGTCAACACATAAATATCCAAATCTCATC[C/T]AAAGANNNNNNNNNNNTTTAATAATACACGGATAAATTTCAAAACCGTTCGCGGTCCGTATGCACAACGACAAGATGGC |
| dDocent_Contig_9701_202 | CGGCCAGTAAACCAGTCCAA | ACCAGTCGTATAATTATGCTGAGCT | CGGCCAGTAAACCAGTCCAA | Seq | CGGCCAGTAAACCAGTCCAAATGAAGACTACTCAGACTAATTATTAT[A/T]ACACTGACAACCAGTCGTATAATTATGCTGAGCT |
| dDocent_Contig_9782_101 | TCAGTTGCTGTTTGAAATTCTGTCA | TGTTGCCCGCCTTATTTGGTG | TCAGTTGCTGTTTGAAATTCTGTCA | Seq | TCAGTTGCTGTTTGAAATTCTGTCAATGAAGATAGTTTTTTGT[A/T]ATGTTGCCCGCCTTATTTGGTG |
| dDocent_Contig_9837_70 | GTTTTCAGCGTCAGAAAGCGT | TCTGTCTTGCTCCAGGTACC | GTTTTCAGCGTCAGAAAGCGT | Seq | GTTTTCAGCGTCAGAAAGCGTATCTTCTTCAAACATGTCTGCTGTAAGTCTGTGAGAATGT[G/A]GCTCCTGACTGTCCCATCTTTGACAAACCTTTCCTTTTTCTGTCTTGCTCCAGGTACC |
| dDocent_Contig_9930_117 | CCACCACTTGTAGCCAAAGC | GGCTAATTGTTCTTCTGTGGCA | CCACCACTTGTAGCCAAAGC | Seq | CCACCACTTGTAGCCAAAGCTCATCTAACTATACTAGGTAAGTG[G/T]GAATTATTTAACCATGTATCTGCTGGNNNNNNNNNNNGTTCCCTACTTTAACAGGCTAATTGTTCTTCTGTGGCA |

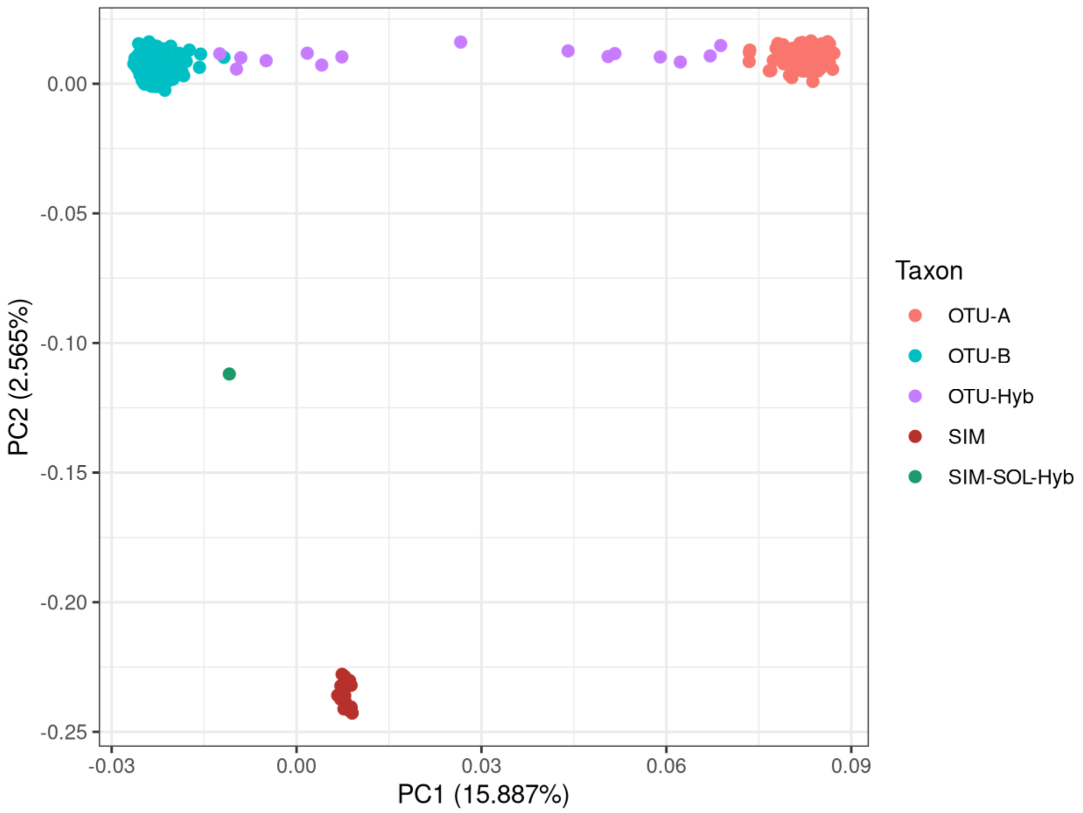


Figure S.1: Principal component plot based on LD-pruned SNPs including all *S.s. similis* (SIM) and *S.s. solidissima* (OTU-A, OTU-B, SOL) samples in this study. The red SIM cluster includes reference *S.s. similis* individuals as well as sampled unknowns. *S.s. solidissima* OTU AxB hybrid (Hyb) individuals are classified as shown in Table 1. The green SIM-SOL-Hyb has admixture proportions consistent with an F1 hybrid.

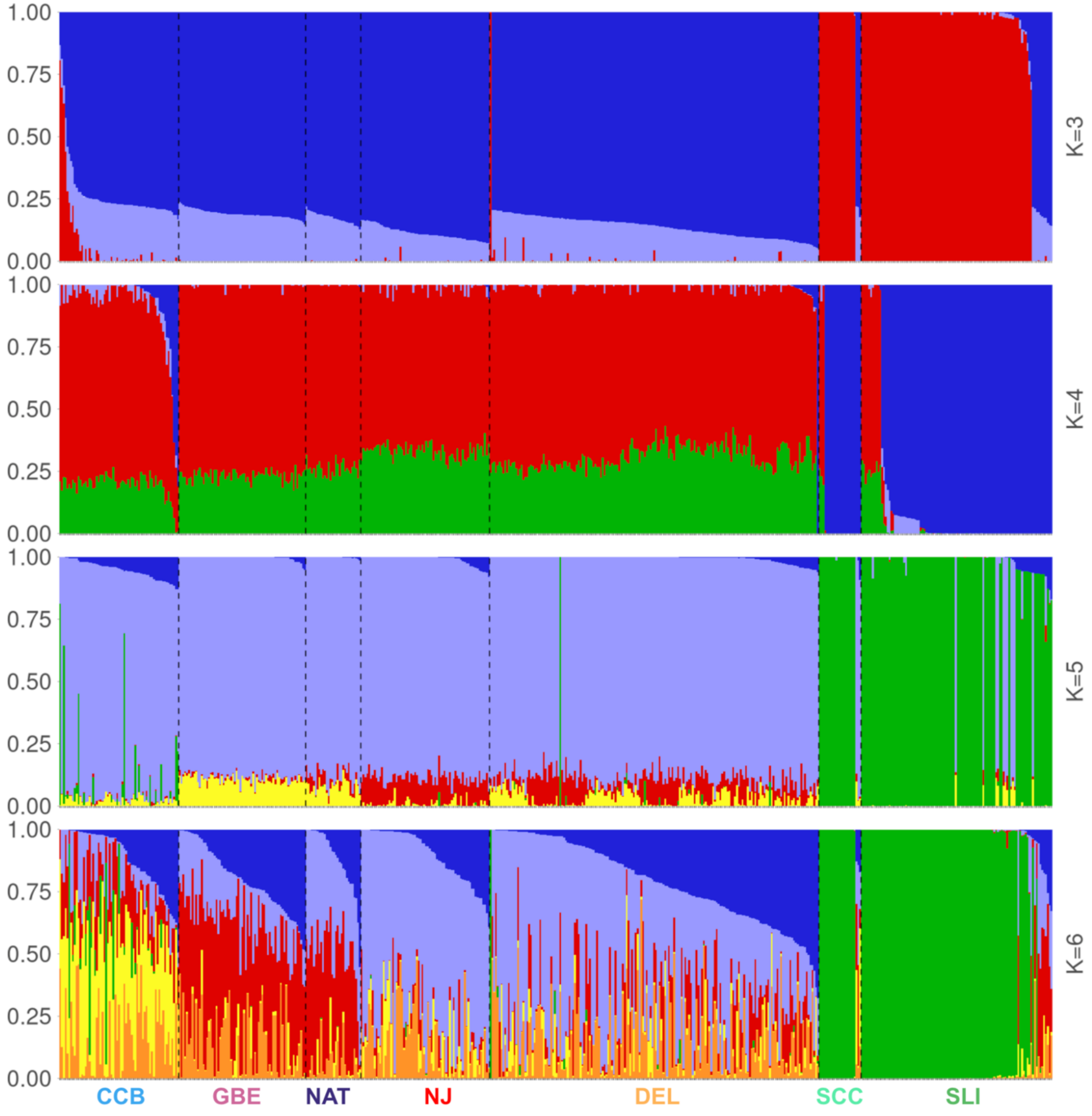


**Figure S.2:** STRUCTURE Clustering of *S.s. solidissima* for K = 3, 4, 5, 6. Each individual is represented with a vertical bar that shows the percentage of admixture attributable to genetic diversity from particular ancestral sources (colors). Analytical region acronyms are shown along the bottom, as defined in Table 1.

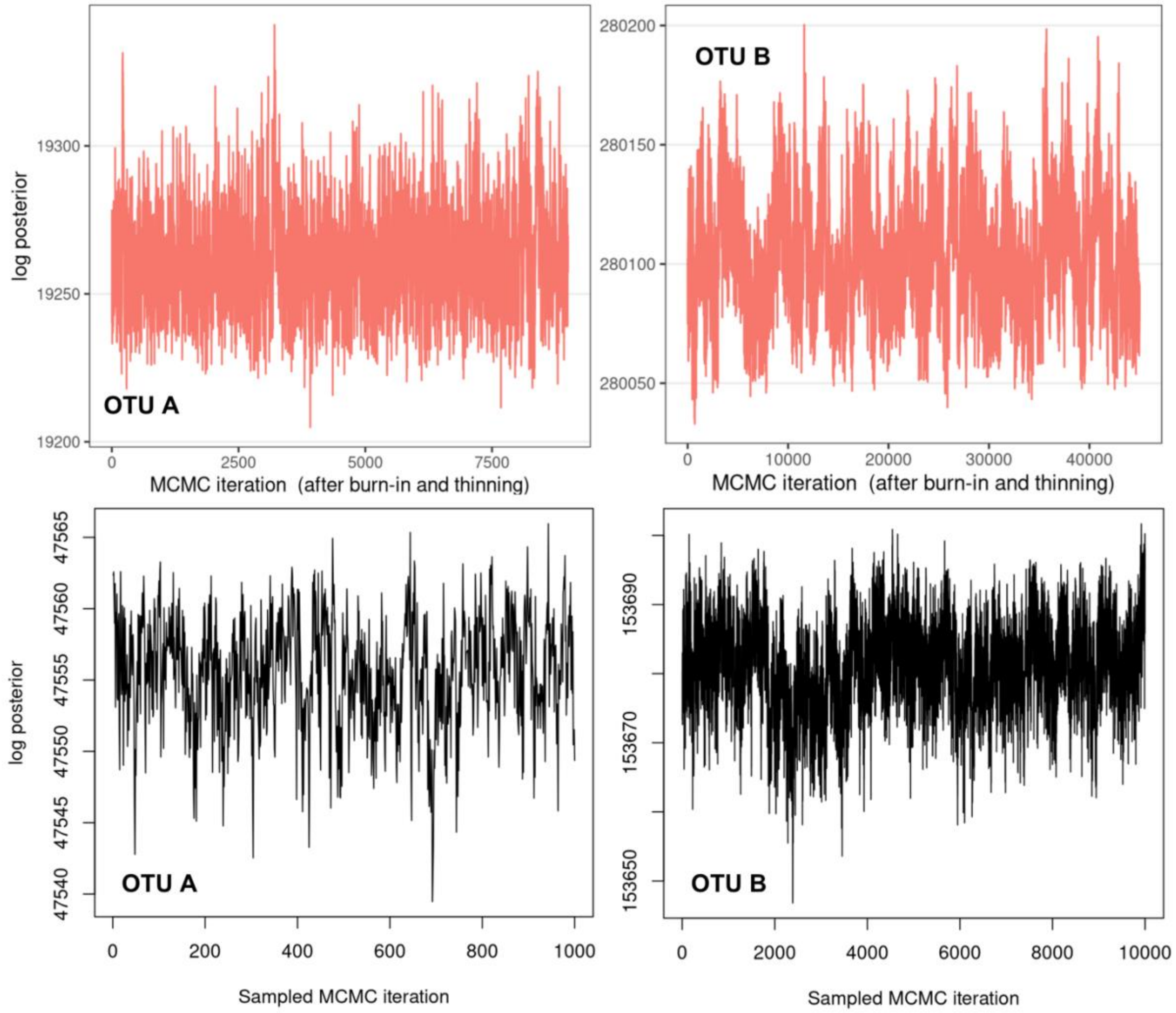


**Figure S.3**: Log posterior plots for `EEMS` and `SpaceMix` MCMC runs to assess convergence. Top row shows `EEMS` plots for the separate analysis of OTU-A (left) and OTU-B (right) individuals. The MCMC was iterated over a total of 10,000,000 iterations, recording every 1000th result. A burnin of 1,000,000 iterations were discarded at the start. Bottom row shows SpaceMix plots for OTU-A (left; 1000 draws from 1 M iterations) and OTU-B (right; 10,000 draws from 10 M iterations).

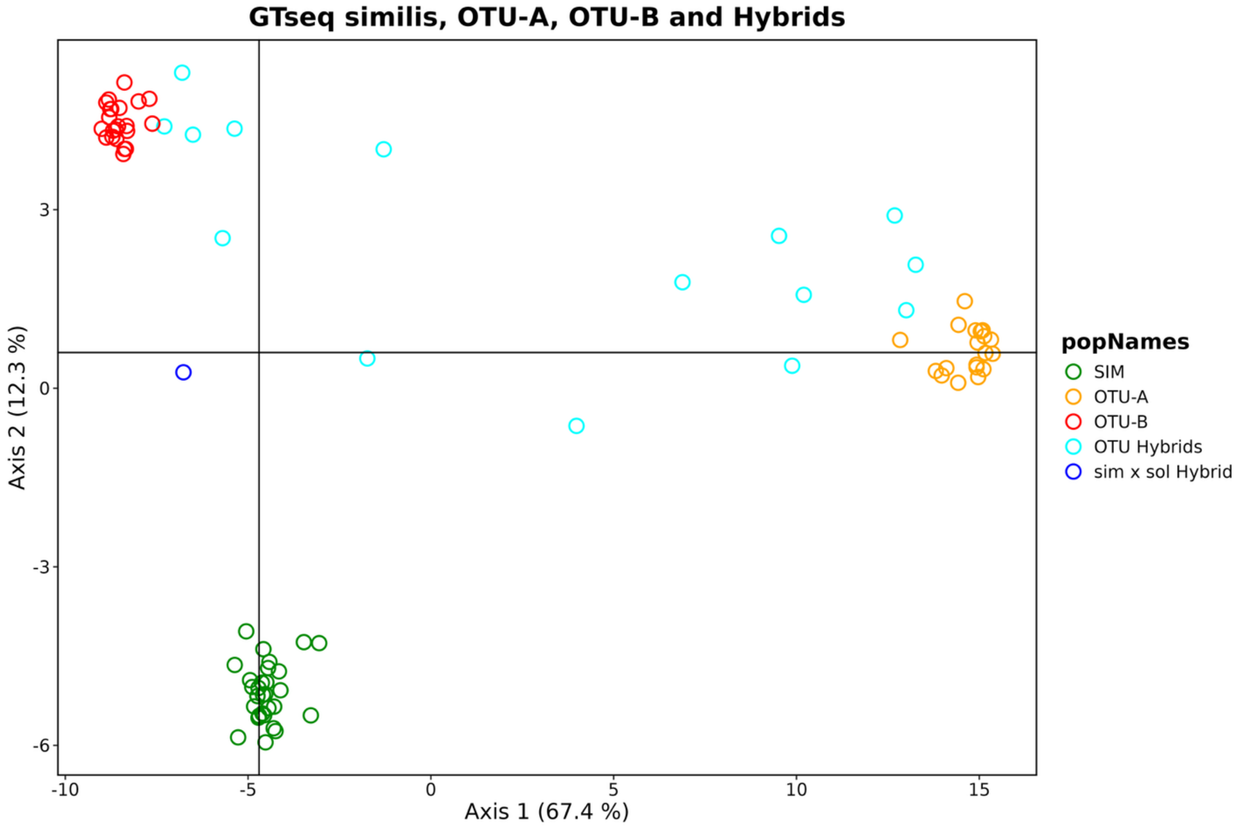


Fig. S.4: Principal component plot of surfclam taxa based on the GTseq panel of 113 informative SNPs. Each open circle is a genotyped individual. "SIM" is *Spisula solidissima similis* and "sim x sol Hybrid" is a putative hybrid between *S.s. similis* and *S.s. solidissima*.

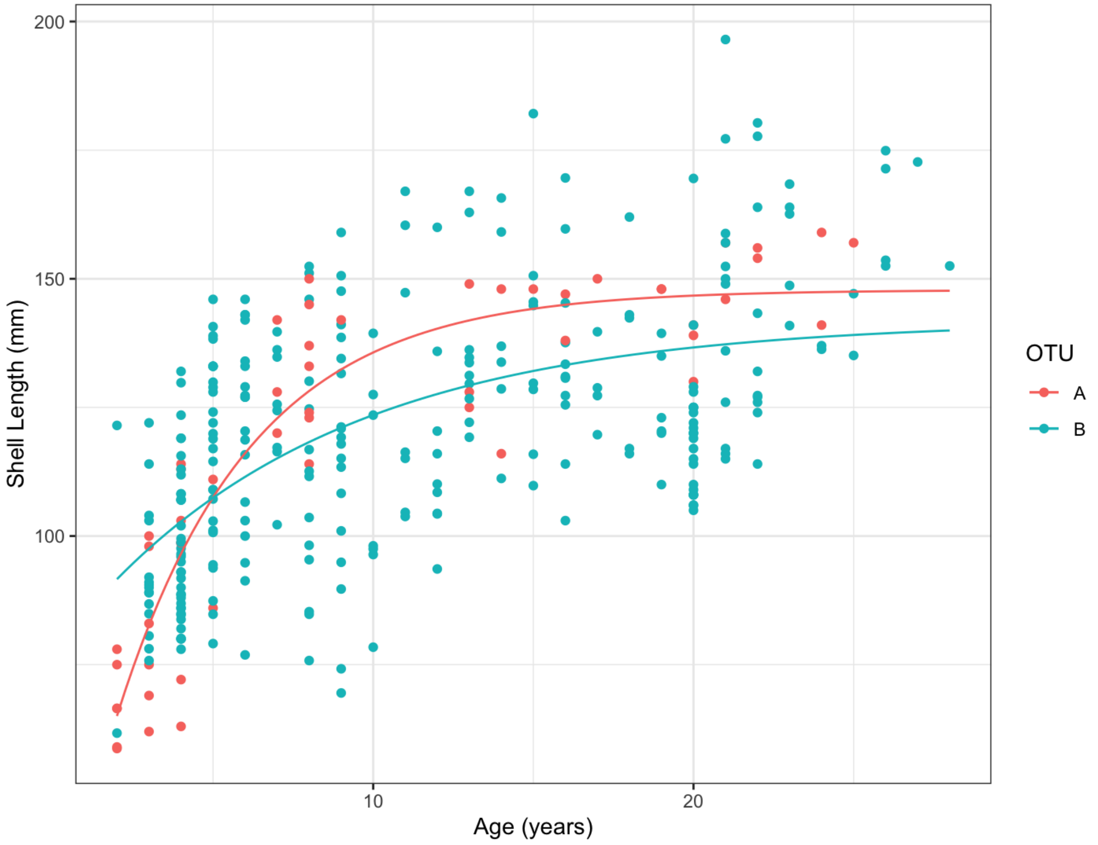


**Figure S.5:** Estimates of Von Bertalanffy growth curves from age and size data for the subset of OTU-A and OTU-B individuals for which age data was available (NY State Department of Environmental Conservation) or obtained for this study (NEFSC).
